# Modeling and Inferring Large-scale Demographic Fluctuations in Structured Populations Through Simulations and PSMC-based Methods

**DOI:** 10.64898/2026.06.17.732814

**Authors:** Camille Steux, Ravi Vishwakarma, Gabriele Sgarlata, Olivier Mazet, Rémi Tournebize, Christophe Thébaud, Benoit Goossens, Lounès Chikhi

## Abstract

The climatic oscillations of the Quaternary have likely affected the demographic history of many species, and PSMC (Pairwise Sequentially Markovian Coalescent) has been widely used to investigate these histories. However, it is increasingly acknowledged that PSMC trajectories are difficult to interpret. First, they are influenced by connectivity changes, even without population size changes. Second, most PSMC curves exhibit a few humps when tens of cycles occurred during the Pleistocene. Finally, responses to ancient habitat change have been shown to be species-specific. To address these issues, we simulated structured populations where connectivity (or population size and connectivity) varied according to successive interglacial and glacial periods during the last 2.6 million years. We computed the IICR (Inverse Instantaneous Coalescence Rate), the function that PSMC estimates, and ran PSMC. We further varied the generation length and assumed that some species were positively or negatively affected by glacials. We found that the IICR carries information regarding the demographic oscillations, but that PSMC fails to recover it for times older than 300 ky. For the last 200 ky, PSMC was often able to reproduce qualitatively the demographic oscillations. We also tested SNIF (Structured Non-stationary Inferential Framework), which produced good results using the IICR curve as an input but not when using the PSMC curve. Altogether, our study suggests that the humps older than 300 ky in PSMC histories are unlikely to represent trends of population size or connectivity. However, improving the estimation of the IICR could potentially help reconstruct some of these past demographic changes.

## Introduction

Reconstructing the demographic history of extant and extinct species has been a central goal in evolutionary biology, from fossil-based approaches to the use of a few genetic markers, to the current availability of genome-wide data. Recent advances in genomic sequencing have been coupled with the development of numerous inference methods designed to leverage the high amount of information generated by these new techniques, in the hope to provide a deeper understanding of historical evolutionary processes (Li and Durbin, 2011; Schiffels and Durbin, 2014; Jouganous et al., 2017; Liu and Fu, 2020). Uncovering species’ demographic history can help shed light on biogeographic events (Salces-Castellano et al., 2021), dispersal and colonization patterns (Li and Durbin, 2011; Martin et al., 2023), selection (Uhl et al., 2025), divergence and speciation (Schiffels and Durbin, 2014), ultimately explaining the processes behind observed patterns of diversity. Furthermore, reconstructing demographic trajectories can improve our understanding of species’ response to past and present environmental and anthropological pressures (Lorenzen et al., 2011; Bergman et al., 2023).

The climatic oscillations of the Quaternary provide a good opportunity to test these demographic inference methods. Throughout the last 2.6 millions years (My), Earth’s climate has undergone several oscillations defined by interglacial periods (such as today) and glacial periods (Williams et al., 1993; Lisiecki and Raymo, 2005; von der Heydt et al., 2021; Herbert, 2023). Glacial periods were associated with a progression of the ice sheets at high latitudes, resulting in a global decrease of sea level and the emergence of land areas previously underwater (Williams et al., 1993; Hewitt, 2000). At low latitudes, climate became generally drier and tropical forests shrank in favor of deserts and savannas, hence increasing the extent of open habitats and decreasing the connectivity between forested habitats (Webb III and Bartlein, 1992; Williams et al., 1993; Comes and Kadereit, 1998). In addition to regional conditions, species’ response to glacial periods was dependent on the habitat that species were adapted to, making glacial periods either favourable or unfavourable depending on the focus species (Fedorov et al., 2020; Sommer and Zachos, 2009). During unfavourable periods, species that did not go extinct may have persisted through plastic response (Valladares et al., 2014), adaptation (Thurman et al., 2020) and/or through a change in their distribution range, either by successfully tracking their habitat through recolonization (Eldredge, 1989; Fedorov et al., 2020) or undergoing local extinctions (Dalén et al., 2007). In all cases, persisting species are expected to have occupied small and isolated refuge areas when conditions were generally unsuitable, and wider and more continuous ranges when conditions were suitable (Hewitt, 2000, 1996; Wüster et al., 2005; Dalén et al., 2007; Sommer and Zachos, 2009; Fedorov et al., 2020). These successive distribution changes are thought to be major drivers of intraspecific diversity and interpopulation divergence, and have been proposed to explain the existence of population structure in various species such as mammals (Brown et al., 2007; Taylor et al., 2021; Cancellare et al., 2024), insects (Sosa-Pivatto et al., 2020), amphibians (Wielstra et al., 2021), or birds (Nadachowska-Brzyska et al., 2015; Wogan et al., 2024).

Understanding the dynamics of genetic diversity under scenarios of connectivity and fragmentation with varying population size is critical to comprehend the evolutionary processes that have shaped current patterns of biodiversity. A small number of studies have aimed to address this issue (Jesus et al., 2006; Arenas et al., 2012; Alcala and Vuilleumier, 2014; Vishwakarma et al., 2025). For instance, under a model with successive phases of panmictic (interpreted as instantaneous expansion) and structured (meta-)population, Jesus et al. (2006) have shown that the expected number of segregating sites in present-day populations is higher than what is expected under a scenario where the population stays panmictic at all time, even if the deme size within the meta-population is small (Jesus et al., 2006). In another work, Alcala and Vuilleumier (2014) derived and analyzed the dynamics of within and between demes heterozygosity in a *n*-island model (Wright, 1931) under successive periods of isolation and connectivity with constant deme size. They showed that cycles of similar length can lead to either an accumulation or a turnover of genetic diversity depending on some species traits (Alcala and Vuilleumier, 2014). Finally, Vishwakarma et al. (2025) have recently shown through spatial simulations of a single glacial cycle that the dynamics of genetic diversity can keep increasing in a refuge population for thousands of years after the start of the habitat contraction, depending on some species-specific traits such as dispersal rate, generation time or local habitat carrying capacity. They also showed that the temporal trajectories of habitat size, census size and inferred effective population size could be highly disconnected, hence questioning our ability to infer past population size changes.

These studies show that the genetic diversity dynamics under glacial cycles is not trivial to predict, questioning the ability of current inferential methods to properly infer the demographic history of species from genetic and genomic data. The Pairwise Sequentially Markovian Coalescent (PSMC) method is now a classic and widely used approach in both model and non-model (possibly endangered) species. Using whole-genome sequence data from a single diploid individual, PSMC estimates the distribution of coalescent times for two haploid samples (*i.e. T*_2_) by walking along the diploid genome, identifying segments delimited by recombination points and inferring the *T*_2_ of each of these segments. The inferred distribution of *T*_2_ is then translated into a trajectory of effective population size, *N_e_*, under the assumption of panmixia and given a fixed mutation rate (Li and Durbin, 2011). Since its publication, the PSMC method has become a routine analysis in genomic studies (Prado-Martinez et al., 2013; Nadachowska-Brzyska et al., 2015; Kuderna et al., 2023; Bursell et al., 2022). Importantly, it has been used to understand species response to repeated past climatic shifts (Nadachowska-Brzyska et al., 2016; Kozma et al., 2016; Ruane et al., 2015; Prates et al., 2016; Bai et al., 2018; Sarabia et al., 2021; Bergman et al., 2023; Sozzoni et al., 2025). For instance, Bergman et al. (2023), analyzed the PSMC curves of 139 megafauna species to determine the role of climate fluctuations and human expansion on megafauna population size dynamics over the last million years.

Such endeavor makes at least two implicit assumptions. First, it assumes that PSMC is inferring effective population size (*N_e_*) changes. Second, it assumes that PSMC is actually able to infer repeated changes in effective population size, such as those caused by the ancient climate cycles that occurred during the Pleistocene. It has now become increasingly clear that the PSMC method infers the Inverse Instantaneous Coalescent Rate, or IICR (see below), and not *N_e_* (Mazet et al., 2016). As for the second point, and to our knowledge, PSMC has not been tested thoroughly under such complex scenarios of repeated cycles of connectivity and population size change in structured populations, or even under panmixia. Furthermore, due to the exponential distribution of coalescent times, the resolution of PSMC inference decreases towards the past, likely limiting the power of the method to detect old and frequent demographic changes. Moreover, it is surprising to note that most PSMC curves, no matter the species, often harbor one, two, or at most three “humps” over the last few millions years, while we know that climate, and also demography, have varied much more frequently across such a long period of time.

Several studies have also employed PSMC to investigate the demographic response of co-occurring or closely related species, to determine if species that rely on the same habitat share similar demographic trends in response to past climatic and/or Anthropocene changes (Ruane et al., 2015; Burbrink et al., 2016; Chattopadhyay et al., 2019; Sozzoni et al., 2025). Interestingly, a lot of these studies have found discordant patterns (Ruane et al., 2015; Burbrink et al., 2016; Germain et al., 2023; Sozzoni et al., 2025) (though not always, see Chattopadhyay et al. (2019)). It then suggests that the demographic response of populations to external factors would be primarily driven by species-specific traits, as proposed by both empirical (Papadopoulou and Knowles, 2016; Germain et al., 2023) and simulation-based works (Arenas et al., 2012; Vishwakarma et al., 2025). It further suggests that comparative demographic studies based on genetic data alone are limited to understand past environmental fluctuations (Hoban et al., 2019), as different species going through similar habitat changes would not be expected to show similar past effective population size histories (Vishwakarma et al., 2025).

As noted above, it has been shown that PSMC infers a complex statistic known as the IICR (Mazet et al., 2016). The IICR is a function of demographic parameters such as deme size, population structure topology and connectivity (as well as the sampling scheme) when such structure exists (Mazet et al., 2016). While in panmictic populations the IICR directly corresponds to *N_e_*, its interpretation under population structure is notoriously challenging (Mazet et al., 2016; Chikhi et al., 2018; Rodríguez et al., 2018; Arredondo et al., 2021; Jouniaux et al., 2026; Chikhi et al., 2026). Although PSMC curves are typically interpreted as changes in population size, they should probably be considered as a summary statistic of historical genome-wide diversity whenever the analyzed species are suspected to be structured. In this context, the program SNIF (for Structured Non-stationary Inferential Framework) has been developed to fit any PSMC curve under a piecewise stationary *n*-island model with varying migration rates and constant deme size (Arredondo et al., 2021; Teixeira et al., 2021; Steux et al., 2025). Using PSMC curves as an estimate of the IICR to infer demographic changes, however, requires that PSMC is able indeed to estimate the IICR reliably. However, it has been shown that PSMC can be biased, depending on the input genomic data quality and mapping (Nadachowska-Brzyska et al., 2016; Mather et al., 2020; Akopyan et al., 2025), PSMC parameters values (Hilgers et al., 2025), or the underlying population model (Nieto et al., 2025).

In this work, we first tested the ability of the IICR and PSMC-based methods (namely PSMC itself (Li and Durbin, 2011) and SNIF (Arredondo et al., 2021)) to infer repeated demographic changes under a structured population model. We simulated genomes under *n*-islands, where either (i) connectivity or (ii) connectivity *and* size oscillated according to a simplified sequence of glacial and interglacial periods (depicted in Figure 1). We computed the IICR and run PSMC and SNIF on the simulated data, and investigated the effect of cyclic demographic changes on the theoretical and inferred objects. To account for species adapted to different habitat and test if differential response to glacial periods have an effect on PSMC curves, we simulated two different responses to glacial periods: (i) connectivity or connectivity and size jointly decrease during glacials, or (ii) they both increase. Finally, we used generation times as a proxy for modeling (*in silico*) species of various types and compared PSMC trajectories obtained under the same demographic scenario but different generation times.

**Figure 1.**
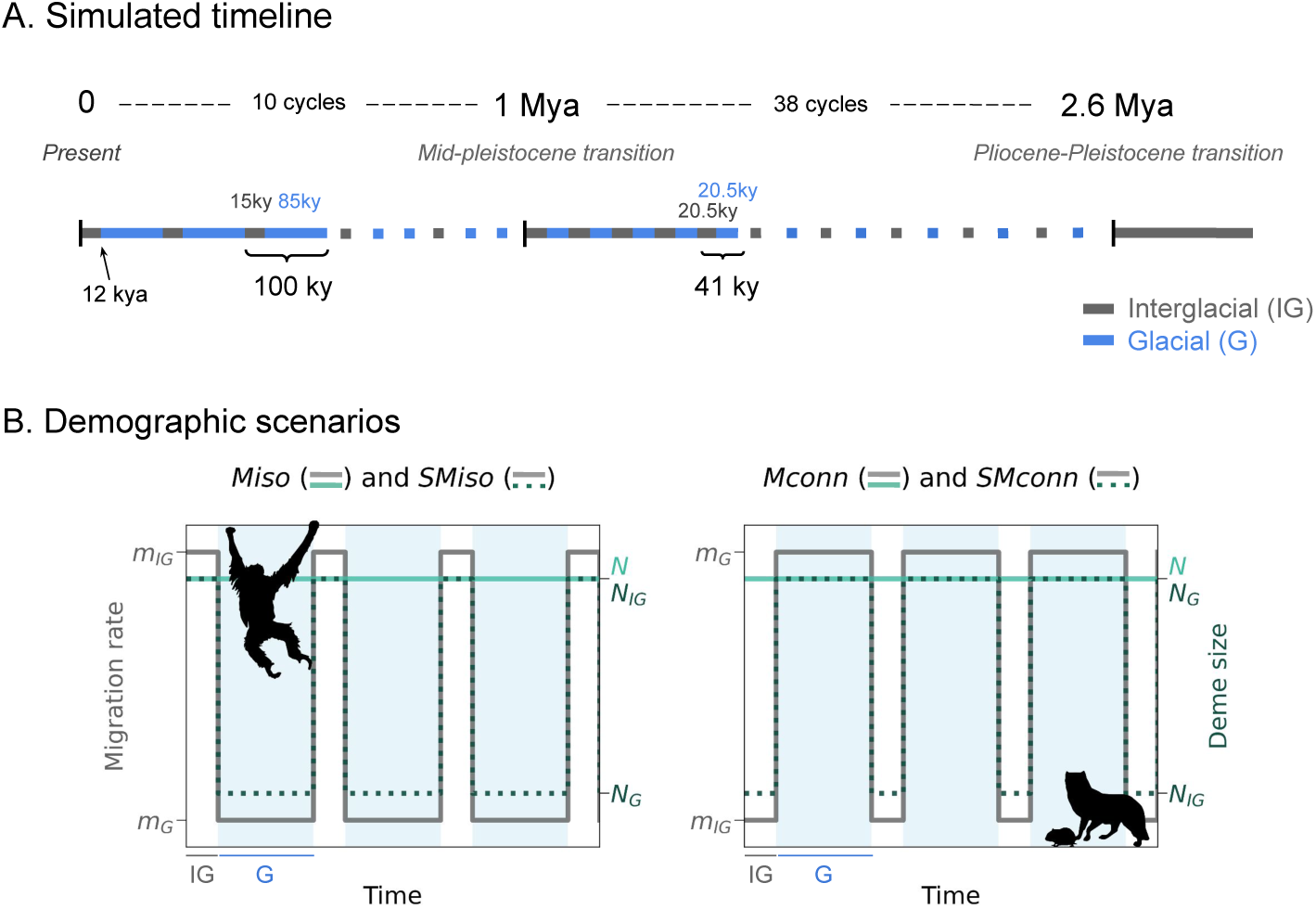
Graphic representation of the simulated timeline and demographic scenarios. Panel A represents the timeline of the simulated glacial and interglacial cycles, in blue and grey, respectively. Panel B represents the simulated demographic scenarios. Left panel shows scenarios with lower migration rates (*Miso*) or lower migration rates and deme sizes (*SMiso*) during glacial periods, which are expected for species dependant on forest habitats such as orangutans (Arora et al., 2010). Right panel shows scenarios with higher migration rates (*Mconn*) or higher migration rates and deme sizes (*SMconn*) during glacial periods, which are expected for species dependant on open and cold habitats such as arctic foxes and collared lemmings (Fedorov et al., 2020; Dalén et al., 2007). Here, interglacial periods (IG) are shorter than glacial periods (G), a situation that corresponds to our timeline between the present and 1 Mya. Animal silhouettes were retrieved from PhyloPic and belong to the public domain.

## Results

### Theoretical IICR

The IICR is the theoretical object that PSMC aims to infer (Mazet et al., 2016; Li and Durbin, 2011), and can therefore be considered the expectation of PSMC inferences if PSMC was a perfect estimator and if molecular data perfectly represented coalescence times. Exact IICR curves for scenarios with connectivity oscillating between “high” (*m_IG_* = 2.5.10^−4^, *M_IG_* = 1) and “low” (*m_G_* = 2.5.10^−5^, *M_G_* = 0.1) migration rates and constant deme size (*Miso*) are presented in Figure 2 for g = 1 and 25 year(s), while results for the other scenarios and parameter values are shown in Supplementary Figure S1-S6, respectively.

**Figure 2.**
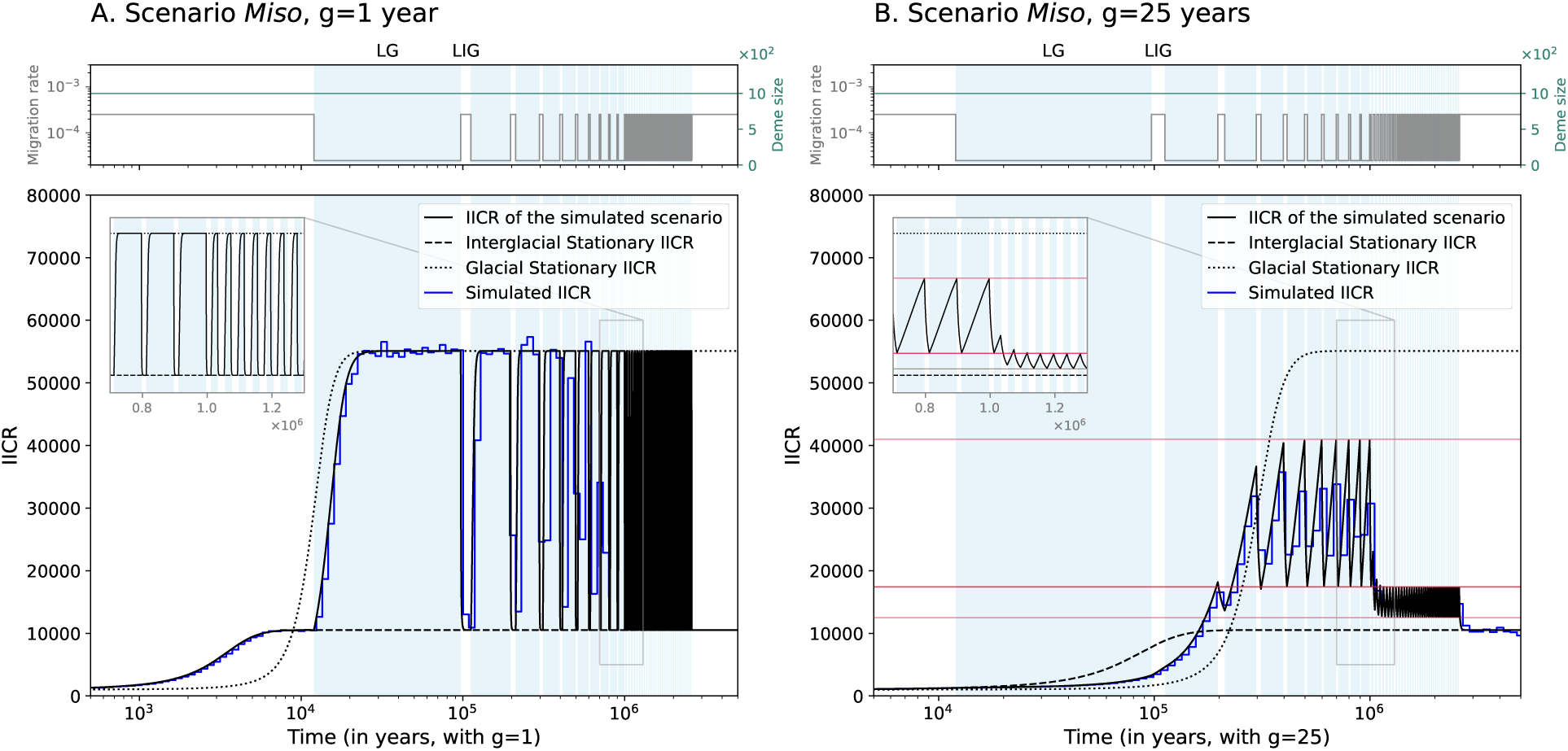
Theoretical IICR of scenarios with lower migration rate during glacial periods and constant deme size (*Miso*). A: Scenario *Miso* for a generation time of 1 year. B: Scenario *Miso* for a generation time of 25 years. Each top panel shows migration rate and deme size (here constant) over time, with grey and turquoise lines, respectively. In each bottom panel, the black continuous curve corresponds to the continuous theoretical IICR of the demographic scenario, and the dotted and dashed curves represent the stationary IICR of the glacial and interglacial periods, respectively. The inserts represent the same objects on a linear-time scale, zoomed-in between 700 kya and 1.3 Mya. The blue curves show the simulated IICR. The red horizontal lines in panel B represent what we refer to as “pseudo-plateaus” in the text. Finally, light blue rectangles in the background of each panel correspond to glacial periods. In this figure, migration oscillates between *m_IG_* = 2.5.10^−4^ and *m_G_* = 2.5.10^−5^, deme size is constant and equal to *N* = 1000 diploids and the number of demes *n* is constant and equal to *n* = 6. LG and LIG stand for Last glacial and Last interglacial, respectively. Note that the x-axis is not the same for the sub-figures for better readability.

When only migration fluctuated (Fig. 2 and Supp. Fig. S1-S3), the theoretical IICR oscillated continuously following the interglacial and glacial period transitions. On Figure 2, for a generation time of one year (2A), the theoretical IICR reached the plateau corresponding to the stationary IICR of the respective migration rate of each period (dashed and dotted curves for the interglacial and glacial periods, reaching IICR values of approximately 10,000 and 55,000, respectively). For a generation time of 25 years (2B), the theoretical IICR did not show the most recent changes in migration rates (< 200 kya) but detected the older fluctuations. Unlike for a generation time of one year though, the theoretical IICR did not reach the corresponding stationary plateaus at each cycle period, but it oscillated between two apparently stable values that we identify as “pseudo-plateaus” (represented with horizontal red lines on the figure on Fig. 2B). Because cycle length in units of generations is reduced for a species with a longer generation time, the glacial and interglacial periods were not long enough for the theoretical IICR to reach the expected stationary IICR curves. Therefore, the “pseudo-plateaus” should depend on both the length of each paleoclimatic period and the species generation length. This is what we saw from simulated data: when the cycle length increased, pseudo-plateaus shifted and the amplitude of the oscillations increased (Supp. Fig. S7, see also Supp. Fig. S1-S3).

These observations also hold for the scenarios with oscillating migration rates and deme sizes (scenarios *SM*) (Supp. Fig. S4-S6). Moreover, unlike the previous results, we observed that changes in deme size (between cycle periods) generated sudden and temporary increases or decreases in the non-stationary IICR, way beyond expectation given by the equivalent stationary IICR. Note that these sudden changes were expected under *n*-island models when deme size changes (based on previous unpublished data by Simon Boitard, Lounès Chikhi and Olivier Mazet).

### Simulated IICR (estimated with simulated *T*_2_)

The theoretical IICR can be estimated from a finite number of *T*_2_ values under a model of interest (here called “simulated IICR”), as noted in the Methods section. This can be seen as the best that PSMC could give if it inferred the exact distribution of *T*_2_ from the distribution of heterozygous sites in the diploid genome. Using 10^6^ values of independent *T*_2_, we observed that the simulated IICR curves show the demographic oscillations and mostly overlap the theoretical IICR curves (Fig. 2, Supp. Fig. S8-S13), in particular for the time period up to 1 Mya. In the older past, the simulated IICR did not capture the oscillations, likely due to the decreasing number of coalescent events. When oscillations were not captured, the simulated IICR curves seemed to approximate the harmonic IICR mean, represented by the red curves on the Supplementary Figures (see for instance Supp. Fig. S8 for g = 5, 10 and 25 years with IICR value of 14,000 between one and 2.5 Mya). We also noted some noise in the most ancient part of the simulated IICR, certainly due again to the limited number of coalescent events in the deep past. Finally, we further observed that the transitory phases of very high or low IICR observed for the scenarios with oscillating migration and deme size (*SMiso* and *SMconn*) almost disappeared when considering the simulated IICR in contrast with the theoretical one (Fig. S11-S13).

### Estimating the IICR with PSMC

When we ran PSMC on genomes simulated under our cyclic scenarios, we found that PSMC generally allowed to recover the oscillations up to 200 kya (Fig. 3A) and sometimes 300 kya (Fig. 3C), and we found little variance in the bootstraps. For scenario *SMiso* and a generation time of one year (Fig. 3A-B), PSMC curves showed some fluctuations seemingly corresponding to the LG, LIG and the glacial period between 100 and 200 kya, albeit with significant departure from the expected oscillations amplitude (see below). For a 25-year generation time (Fig. 3C,D), PSMC curves also exhibited some oscillations up to 200 kya, and even up to 300 kya for scenario *SMiso*, though it that last case PSMC curves did not detect the most recent demographic change 12 kya (as the theoretical IICR, Fig. 3C). We note however that the time at which the PSMC trajectories changed rarely corresponded to the timing at which the actual demographic changes occurred. For instance, on Figure 3A, the PSMC curves increased forward in time (from right to left) from 100 kya to 40 kya, and then decreased up to the end of the LG 12 kya. If we were to interpret PSMC trajectories at face-value, we would consequently suggest a demographic change in the middle of the LG that actually did not occur. Finally, we observed that very few of the PSMC curves of our tested scenarios detected the demographic cycles older than 200 to 300 kya, except for Figure 3C, where PSMC curves do exhibit some fluctuations for times older than 300 kya. However, these fluctuations appear disconnected from the simulated demographic cycles (in terms of timing, amplitudes and duration).

**Figure 3.**
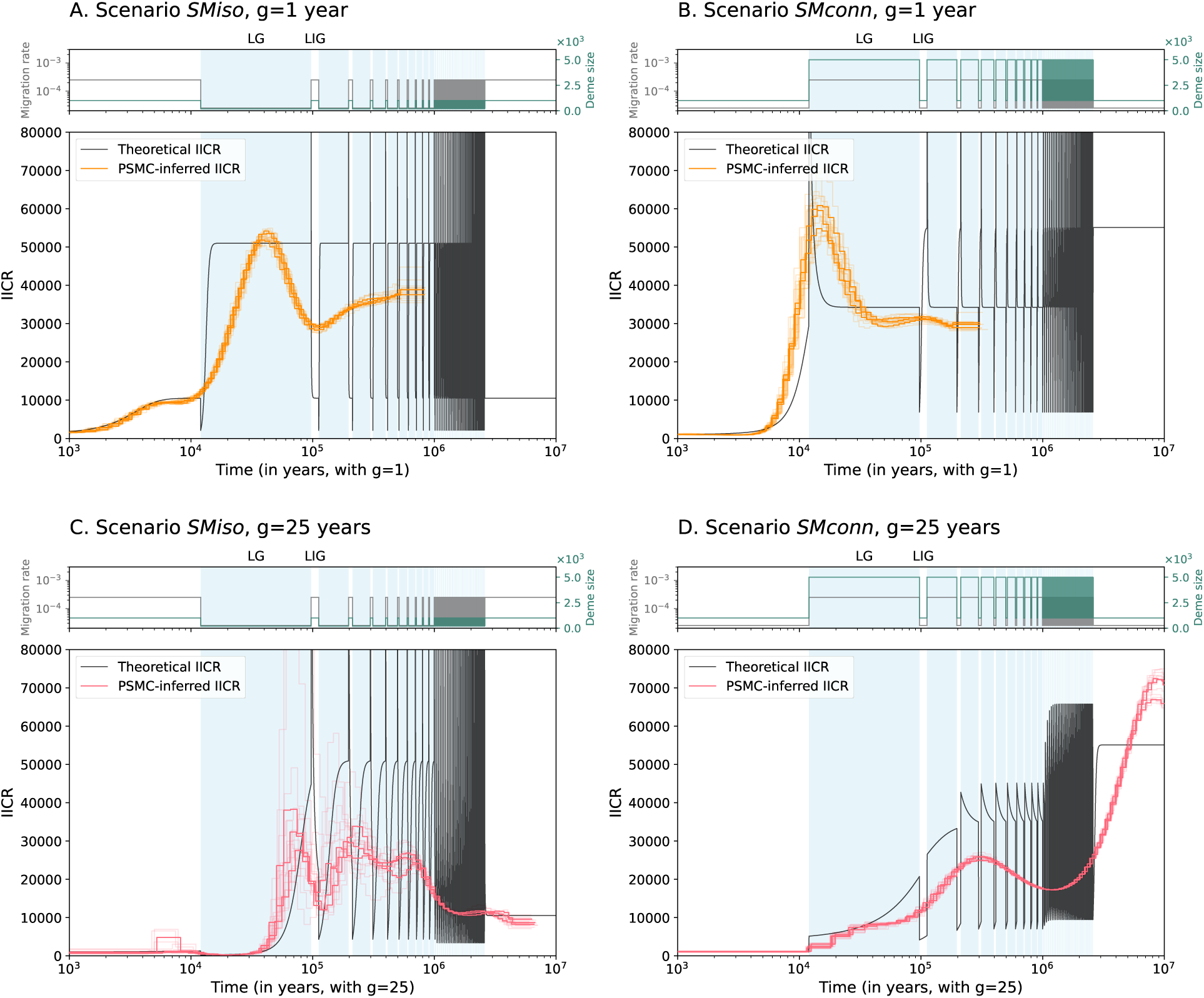
PSMC and theoretical IICR curves for demographic scenarios with oscillating migration rates and deme size (*SMiso* and *SMconn*). A: Scenario *SMiso* for a generation time of 1 year. B: Scenario *SMconn* for a generation time of 1 year. C: Scenario *SMiso* for a generation time of 25 years. D: Scenario *SMconn* for a generation time of 25 years. In each top panel, migration rates and deme sizes are represented over time with grey and turquoise lines, respectively. Migration fluctuates between *m* = 2.5.10^−5^ and *m* = 2.5.10^−4^, deme size fluctuates between *N* = 200 and *N* = 1000 (*SMiso*) or *N* = 1000 and *N* = 5000 (*SMconn*), and the number of demes *n* is constant and equal to *n* = 6. In each bottom panel, the blue curve represents the theoretical IICR, and the coloured curves correspond to the PSMC curve of one diploid genome sampled in the present and in the same deme for three repetitions of the simulations. The lighter PSMC curves correspond to five bootstraps per repetition. The pale blue areas in the background correspond to glacial periods.

While PSMC was able to recover the two to three most recent cycles, we note that the curves departed quite significantly from the theoretical IICR for times older than around 12 kya, and that the amplitude of the PSMC curves fluctuations were less marked than expected from the IICR oscillations. This was confirmed by a general pattern of increasing mean relative error (MRE) values as cycles get older (mean ± std across all tested scenarios: MRE_1-12 kya_ = 0.24 ± 0.13 vs. MRE_12kya-1Mya_ = 0.53 ± 0.48, Supp. Fig. S20-S25). Note however that quantitatively assessing PSMC’s ability to estimate the theoretical IICR and detect the oscillations was difficult, for reasons highlighted in the Material and Methods and Supplementary Information S1. Therefore, these error values should not be taken at face value.

Although drawing general conclusions from all simulated cases is challenging, the overall observation that PSMC responded to the fluctuations up to around 200 kya seems to hold for most cases (Supp. Fig. S14-S19), and has the potential to help identify qualitative changes in the IICR up to 300 ky ago for the longest generation time we tested, here 25 years. However, within the space of our simulated scenarios, the PSMC approach seemed unable to quantitatively or qualitatively infer older demographic oscillations.

### Inferring migration rates over time from PSMC curves with SNIF

The SNIF method was developed to infer the parameters of a piecewise stationary n-island model using a PSMC curve as a summary statistics (Arredondo et al., 2021). It allows to infer the number of demes *n* and their diploid size *N* (both assumed constant here) and the population scaled migration rates (*M_i_*) over time (see the Material and Methods section) (Arredondo et al., 2021). The SNIF-inferred theoretical IICR curves fit quite well the observed PSMC curves for all tested scenarios (Supp. Fig. S26 and S27). In addition, deme sizes were well inferred (1077 ± 317 diploids across all tested scenarios, true value = 1000). The number of demes was also reasonably well estimated for a generation time of one year for both scenarios *Miso* and *Mconn* (7.5 ± 1.6 demes, true value = 6), and were overestimated for the longer generation times (12.8 ± 5.7 demes, true value = 6). We note however that the estimated number of demes is not related to the generation time in any simple way. Importantly, the oscillations of the piece-wise migration rates were not correctly inferred, in particular for times older than the LG (> 97 kya) (Fig. 4, Supp. Fig. S26 and S27). If sometimes SNIF infers migration rates and the most recent demographic change (12 kya) reasonably well (Fig. 4A, Supp. Fig. 4B,C,D), it does not recover older demographic changes (*i.e.* > 12 kya) in any of the tested scenarios. In fact, except for the previously mentioned scenarios, SNIF always infers high connectivity for the times older than the last glacial period, sometimes even higher than the highest simulated migration rate. For instance, for *Mconn* with g = 1 year (Fig. 4B), the mean SNIF-inferred population scaled migration rate goes from 2.3 to 5 between 12 kya and 130 kya, while the highest simulated migration rate in that time period is 1 (thus between 2.3 to 5 times lower).

**Figure 4.**
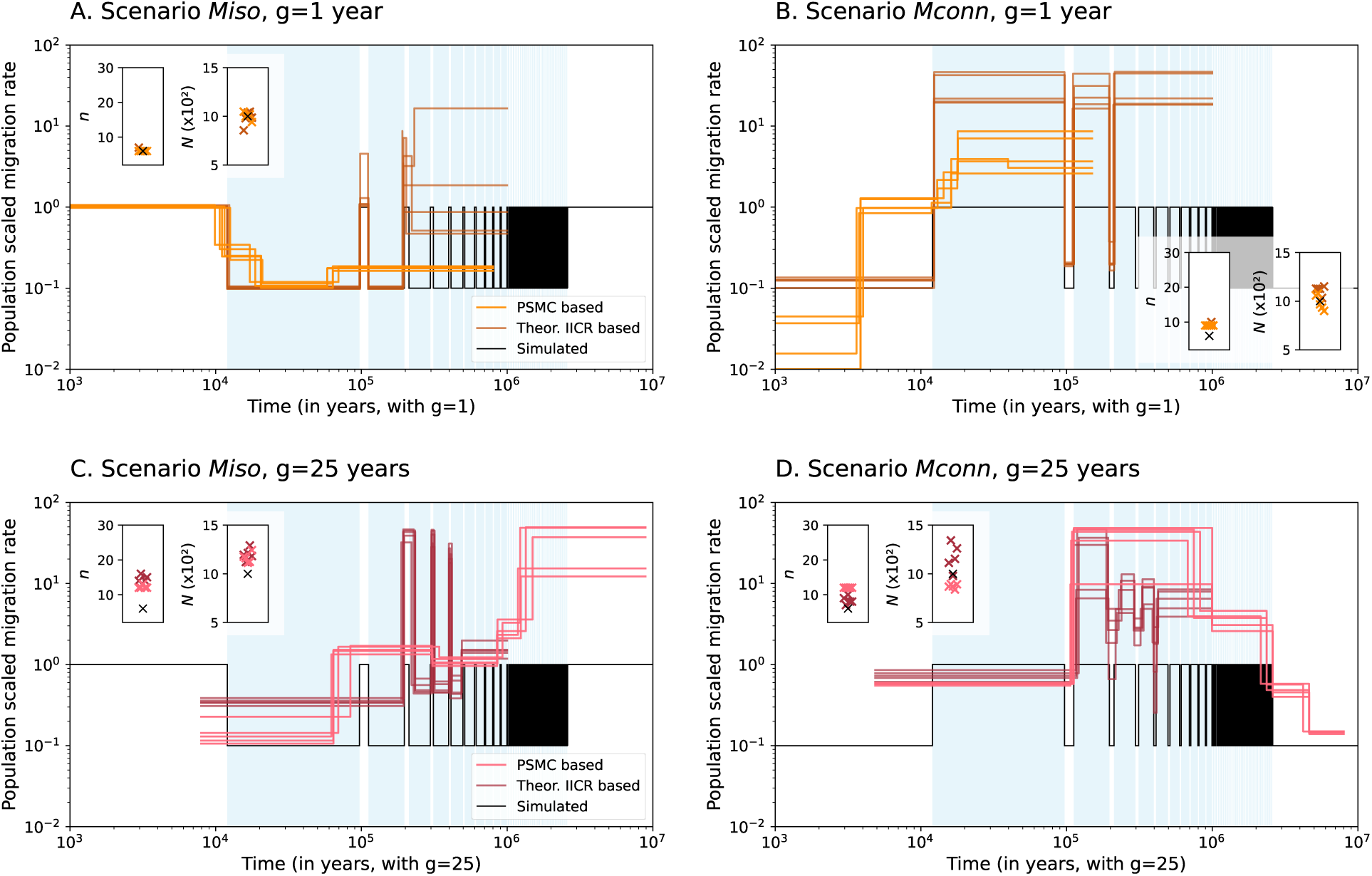
SNIF-inferred deme size *N*, number of islands *n* and population scaled migration rates *M_i_* (= 4*Nm_i_*) over time, using five repetitions of SNIF on PSMC and theoretical IICR curves for scenarios *Miso* and *Mconn*. A: Scenario *Miso* for a generation time of 1 year. B: Scenario *Mconn* for a generation time of 1 year. C: Scenario *Miso* for a generation time of 25 years. D: Scenario *Mconn* for a generation time of 25 years. Simulated migration oscillated between *m* = 2.5.10^−4^ (*M* = 1 given constant *N* = 1000) and *m* = 2.5.10^−5^ (*M* = 0.1), deme size *N* was constant and set to 1000 diploids, and the number of demes *n* was constant and equal to *n* = 6. In each panel, the black line corresponds to the target (simulated) migration rate over time, and the dark and light coloured curves correspond to the SNIF-inferred scaled migration rates using as input the theoretical IICR or a PSMC curve, respectively. Migration rates here correspond to *M_i_*= 4*Nm_i_*. The embedded plots show the inferred number of demes *n* (left) and the diploid deme size *N* (right). Black cross-points correspond to the target (simulated) values and the coloured cross-points correspond to the inferred values for each repetition of SNIF.

Given that PSMC curves are an approximation of the theoretical IICR, we hypothesised that the poor migration rate estimates obtained with SNIF come from the input PSMC curves, and not SNIF itself. To test this hypothesis, we further ran SNIF on the theoretical IICR curves of the demographic scenarios (darker coloured curves in Fig. 4). For computation reasons, we limited the inference to the ten most recent 100ky-long cycles, up to 1 Mya. SNIF can indeed fit the theoretical IICR curves up to 200 kya (Supp. Fig. S28A,B) and 500 kya (Supp. Fig. S28C,D). Even though the values of migration rates are mostly overestimated (e.g. Fig. 4B, brown curves, for which the highest migration rates are 20 to 50 times higher than the highest simulated migration rate, or Fig. 4C for which the lowest inferred migration rates are between 3 to 7 times higher than the lowest simulated migration rate), some oscillation patterns are recovered by SNIF for all four simulated scenarios. For a generation time of one year, SNIF infers the two (<200 kya) and three (<300 kya) most recent cycles for the scenario with isolating glacials (Fig. 4A in dark orange) and for the scenario with connecting glacials (Fig. 4B in dark orange), respectively. For a generation of 25 years, SNIF does not infer the most recent demographic oscillations but infers three cycles between 200 and 500 kya for the scenario with isolating glacial periods (Fig. 4C in dark pink) and between around 100 and 400 kya for the scenarios with connecting glacials (Fig. 4D in dark pink). We note however that, in all cases except *Miso* and g = 1 year, the amplitude of the oscillations are very poorly estimated.

### Comparing inferred trajectories for different generation times

To further understand the effect of generation time on PSMC and SNIF inferences, we compared PSMC curves obtained under the *same* demographic scenario (in units of years) across different generation lengths. Note that, therefore, simulated demographic scenarios were different in units of generations. Figure 5 shows PSMC curves obtained for scenarios with oscillating connectivity (between *m* = 2.5.10^−4^ and *m* = 2.5.10^−5^) and deme sizes (between *N* = 1000 and *N* = 5000) for five different generation times varying from 1 to 25 years. Comparative plots of PSMC curves for scenarios *M* and scenarios *SM* with different demographic parameters are shown in Supplementary Figures S29 and S30.

**Figure 5.**
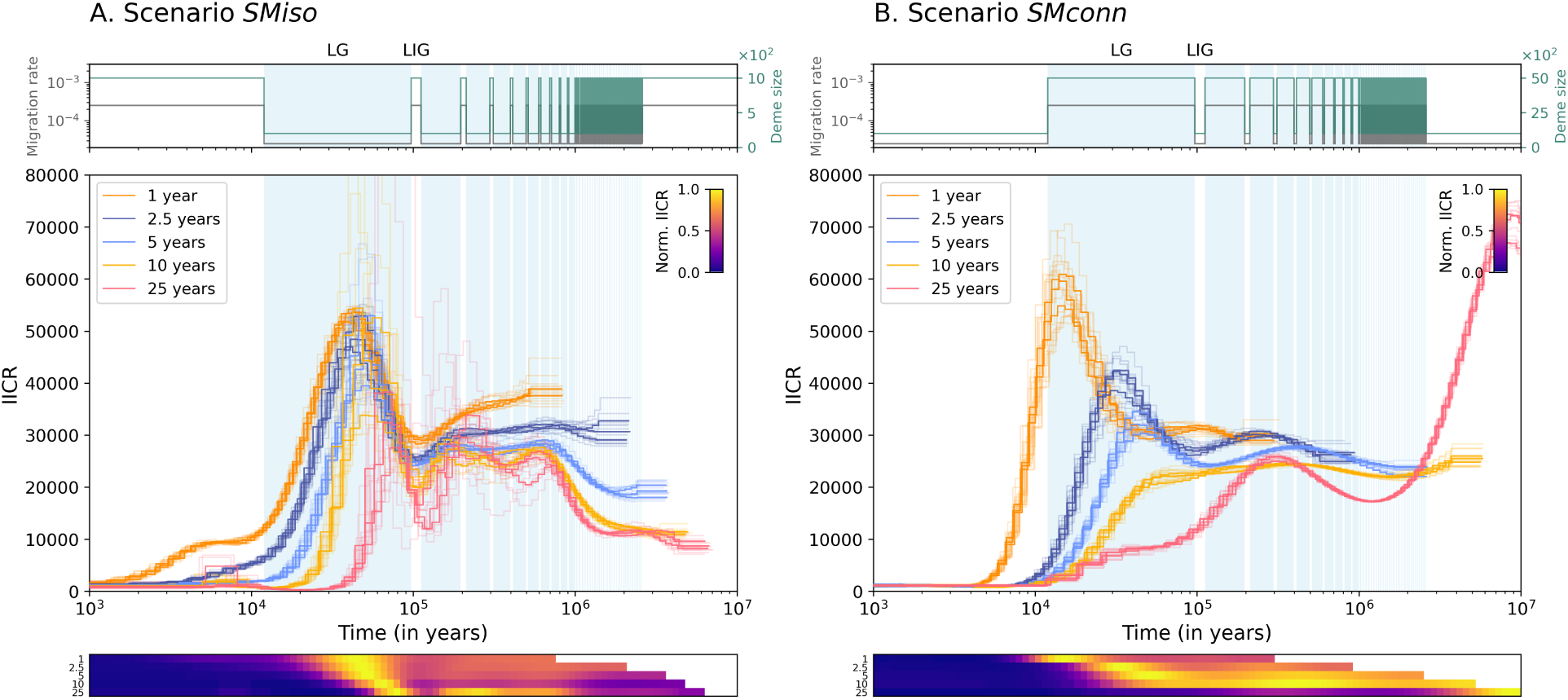
PSMC curves for scenarios with oscillating migration rates and deme sizes (scenarios *SM*) for different generation times. A: Scenario *SMiso*. B: Scenario *SMconn*. Migration oscillates between *m* = 2.5.10^−5^ and *m* = 2.5.10^−4^, deme size oscillates between *N* = 200 and *N* = 1000 (*SMiso*) or between *N* = 1000 and *N* = 5000 (*SMconn*), and the number of demes *n* was constant and equal to *n* = 6. Top panels show the simulated migration rates and deme sizes over time with grey and turquoise lines, respectively. Middle panels show the PSMC curves coloured by generation times: 1, 2.5, 5, 10 and 25 years in orange, dark blue, light blue, yellow and pink, respectively. Lighter curves correspond to bootstraps (five per PSMC curve). Blue rectangles in the background correspond to glacial periods. The heatmaps in the bottom panels represent the normalized PSMC-inferred IICR per time bin *t*, i.e. *PSMC_t_/PSMC_maximum_* for each generation time. The chromatic scale indicates normalized IICR values, with colors approaching yellow corresponding to higher IICR values, and those approaching dark blue indicating lower IICR values.

For scenarios with lower connectivity and deme size during the glacial periods (scenario *SMiso*, Fig. 5A), PSMC curves from 1 kya to the LIG (∼97-112 ya) show a similar pattern across generation times (going forward in time, from right to left): low IICR during the LIG, an increase at the beginning of the LG (97 kya), followed by a decrease during the LG. The timings of the fluctuations, however, do not coincide and are shifted to the present as generation time decreases. For instance, PSMC curves are at their maximum at the beginning of the LG (around 80 kya, see bottom panel in Fig. 5A) for a generation time of 25 years and then sharply decrease, whereas this maximum occurs in the middle of the LG (around 40 kya) for a generation time of one year and the curves decrease more gradually. For times older than the LIG, we observe further divergence between the PSMC curves: while curves for a 1-year generation time seem to decrease up to the LIG (forward in time), curves for a generation time of 2.5 and 5 years are more or less flat, whereas curves for a generation time of 10 and 25 undulate synchronously.

For scenarios with higher connectivity and deme size during the glacial periods (scenario *SMconn*, Fig. 5B), we note little overlap between all inferred PSMC curves, even for the period younger than the LIG. During the LG (12-97 kya) and forward in time (from right to left), PSMC curves show a continuous decrease in IICR for 10 and 25-year generation times, whereas they first increase at the beginning of the LG (97 kya) and then decrease for the generation times of 2.5 and 5 years, while it is stationary for the first half and then increases for a 1-year generation time. Again, the timings at which the trends change and the speed of IICR increase or decrease differ across generation times (see bottom panel in Fig. 5B). For instance, for the 5-year generation time, the maximum in IICR occurs in the middle of the LG at ∼40 kya, while it occurs at the very end of the LG ∼12 kya for 1-year generation length. Altogether, our results show that varying the generation length, and therefore varying the underlying biological scenarios in terms of generations, leads to fluctuations in the PSMC-estimated IICR that do not always coincide for species with variable generation times (see also Supp. Fig. S29 and S30). In particular, even if some curves exhibit similar patterns under the same scenario, they are not synchronized in time.

Comparing the two simulated scenarios *SMiso* and *SMconn* (isolating vs. connecting glacials respectively, left and right panels), we further note that PSMC curves exhibit similar shapes and trends for the generation times 2.5 and 5 years. However, for generation time of 1, 10 or 25 years, PSMC curves are quite different depending on whether migration and deme size increased or decreased during glacial periods (note that this was also observed in Fig. 3).

Looking now at the SNIF-inferred migration rates across the same five generation times (1, 2.5, 5, 10 and 25 years) for scenarios with oscillating migration rates and constant deme sizes (scenarios *M*, Fig. 6 for *m* oscillating between *m* = 2.5.10^−4^ and *m* = 2.5.10^−5^, with *N* = 1000), we find both consistent and inconsistent patterns in the connectivity graphs across the different generation times. For scenario *Miso* (Fig. 6A) and forward in time (from right to left), we observe a shared decrease in connectivity in the ancient past between 1 and 2 Mya for all generation times, except for g = 1 year with no *T*_2_ at this temporal horizon. Inferred migration rates remain overall constant then increase between 120 and 200 kya, this increases being sharper for g = 2.5 years (Fig. 6A, in dark blue) than for the other generation times. Finally, all curves decrease between 50 and 80 kya, slightly postdating the LIG-LG transition, 97 kya. In the recent past, SNIF further infers an increase in migration rate for a 1-year generation time, between 10 and 20 kya, which is not detected for the longer generation times.

**Figure 6.**
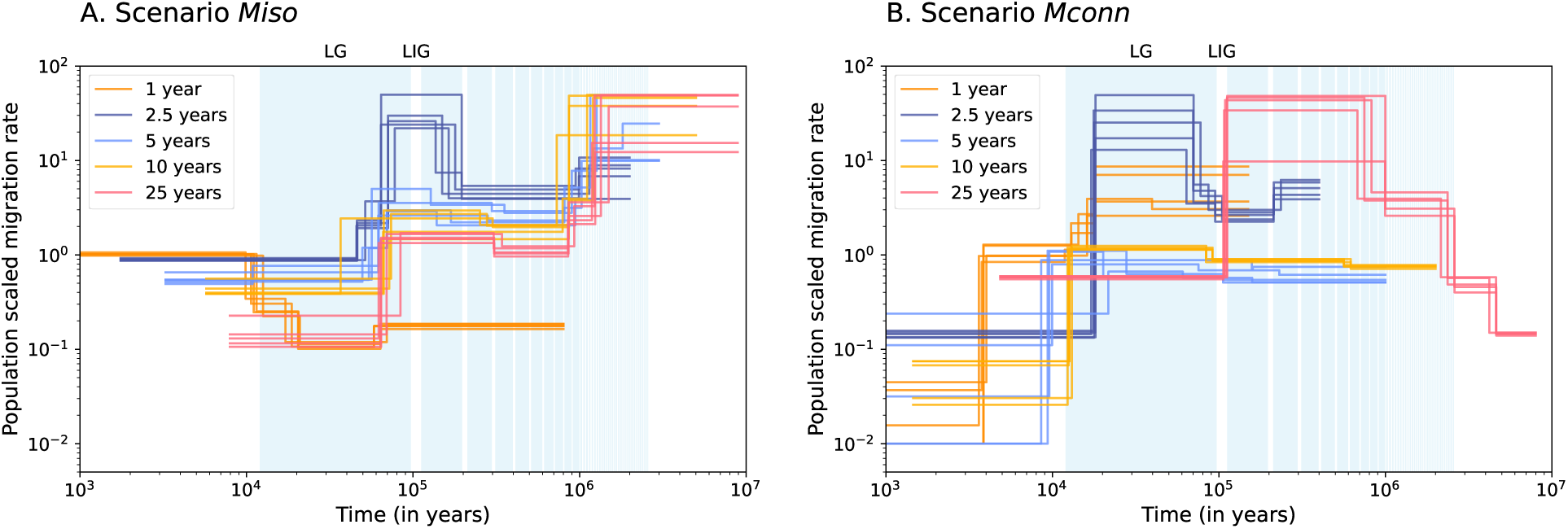
SNIF-inferred population scaled migration rates *M_i_* (= 4*Nm_i_*) using PSMC curves obtained under scenarios *M* for different generation times. A: Scenario *Miso*. B: Scenario *Mconn*. Migration oscillates between *m* = 2.5.10^−5^ and *m* = 2.5.10^−4^, deme size is constant and equal to *N* = 1000 and demes number was constant and equal to *n* = 6. Coloured curves correspond to one repetition of SNIF inference of a single PSMC curve, each colour corresponding to one generation time: 1, 2.5, 5, 10 and 25 years in orange, dark blue, light blue, yellow and pink, respectively. Blue rectangles in the background correspond to glacial periods. The PSMC curves input to SNIF correspond to the PSMC curves inferred under scenario *M* and shown in Supplementary Figure S29A-B.

Regarding scenario *Mconn* (Fig. 6B) and forward in time (from right to left), SNIF infers an increase in migration rate between 8 to 1 Mya for a generation length of 25 years, not observed for the shorter generation times due to the lack of *T*_2_ at this temporal horizon. We further observe for g = 25 years a sharp decrease in migration rates ∼110 kya. For g = 2.5, 5 and 10 years, inferred migration rates increase between 200 kya to 20 kya, with different magnitudes and timings. Finally, all inferred migration rates show a drop in connectivity towards the end of the LG ∼10-20 kya, for all generation times except 25 years. It looks as if the connectivity trajectory for g = 25 years was shifted towards the upper-right corner of the plot, i.e. to the older past and higher connectivity rates, compared to the lower generation times’ trajectories.

## Discussion

The objective of this work was to study the ability of PSMC-like methods to infer demographic oscillations by applying PSMC (and SNIF) to data simulated under repeated demographic changes in (i) migration rates alone or in (ii) migration rates and deme sizes altogether. We investigated how generation time influences the demographic inference of those cyclic changes, considering different generation times as different proxies for *in silico* species. Our rational was that, during the last 2.6 My, most species have likely seen their habitat and distribution range change with the climatic fluctuations of the Quaternary, which is especially true in areas of high climate change velocity (Sandel et al., 2011). These often dramatic habitat changes must have affected population size as well as connectivity between populations through time, both effects potentially leaving signatures in the genomes. We believe this work is important for several reasons. First, PSMC is one of the most widely used methods in conservation genetics and evolutionary biology to reconstruct the demographic history of species because it only requires one diploid genome, per species or population. Second, the theoretical work on the IICR carried out in previous studies and in the current one, provides significant improvements in our understanding of genomic diversity. However, previous studies have assumed that PSMC is a good estimator of the IICR, which was reasonable when the expected number of demographic events was limited (say less than eight) (Arredondo et al., 2021; Steux et al., 2025). Third, PSMC curves have been used in large comparative studies to make claims about past climatic and anthropogenic effects on many species declines or expansions (Bergman et al., 2023).

### The theoretical and simulated IICR carries information about the Quaternary demographic cycles

The IICR is a function of population size and migration rates, among other parameters, therefore it is likely to contain information regarding past demographic oscillations (Mazet et al., 2016). In our simulated scenarios, we have shown that the theoretical IICR indeed carries such information. However, this is not always the case, since the signal of recent demographic fluctuations seems to disappear for species with longer generation times (Fig. 2B, Supp. Fig. S3-S11).

The theoretical IICR can be estimated from the distribution of coalescent times (*T*_2_), however it is unclear whether the *T*_2_ distribution contains enough information about the past demographic cycles. We found that the IICR computed from independent *T*_2_ (“simulated IICR”) reproduces the theoretical IICR fairly well, as long as coalescent events are still available (Supp. Fig. S8-S13). We also showed in our simulations that oscillations older than ∼ 1 Mya are not seen in simulated IICR curves (Supp. Fig. S8-S13). This has likely two causes: first, that time has to be discretized to compute the IICR and, secondly, that the number of coalescence events across independent loci remaining to inform the inference decreases with time. Therefore, the resolution in the most ancient past (where fewer coalescent events remain), is not high enough to detect the repeated and short cycles occurring between 1 and 2.6 Mya. Overall, though, we show that both the theoretical and the simulated IICR curves carry information regarding cyclic demographic changes, especially so within a temporal horizon below 1 Mya, and could be used to develop an inferential framework, as long as both curves can be estimated with thin precision.

### PSMC lacks resolution in ancient times when climatic oscillations are frequent

PSMC is a powerful and useful method that allows researchers to estimate a complex statistic from the genome of a single individual. Simple scenarios have been tested in the original and later studies, and the method has proven powerful when species are (or are assumed to be) subjected to a small number of demographic changes (Li and Durbin, 2011; Steux et al., 2025). However, most species have likely been subjected to more than a few demographic changes in the last 2.6 My (as suggested by the 40-50 glacial cycles of the Quaternary), and our results suggest that the method is not able to identify so many cycles. Indeed, for most scenarios and generation times tested here, we found that PSMC estimates the IICR well at most the two to three most recent interglacial-glacial cycles, (i.e. during the last ∼ 100-300 kya), but has difficulty beyond such time periods. For instance, we found that for a generation time of 25 years, PSMC sometimes manages to infer the last two interglacial-glacial cycles (∼ 300 kya), though the IICR values do not entirely match the theoretical IICR expectation (see for instance Fig. 3B). However, older demographic cycles are not inferred by PSMC, and these observations are consistent across different values of migration rates (Supp. Fig. S14-S19). Furthermore, even when PSMC curves show some oscillations, the tempo of the variations rarely match the times at which actual changes in migration rate, or in migration rate and deme sizes, occur. A good example is the scenario *SMconn* with a generation time of 25 years (Fig. 3D): PSMC curves seem to follow a general trend of variation in IICR but, if we were to take these curves at face value, they would not correspond to any actual demographic change. Altogether, our results show that repeated demographic oscillations under our structured models cannot be inferred precisely and sometimes even approximately by PSMC. We note however that this issue is unlikely caused by the structured population model assumed in this work. Similar oscillations under a panmictic model would generate similar IICR curves whose oscillations would also be missed by PSMC for a large part. Furthermore, as demographic events get older and new ones occur, it is difficult to know which signals are left in the genomes and it is likely that the genomic signals of the older events get “over-written” by more recent ones. Overall, PSMC seems to infer a smoothed average across the simulated cycles, especially in the most ancient past (Supp. Fig. S14A,C,D,E,F between 100 and 800 kya for instance), suggesting that PSMC curves can recover some time-averaged information about the cycles, but will likely lose information about most individual cycles.

When we ran the SNIF method (Arredondo et al., 2021) on simulated PSMC curves, SNIF correctly inferred the most recent (12 kya) change in migration rates in 6 out of 10 scenarios (Supp. Fig. S26A,B,D and S27B,D,E). As expected, SNIF misses the other more ancient demographic events since it uses the PSMC curve as input. It also often seems to infer periods of high connectivity when connectivity actually fluctuated. When providing SNIF with theoretical IICR curves instead of simulated IICR or PSMC curves, SNIF could indeed detect several of the most ancient demographic oscillations. This confirms that SNIF can use information carried by the theoretical IICR regarding the demographic oscillations. This information however is to a large extent lost when simulating genomic sequences and running PSMC. This suggests that demographic scenarios as complex as the ones that could have occurred under the Quaternary oscillations might be difficult to recover from genomic data and PSMC curves alone. This also suggests that other sources of genomic information such as the AFS, containing information from multiple individuals, or more complex methods that estimate genealogy from multiple individuals (Kelleher et al., 2019), should be used together with PSMC, as suggested by several previous pieces of work (Ragsdale and Gravel, 2019; Chikhi et al., 2018; Rodríguez et al., 2018; Terhorst et al., 2017; Weissman and Hallatschek, 2017; Beaumont et al., 2002).

Overall, our simulations indicate that using PSMC curves as direct means to investigate species’ demographic history and response to past climatic oscillations is reliable only for time periods younger than 100 kya (for g = 1 year) to 200 kya (for g = 25 years). Yet, several studies have employed PSMC for this purpose (Bergman et al., 2023; Nadachowska-Brzyska et al., 2016; Kozma et al., 2016; Ruane et al., 2015; Prates et al., 2016; Bai et al., 2018; Sozzoni et al., 2025). For instance, Bergman et al. (2023) attempted to correlate the PSMC trajectories of 139 megafauna species with temperature and precipitation fluctuations over the past 700 ky. Finding no predictive power of these two variables on recent PSMC trends, they concluded that climate alone could not explain the species effective population size trajectories of these species over the last hundred thousand years. However, their study not only implicitly treated PSMC curves as an estimate of *N_e_* and a proxy for census population size, both of which are questionable assumptions (Mazet et al., 2016; Chikhi et al., 2018), but also did not assess whether PSMC was capable of accurately reconstructing such demographic cycles in the first place (Bergman et al., 2023). Several other studies have attempted to correlate PSMC trajectories with temperature, as well as with species past distribution areas using ecological niche modelling (ENM) (Sozzoni et al., 2025; Chattopadhyay et al., 2019; Brüniche-Olsen et al., 2021). For instance, Sozzoni et al. (2025) focused on 22 species of tortoises and terrapins and computed the correlation between PSMC curves (interpreted as effective population size, *N_e_*), and past distribution area or temperature (Sozzoni et al., 2025). They did so by choosing two time periods, from the MIS19 interglacial (∼787 kya) to the LIG and from the LIG to the LG, and computed the global trend of *N_e_* between the beginning and the end of each period (increasing, stable or decreasing) and correlated it with the same trend in temperature and/or distribution area. They did not find any correlation between *N_e_*and the distribution area, but found a correlation with temperature trend for their two time periods (Sozzoni et al., 2025). Our results suggest that between the LIG and the LG, PSMC curves correlate well with actual demographic fluctuations, at least for species with short generation times, suggesting that indeed PSMC is able to infer a demographic change during that time period. However, for the older time period between 1 Mya and the last couple of glacial periods ∼100-300 kya, their results seem at odds with our observations that PSMC does not correlate with the demographic (and therefore climatic) oscillations. We stress however that this study did not try to correlate *N_e_* with the temperature oscillations *per se*, but rather between the global temperature trend across 1 My, i.e. a time period during which temperature has oscillated a lot. Note also that the two time points they chose both correspond to an interglacial period, the MIS19 occurring ∼ 787 kya being thought to be the closest analog to the current Holocene (Vavrus et al., 2018). Overall, our results call for caution when using PSMC *N_e_* estimates for deep time periods (older than 100 kya to 200 kya depending on the generation time) under structured models to study species response to ancient climatic fluctuations. Under such complex scenarios, our work further emphasises the importance of validating inference results through simulations. Again, and as noted above, similar problems would also occur under panmixia if there were similar cycles of changes in population size. We also wish to stress the limits of our study, since we focused on *n*-island models with a small number of islands. We do not expect that models incorporating space such as stepping stones or larger numbers of demes would qualitatively change our results but we believe that caution is important and more work is needed to improve our understanding of the genetics of species that went through multiple glacial cycles.

### Different generation times lead to asynchronous PSMC trajectories under the same demographic cycles

Under the assumption that species’ demographic histories are primarily influenced by extrinsic factors such as temperature, species living in similar habitats are often expected to experience synchronous demographic fluctuations in response to climatic change (Hewitt, 2000, 2004). This assumption of temporal concordance has been broadly used in the field of phylogeography (Papadopoulou and Knowles, 2016), to test for instance whether local versus regional environmental disruptions shaped species demographic history (Gabrielli et al., 2024). If some studies have found shared and synchronous demographic trends among ecologically similar species (Chattopadhyay et al., 2019), a growing number of work however has shown rather idiosyncratic demographic responses to the late Quaternary oscillations (Ruane et al., 2015; Prates et al., 2016; Burbrink et al., 2016; Bai et al., 2018; Walton et al., 2021; Sozzoni et al., 2025). For instance, Burbrink et al. (2016) gathered data from 74 lineages across snakes, lizards, mammals, birds, turtles, salamanders and frogs distributed in the Nearctic realm, expecting shared and temporally concordant demographic histories during the recent Pleistocene (Burbrink et al., 2016). Using hABC (a coalescent-based method), they found both growing and declining demographic trends across their taxa, though expansion was the most common trajectory (75% of their species). Furthermore, the timing of expansion for the expanding species was synchronous on average for half of their species across their taxonomic groups (Burbrink et al., 2016). Another study, Ruane et al. (2015), focused on six species of milksnakes from temperate and tropical Americas and found independent species responses even among species adapted to the same climate (Ruane et al., 2015). PSMC inferences of Sozzoni et al. (2025) on the 22 species of tortoises and terrapins also show very variable demographic trends when differentiating temperate from tropical species (Sozzoni et al., 2025). Overall, these observed idiosyncratic responses suggest an effect of species-specific traits on top of extrinsic such as environmental changes, therefore questioning the expectation of concordant genetic patterns among species sharing similar habitat (Papadopoulou and Knowles, 2016; Germain et al., 2023).

In this work, we simulated simplistic demographic cycles in relation to the climatic oscillations of the Quaternary, assuming that climate could be modelled as changes in migration rate and deme size, and that climate was therefore the only or main predictor of demographic changes. Considering species that would only differ by their generation time, PSMC curves show asynchronous and sometimes idiosyncratic trajectories despite similar demographic fluctuations in units of years. This is not entirely surprising, given that under the coalescent theory, it is the number of generation that matters and not the number of years. Therefore, for a same cycle length in years, a cycle will be shorter for species with long generation time than for species with short generation time. Our results reinforce the idea that it might not be so relevant to interpret PSMC curves as a direct reconstruction of the evolutionary history of a species, and to compare different species’ PSMC irrespective of their species-specific traits. It might be more productive to use PSMC and IICR curves as summary statistics that are influenced by both extrinsic (climatic, environmental or anthropogenic) and intrinsic factors (species-specific traits) (Chikhi et al., 2018; Vishwakarma et al., 2025). Under this assumption, our results support the idea that comparative studies need to account for species-specific traits (in our case generation time) before we can draw conclusion on species genetic responses to the Quaternary oscillations (Papadopoulou and Knowles, 2016; Vishwakarma et al., 2025). Furthermore, our study indicates that drawing global demographic trends across different geographic and climatic regions for a wide range of taxa (likely to differ in their life-history traits) might not be the most appropriate and relevant way to study species responses to climate change.

We further note here that real species likely differ by many other parameters, such as their population density or the amplitude of the demographic changes they might have been subjected to. It might seem reasonable to assume that PSMC curves might differ even more if these parameters varied. However, there is also the possibility that some of these differences compensate each other and give similar genomic signals at least from a PSMC point of view. We plotted on Supplementary Figure S35 the PSMC curves of all the parameters we tested for this study, to see out of curiosity what the global trend would look like. We stress here that the parameters we tested do not correspond to a thorough exploration of the parameter space, we are simply showing some scenarios with various combinations of number of demes, deme sizes, migration rates values and oscillation amplitudes. Again, all the scenarios follow the same timeline, therefore all demographic changes occur at the same time in years. We observe that PSMC curves vary significantly. Interestingly, we see a global trend in the IICR in the recent past, with a sharper decrease starting between 10 and 100 kya (forward in time), as inferred in many species of the megafauna, some reptiles, fishes or birds (Li et al., 2021; Iannucci et al., 2021; Dong et al., 2021; Bergman et al., 2023; Sozzoni et al., 2025), though not in all bird species (Germain et al., 2023). The 10-100 kya period corresponds to the period when humans colonised different regions of the world, as well as the LG period. This kind of coincidence, which depends on properties of the IICR and PSMC, have been interpreted as (plausible) evidence of the impact of human activities by some authors cited Dong et al. (2021); Iannucci et al. (2021); Bergman et al. (2023). Interestingly though, avoiding the direct “visual” interpretation of PSMC curves and using SNIF (that accounts for generation time in the inferential process) to estimate migration rates over time, leads to detect synchronous changes that occurred since the LIG (Fig. 6). This is surprising given that SNIF gets its information from the input PSMC curves and that they do not fluctuate synchronously. For instance, for scenarios with low migration during glacials (*Miso*, Fig. 6A), SNIF could infer for the five *in silico* species (defined by their generation times) the decrease in migration rate at the beginning of LG (∼ 97 kya), despite the PSMC curves harbouring asynchronous behavior during that time period (Supp. Fig. S29A). Even more interestingly perhaps, on that same figure, SNIF also detected the most recent change in migration rate between the LG and the current interglacial for a generation time of 1 year. For scenarios with higher connectivity during glacials (*Mconn*, Fig. 6B), SNIF detected a relatively synchronous decrease in migration rate between the LG and the current interglacial (12 kya) for generation times lower or equal to 10 years, whereas it detected the decrease in migration rates between the LIG and the second-to-last glacial period (∼ 110 kya) for a generation time of 25 years. Thus, the timing of the inferred demographic events varied significantly depending on the generation time. This pattern is even more pronounced in SNIF inferences from theoretical IICR curves (Supp. Fig. S28), where shorter generation times led to higher temporal sensitivity to detect more recent cycles, while longer generation times resulted in higher sensitivity to detect older cycles. These findings suggest that it might not be meaningful to directly and visually compare PSMC curves across species with different generation times to infer shared demographic responses to climate change. However, co-analyzing their PSMC curves could potentially enhance our ability to detect demographic changes over a broader time scale (through the use of a meta-inferential approach like SNIF). It would also be interesting to see if using the extended version of PSMC on multiple sequences (MSMC (Schiffels and Durbin, 2014)) could help recover more recent events for species with longer generation time (Chikhi et al., 2024).

### Going beyond PSMC

Since its publication, the PSMC method has been widely used and with good reasons as, for instance, it allows researchers studying species potentially under threat to make use of genomic data and reconstruct past demography in an attempt to understand the drivers of population decline. However, interpreting the resulting curves remains challenging. As pointed out in previous studies (Mazet et al., 2016; Chikhi et al., 2018; Rodríguez et al., 2018; Arredondo et al., 2021), PSMC curves should be interpreted carefully for structured species and should rather be considered as a summary statistic of genomic information rather than a direct reconstruction of past demographic history. The SNIF method builds on this idea (Arredondo et al., 2021), using PSMC curves to infer migration rates over time. However, it makes the crucial assumption that PSMC can accurately infer the IICR. It has been shown in several studies that the PSMC method is indeed able to properly estimate the IICR but these studies were focused on a limited number of demographic changes (not always “cycles” *per se*) (Mazet et al., 2016; Chikhi et al., 2018; Arredondo et al., 2021; Steux et al., 2025). For instance, in Steux et al. (2025), we assumed that the observed PSMC curves from chimpanzees could be used to infer changes in connectivity in the different chimpanzee subspecies (Steux et al., 2025). We also showed with simulations of the inferred scenarios that the inferences were robust, testable and trustable. Our results in this current study suggest that some parameters inferred with SNIF are still trustable (for instance the deme size or, to a certain extend, the number of demes) within the same structured model framework, but that one should interpret the older past with caution, and that more work is needed before we understand the recent evolutionary history of chimpanzees. Our results suggest that, under complex cyclic demographic scenarios, PSMC does not estimate the IICR well. Therefore, if the IICR is informative about past demographic cycles of a population, PSMC curves provide limited information to infer the details of complex demographic histories from real genomic data.

To our knowledge, few of the other demographic inference methods have been tested on scenarios with repeated demographic changes under structured population models. In panmictic settings, methods such as MSMC and Stairway Plot 2 have been tested on scenarios involving two demographic cycles: (i) a first cycle between 10 kya and 100 kya, and (ii) a second cycle between 100 kya and 1 Mya, with both cycles successfully inferred. If both cycles appear to have the same length on a logarithmic time scale, they differ considerably on a linear time scale, making it difficult to reconcile them with the Quaternary climatic oscillations (Schiffels and Durbin, 2014; Liu and Fu, 2020). We conducted a brief analysis using Stairway Plot 2 (Liu and Fu, 2015, 2020) on some of our cyclic scenarios (Supp. Section S2), given that it remains a widely applied approach, although see some limitations of the method in Omarjee et al. (2026). Results were difficult to interpret, showing little consistency across replicates, likely because of the noise in the simulated SFS, the panmictic model assumed by the method and the complexity of our scenarios (Supp. Fig. S31 and S32). Nevertheless, our results suggest that Stairway Plot does not recover the cyclic demographic patterns, in line with the findings of Milesi et al. (2024) who also reported that population size oscillations in a panmictic setup were not reliably inferred by the method (see their Supplementary data, Milesi et al. (2024)).

In a yet unpublished work, Linck and Battey (2019) used Moments, another SFS-based method (Jouganous et al., 2017), to infer migration rates over time in a model with two populations either connected or fully isolated in a cyclic manner. If the method managed to discriminate models with continuous migration from models with cyclic migration, it did poorly at inferring the migration rate values over time (Linck and Battey, 2019). Finally, it is worth mentioning the study by Grant and Cheng (2012), who conducted simulations of temporal fluctuations in population size corresponding to variations in habitat availability for the red king crab, under a panmictic model. Their results showed that the Bayesian Skyline Plot method (a precursor to PSMC (Drummond et al., 2005)), failed to detect population size changes occurring prior to the LIG, likely due to the relatively small effective population sizes during that time period (Grant and Cheng, 2012).

Despite the limitations of PSMC, it is certain that PSMC curves remain highly informative summary statistics. In particular, under some parameter values (generation time, migration rates or deme sizes, Supp. Fig. S14-S19), PSMC curves exhibit different shapes whether glacial periods entailed more isolation and lowere deme size or, on the contrary, more connectivity and higher deme size for the same generation time. This suggests that PSMC curves could theoretically be used to discriminate between these two scenarios. PSMC has been used to that end before, for instance to compare the demographic response to past glaciations of closely related species (Vijay et al., 2018; Natesh et al., 2020), interpreting PSMC curves at face value. If we have shown that PSMC trajectories are hardly a direct representation of species demographic response to environmental changes, our results also suggest that these statistics could be useful, in an Approximate Bayesian Computation (ABC) framework for instance (Beaumont et al., 2002; Csilléry et al., 2010). Because summary statistics in ABC are not required to perfectly capture the entire evolutionary history but only to retain information relevant to specific hypotheses (Beaumont et al., 2002; Csilléry et al., 2010), PSMC curves could be exploited for distinguishing among alternative demographic models. Moreover, the information contained in PSMC curves could be enhanced by combining them with additional summary statistics such as the allele frequency spectrum (as pointed out earlier in this discussion). This type of work would likely further benefit from the integration of other type of data from other disciplines such as ecology, to account for the biology of the focus species and their specific life-history traits (Hoban et al., 2019).

Given the limits of current inferential frameworks to detect the demographic oscillations of the Quaternary, we further asked whether having a direct access to the past population genetic diversity would help recover these cycles. We numerically computed the expected within-deme heterozygosity over time under some of our scenarios using recurrence relation of heterozygosity (Maruyama, 1970; Alcala and Vuilleumier, 2014) and performed simulations to confirm these numerical trajectories (Supp. Section S3, and Supp. Fig. S33, S34) (Vishwakarma et al., 2026). In the tested scenarios (*Miso* and *Mconn* with *m* fluctuating between 2.5.10^−5^ and 2.5.10^−4^ for g = 1 and 25 years), temporal trajectories of genetic diversity followed the demographic oscillations, decreasing during periods of isolation and increasing during periods of connection. However, the amplitude of these fluctuations was a function of the generation time: for g = 1 year, genetic diversity varied between 2.8 (*Mconn*, Supp. Fig. S33B, time < 1 Mya) and 4.2 (*Miso*, Supp. Fig. S33A, time < 1 Mya) times higher during periods of high connectivity. For g = 25 years, amplitudes were smaller, with the genetic diversity being between 1.2 (*Mconn*, Supp. Fig. S34B, time < 1 Mya) and 1.7 (*Miso*, Supp. Fig. S34A, time < 1 Mya) times higher only. Therefore, these results suggest that demographic oscillations of the Quaternary could be recovered using historical or ancient samples, in particular for species with short generation times. In the case of long-lived species, more extensive sampling schemes (*i.e.* more samples from the same glacial and interglacial periods) would be required in order to discriminate noise from actual differences in genetic diversity.

Climatic oscillations during the Quaternary almost certainly induced cyclical demographic changes in many species (Hewitt, 2000, 2004), although the precise form and timing of these responses remain difficult to reconstruct. In this work, we assumed instantaneous demographic changes and limited ourselves to a small number of deme size / migration rate combinations, and we did not test for instance cases of negative density-dependence migration (Vishwakarma et al., 2026). Furthermore, we simulated only two different types of cycles (before and after 1 Mya), but the climatic oscillations have been more heterogeneous in amplitude and period (Lisiecki and Raymo, 2005). Nevertheless, demographic cycles must have occurred in one form or another. Explicitly incorporating cyclical models into inferential frameworks offers a promising way to better understand the genetic legacy of the Quaternary climate oscillations, and to characterize the demographic oscillations that many species are expected to have experienced (Hewitt, 2000).

## Material and Methods

### Simulated demographic scenarios

#### Timeline

We defined a simplified timeline of the climatic oscillations of the Quaternary as follows (Fig. 1A). From the present to 12,000 years ago (12 ky), we assumed an interglacial period corresponding to the Holocene. Then, from 12 kya to 1 Mya, we simulated a series of 10 cycles of 100 ky-duration each, with 85 ky-long glacial periods and 15 ky-long interglacial periods (Herbert, 2023; Shackleton and Opdyke, 1976; Palacios et al., 2022). After these 10 cycles, at approximately 1 Mya, we marked the mid-Pleistocene transition by simulating 38 shorter and symmetrical cycles up to approximately 2.6 Mya: cycles were 41ky long and equally divided into one interglacial and one glacial period, of 20.5 ky each (Shackleton and Opdyke, 1976; Clark et al., 2006). Finally, we stopped the cycles and assumed a single stationary interglacial period, as paleo-climatic records suggest that Pliocene climate was warmer and more stable than the Pleistocene (Lisiecki and Raymo, 2005; Palacios et al., 2022; Dowsett et al., 2009).

#### Demographic scenarios

We simulated a *n*-island model with a constant number of demes *n*. In the first set of scenarios (scenarios *M*), the only parameters allowed to change were the migration rates, alternating between interglacial and glacial periods (*m_IG_*and *m_G_*, respectively) while the deme size, *N*, remained constant. In the second set of scenarios (scenarios *SM*), both deme sizes and migration rates were allowed to alternate between periods with parameters (*m_IG_, N_IG_*) and (*m_G_, N_G_*) for the interglacial and glacial periods, respectively (Fig. 1B).

Because species respond to glacial periods depending on the type of habitat they are adapted to, we simulated two plausible demographic response to glacial periods. Either the populations were (i) more isolated during glacial periods (scenarios *Miso* with *m_G_* < *m_IG_*) along with contracted size (scenarios *SMiso* with *m_G_* < *m_IG_* and *N_G_* < *N_IG_*), or (ii) the populations were more connected during the glacial periods (scenarios *Mconn* with *m_G_* > *m_IG_*) along with expanded size (scenarios *SMconn* with *m_G_* > *m_IG_* and *N_G_* > *N_IG_*). Thus, scenarios *Miso* and *SMiso* correspond to temperate or tropical species that lived in more isolated (*Miso*) and also restricted areas (*SMiso*) during glacial periods (Sommer and Zachos, 2009; Arora et al., 2010), while scenarios *Mconn* and *SMconn* correspond to species that were more connected (*Mconn*) while also occupying wider ranges (*SMconn*) during colder conditions, for instance arctic species (Fedorov et al., 2020; Dalén et al., 2007) or insular species in archipelagos due to changes in sea level (Cibois et al., 2010) (though not always (Salces-Castellano et al., 2021)).

#### Model parameters

Migration rates *m_IG_* and *m_G_* were defined as the probability for a lineage in a subpopulation to migrate (backward in time) to another deme each generation, and therefore *M_IG_/*4 = *Nm_IG_*, *M_G_/*4 = *Nm_G_* for scenarios *M* and *M_IG_/*4 = *N_IG_m_IG_* and *M_G_/*4 = *N_G_m_G_* in scenarios *SM*, correspond to the number of emigrating (diploid) individuals at each generation. Deme sizes (*N* in scenarios *M*, and *N_IG_* and *N_G_* in scenarios *SM*) correspond to the number of diploid individuals in a deme. The number of demes *n* remained constant across all scenarios, and was set to *n* = 6 arbitrarily and for computational reasons.

Changes in migration rate and deme size were instantaneous, therefore varying in a step-like manner at each transition between interglacial and glacial periods. In scenarios *M*, we used the following migration rate values (per lineage per generation): *m_IG_, m_G_* ∈ {2.5 × 10^−5^, 2.5 × 10^−4^, 2.5 × .10^−3^}, therefore with *N* = 1,000 diploids: *M_IG_, M_G_* ∈ {0.1, 1, 10}. In scenarios *SM*, we used the same values for *m_IG_* and *m_G_* as for scenarios *M*, and deme size increased or decreased by a factor 2 or 5 during glacial periods. We set *N_IG_* to 1,000 diploids, and therefore *N_G_* ∈ {200, 500} (scenario *SMiso*) or *N_G_*∈ {2000, 5000} (scenario *SMconn*).

Throughout the paper, we will be mainly showing and discussing results for scenarios *M* for migration rates oscillating between 2.5 × 10^−5^ and 2.5 × 10^−4^ (and *N* = 1000 constant), and scenarios *SM* with migration rates oscillating between 2.5 × 10^−5^ and 2.5 × 10^−4^ and deme sizes 5-fold increasing or decreasing. All tested parameter sets not shown in the main text are available in Supplementary Material, except for the three scenarios *SM* where deme sizes increase or decrease by a factor 2, as their corresponding IICR and PSMC curves are similar to those of scenarios *M* with constant deme size.

### Simulation parameters

Simulations were carried out using msprime (Kelleher and Lohse, 2020; Baumdicker et al., 2022), with three repetitions for each scenario and set of parameters. For each parameterized scenario, we calculated the IICR by simulating one million coalescent times (*T*_2_) between two haploid individuals sampled at present in one deme. We designate this type of IICR as a “simulated IICR”, in comparison to the “theoretical IICR”, described in the next section. This simulated IICR was computed using python functions implemented in Rodríguez et al. (2018). To run PSMC, we also simulated one diploid individual sampled in one deme, consisting in 10 chromosomes of 100 Mb each, for a total genome length of 1 Gb. We used a recombination rate of 10^−8^ per lineage per generation and between adjacent base pairs, a mutation rate of 10^−8^ per lineage per generation per base pair, and a binary mutation model. Finally, we tested five different generation times: 1, 2.5, 5, 10 and 25 years per generation. Note that the generation time had no impact on the simulations *per se*, but impacted the times at which demographic changes occurred in units of generations, since paleoclimatic oscillations occurred at the same absolute (calendar) time for every simulation.

### Computing the theoretical IICR

We also calculated the theoretical IICR of each parameterized demographic scenario using the msprime function coalescence_rate_trajectory() (Kelleher and Lohse, 2020; Baumdicker et al., 2022). Conditioned on each parameterized scenario, we also computed two stationary theoretical IICR curves, corresponding to two stationary scenarios where either the interglacial or glacial parameters were kept constant. These two scenarios correspond to the two extreme scenarios which set limits to the oscillating one. This allowed us to compare the IICR under fluctuating demography with the expected IICR curves for each period, since the IICR curve of species going through the glacial cycles are expected to oscillate from one stationary IICR to the other, as shown in previous IICR studies (e.g., Arredondo et al. (2021)).

### Running PSMC

PSMC was run on the simulated sequences using the flags -N 25 -t 15 -r 1 -p “2+56*1+6” (Li and Durbin, 2011). The parameter *r*, the ratio of the recombination/mutation rates, was set to 1 as both mutation and recombination rates were identical in our simulations. Regarding the parameter *p*, the distribution of time windows for which an effective population size will be inferred, it was chosen to avoid on one hand over-fitting in the recent past which has been shown to occur with the default value (4+25*2+4+5) (Hilgers et al., 2025), and on the other hand to ensure that at least 10 coalescent events were identified to be within each time window. We ran five bootstrap replicates for each simulated genome to obtain an estimate of PSMC variance. To compare generation times, we computed the normalized PSMC per time bin *t*, computed as *PSMC_t_/PSMC_maximum_*, to obtain a visual representation of the PSMC-inferred IICR fluctuations and allow easier comparison between the different generation lengths (bottom panels in Fig. 5 and Supp. Fig. S29-S30).

### Quantifying PSMC performance

Quantifying PSMC performance is a difficult task for various reasons which we briefly mention here. One of them is that PSMC curves are typically obtained over a logarithmic scale for the x-axis, making it difficult to visually evaluate the fit between an expected IICR and the estimated PSMC curve, as noted by Arredondo et al. (2021). Here another issue stems from the fact that we simulated demographic cycles, questioning the time scale over which to compute an error. In the Supplementary Section S1 and Supplementary Figures S20-S25, we explain and provide plots for two options of error computation. The first option is a mean relative error (RE) computed from 1 kya to 1 Mya and averaged over each interglacial (15-ky-long) and glacial (85-ky-long) period. The second option is a mean RE computed at regular time intervals, every 999 years, also from 1 kya to 1 Mya. In practice, the RE was defined as 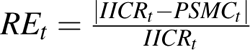. Details about these computations are given in Supplementary Section S1. We stress here that these errors should be interpreted with care, as we try to do in the main text.

### Running SNIF on the simulated PSMC curves

SNIF (Structured Non-stationary Inferential Framework, (Arredondo et al., 2021)) is an inferential method that assumes a piece-wise *n*-island model with varying migration rates and constant number and size of demes. Using a PSMC curve as the input summary statistic and splitting time in *c* different components indexed from 0 to *c* − 1, SNIF infers the diploid deme size *N*, the number of demes *n*, the scaled migration rate *M_i_* = 4*Nm_i_* for every time component *i* and the times at which migration rate changes from *M_i_*_−1_ to *M_i_* (therefore delimiting components *i* − 1 and *i*). We applied SNIF on PSMC curves simulated under the *M* scenarios but not the *SM* scenarios, as the latter involve changes in deme size that violate the SNIF model assumptions. For each scenario, we tested several values of *c* (the number of time components) going from 4 to 8, and *ω* (a parameter that weights the fit to the target PSMC) from 0.2 to 2. The best fits to the target PSMC curves were obtained using the parameters defined in the Supplementary Table S1. Finally, we ran five repetitions of the SNIF inference for each PSMC curve, as SNIF relies on a stochastic genetic algorithm optimization process.

### Running SNIF on the theoretical IICR curves

We further applied SNIF on the theoretical IICR of several scenarios to test whether SNIF could infer the demographic cycles from the information contained in these theoretical objects. Theoretical IICR represents the most ideal IICR curve, as it contains no stochastic noise nor sampling variance, what distinguishes it from simulated IICR or PSMC curves. The theoretical IICR should therefore carry the maximum information about past demographic changes, with the lowest (null) ratio of noise-over-signal regarding past demographic changes. For each scenario, we tested several values of *c* going from 4 to 8 and with *ω* equal to 1, 0.8, 0.5, 0.2, 0.1. Because of the high number of oscillations in the theoretical IICR, we only conducted the SNIF inferences up to 1 Mya, and up to 8 components, corresponding thus to a maximum of four cycles. We further specified time bounds for each change in migration rate to facilitate the fit to the input IICR curve. These bounds were determined based on preliminary bound-free inference runs, and they were kept broad not to constrain too much the inference. Time bounds and all the parameters that gave the best fits to the theoretical IICR inputs are shown in the Supplementary Table S2. Finally, we ran five repetitions of the SNIF inference for each IICR curve.

## Code Availability

Codes for the simulations and the inferences will be made available at the following github repository: https://github.com/camillesteux/inferringoscillations

## Acknowledgments

The authors thank Robin Aguilée, Marius Albino and Amandine Vidal-Hosteng for their helpful ideas and insightful discussions about the Quaternary oscillations and their impact on biodiversity patterns, as well as Abdelmajid Omarjee for his useful feedbacks on the SFS simulations and Stairway Plot 2 inferences. We acknowledge the financial support of a PhD studentship from the Ministère de l’Enseignement Supérieur et de la Recherche to CS. LC’s research was supported by the DevOCGen project, funded by the Occitanie Regional Council’s “Key challenges BiodivOc” program. This work was also supported by the LABEX entitled TULIP (ANR-10-558 LABX-41 and ANR-11-IDEX-0002-02) as well as the Investissement d’Avenir grant of the Agence Nationale de la Recherche (CEBA: ANR-10-LABX-25-01). We thank the IRP BEEG-B (International Research Project Bioinformatics, Ecology, Evolution, Genomics and Behaviour) for facilitating travel and collaboration between Toulouse (CRBE, IMT and INSA) and Lisbon (IGC and cE3c).

