## Supplementary Material for "Modeling and Inferring Large-scale Demographic Fluctuations in Structured Populations Through Simulations and PSMC-based Methods"

### Supplementary Information

#### S1 Quantifying PSMC performance

In an attempt to obtain a quantitative assessment of PSMC's ability to estimate the theoretical IICR, we tried to compute the relative error (RE) between the theoretical IICR and the PSMC curves. We represented in Supplementary Figure S20-S25 the error computed in two different ways for each scenario and set of parameters. First, we computed the mean RE across the three PSMC curves and the 15 bootstraps every 99.9 years (corresponding to 10,000 time points between 1 kya and 1 Mya linearly sampled), defined as $RE_t = \frac{|IICR_t - PSMC_t|}{IICR_t}$. We then computed the average of this mean RE per interglacial and glacial cycle: we computed one average RE for the period between 1 kya and 12 kya (the current interglacial), then for the LG between 12 kya and 97 kya, for the LIG between 97 and 112 kya, and so on, up to 1 Mya. This way of representing the error corresponds to the box-plots (Supp. Fig. S20-S25). Furthermore, because of the cyclic nature of the theoretical IICR (and to a certain extend the PSMC curves), we also represented a "continuous" error, corresponding to the mean RE across the three PSMC curves and the 15 bootstraps every 999 years (corresponding to 1,000 time points between 1 kya and 1 Mya linearly sampled). This "continuous" error is represented with the grey curve (Supp. Fig. S20-S25).

The objective of computing an error was to try to quantify the fit of the PSMC-inferred IICR to the theoretical IICR. However, note that due to the cyclic nature of our scenarios and the theoretical IICR oscillations, a relative error averaged across each interglacial or glacial period does not inform whether or not the PSMC-inferred IICR follows the oscillations of the theoretical object. For instance, in scenario *Miso* with $m_{IG} = 2.5 \times 10^{-4}$ and $m_G = 2.5 \times 10^{-5}$ for g >= 2.5 years (Supp. Fig. S22C,E,G,I), the average RE per time periods older than 12 kya are equal or lower than the mean RE for the period < 12 kya. However, looking at the corresponding PSMC curves (Supp. Fig. S16C,E,G,I), we see that although the PSMC curves depart further from the theoretical curves in the recent times (< 12 kya) than in the older times (as confirmed by the mean RE values), their shape is similar to the theoretical curve (i.e. increasing forward in time) whereas the PSMC curves completely miss the oscillations in the older times (> 12 kya). Therefore, a lower mean RE does not necessarily mean that the oscillations are better inferred, hence the difficulty to quantitatively assess PSMC performance under our cyclic scenarios.

#### S2 Stairway Plot 2 Analysis

Stairway Plot 2 is a demographic inference method that relies on the (1D) Site Frequency Spectrum (SFS), *i.e.* the distribution of the number of sites in a genome at given allele frequencies in a population (Liu and Fu, 2020). We only tested Stairway Plot on a subset of our scenarios for generation times of 1, 10 and 25 years, namely (i) scenario $M$ with migration rates fluctuating between $m = 2.5.10^{-5}$ and $m = 2.5.10^{-4}$ and constant deme size $N = 1000$, and (ii) scenario $SM$ with $m_{IG} = 2.5.10^{-4}$, $N_{IG} = 1000$ and $m_G = 2.5.10^{-5}$, $N_G = 200$ (*SMiso*) or (iii) scenario $SM$ with $m_{IG} = 2.5.10^{-5}$, $N_{IG} = 1000$ and $m_G = 2.5.10^{-4}$, $N_G = 5000$ (*SMconn*), .

We simulated 50 diploid genomes with msprime (Kelleher and Lohse, 2020; Baumdicker et al., 2022) using the same parameters as the simulations presented in the main text: each genome was made of ten chromosomes of 100 Mb each (for a total genome length of 1 Gb), recombination rate was set to $10^{-8}$ per lineage per generation and between adjacent base pairs, mutation rate was set to $10^{-8}$ per lineage per generation per base pair and the mutation model was binary. The 50 diploid genomes were sampled in the same deme and in the present, and the SFS was computed using a custom-made python script.

The model assumed by Stairway Plot is panmictic, and it is known that population structure creates U-shaped SFS (that is, an excess of high-frequency variants) that cannot be fitted by a panmictic model with only changes in population size under the Kingman coalescent (Freund et al., 2023). We therefore tested two ways to remove the U-shape of the SFS: we ran Stairway Plot on the folded SFS or on the unfolded SFS truncated to the frequency bin of 70 (parameter *largest_size_of_SFS_bin_used_for_estimation*), as suggested by Omarjee et al. (2026). Furthermore, note that results here will be interpreted as an inverse coalescent rate (and not an effective population size like Stairway Plot outputs (Liu and Fu, 2020)).

Though Stairway Plot inferred trajectories differed a little between using the folded or unfolded truncated SFS, our main observations are identical and we are therefore only showing results for inferences based on the folded SFS, in Supplementary Figures S31 (scenario $M$) and S32 (scenario $SM$). First, we observe some variations across replicates, likely coming from noise when simulating the SFS. Furthermore, Stairway Plot curves show several oscillations for all simulated cases, but they seem uncorrelated to actual simulated changes. It is particularly striking for the period between 1 to 12 kya for which Stairway Plot curves suggest demographic oscillations, despite the simulated demography being stationary. Therefore, it makes it difficult to interpret the inferred fluctuations for the period when demographic oscillations actually happen. For instance, for scenario $M$ with g = 25 years (S31E), the inferred Stairway Plot trajectories show two humps in the three replicates that could correspond to the underlying demographic cycles, but because humps appear even when the demography is stationary, it is hard to know whether they should be trusted or whether they coincide with simulated oscillations by chance.

Overall, Stairway Plot did not perform well on our simulated scenarios, and the results were difficult to interpret with regards to the demographic oscillations. This is likely due to a combination of issues, namely the simulated noisy SFS, the $n$-island model violating the panmictic model assumed by Stairway Plot, the complexity of our simulated scenarios, and finally, the fact that Stairway Plot itself is not always consistent in the scenarios it infers when given noisy SFS, as suggested in Omarjee et al. (2026).

#### S3 Following heterozygosity over time

Taking the opportunity of working with simulated data, we further asked if demographic cycles such as the ones we simulated in this study could be detected if we had a direct access to the past genetic diversity. To answer this question, we followed the work of Maruyama (1970); Alcala and Vuilleumier (2014); Vishwakarma et al. (2026) to compute the genetic diversity trajectories over time on some of our simulated scenarios: scenarios *Miso* and *Mconn*, with constant deme size $N = 1000$ diploids, constant number of demes $n = 6$ and migration rates fluctuating between $2.5.10^{-5}$ and $2.5.10^{-4}$.

The expected within-deme heterozygosity over time was computed numerically using formulas derived by Maruyama (1970); Alcala and Vuilleumier (2014) and used in Vishwakarma et al. (2026). Simulated genetic diversity was obtained with msprime, sampling 20 individuals in each sub-population (6 in total) every 1000 generations. The genetic diversity of each sample was computed on 50 independent replicates of 100-bp-long segments, using the function compute_diversity_samples() of msprime (Kelleher and Lohse, 2020; Baumdicker et al., 2022). Simulations were carried out using a mutation rate of $10^{-8}$ per lineage per generation per base pair, and a binary mutational model. The mean and standard deviation of the genetic diversity were computed across replicates for each sub-population (top and bottom left panels in Supp. Fig. S33 and S34), and the mean and standard deviation across replicates and across sub-populations were also computed (bottom right panels in Supp. Fig. S33 and S34).

Supplementary Figures S33 and S34 show that the expected within-deme genetic diversity increased or decreased according to the periods of high and low connectivity, respectively. This was also observed in the simulated trajectories, which reproduced well the numerical genetic diversity. Although there was significant variability between the trajectories of the different sub-populations, particularly for g = 25 years, the mean across replicates and sub-populations followed the demographic changes. Furthermore, we noted that the amplitude of the oscillations of the expected and simulated genetic diversity was dependent on the generation time. For g = 1 year, genetic diversity was between 2.8 (*Mconn*, Supp. Fig. S33B, time < 1 Mya) and 4.2 (*Miso*, Supp. Fig. S33A, time < 1 Mya) times higher during periods of high connectivity as compared to periods of low connectivity. For g = 25 years, amplitudes were smaller, with genetic diversity being between 1.2 (*Mconn*, Supp. Fig. S34B, time < 1 Mya) and 1.7 (*Miso*, Supp. Fig. S34A, time < 1 Mya) times higher during periods of high connectivity as compared to periods of low connectivity. Note that this was expected, as a longer generation time translates to shorter periods of high or low migration, limiting the amount of time during which genetic diversity can shift between equilibrium value.

Our results suggest that demographic oscillations of the Quaternary could be detected using historical or ancient samples, which is promising given the latest advancements in the technologies allowing ancient and historical DNA recovery (Raxworthy and Smith, 2021; Orlando et al., 2021). However, the power to detect these oscillations would depend on the focus species and its generation time (all else being equal): it would be easier to detect differences in genetic diversity between interglacial and glacial periods for short-lived species than for species with long generation time. For the latter, more samples from the same glacial and interglacial periods would be required in order to discriminate noise from actual difference in genetic diversity.

### Supplementary Tables

Table S1: Parameters used to run SNIF on simulated PSMC curves. Note that the time values are given here in years before present (column $t_i$), but they have to be divided by the generation time when actually given to SNIF.

| Scenario | g (years) | $c$ | $\omega$ | $n_{min}, n_{max}$ | $N_{min}, N_{max}$ | $M_{i,min}, M_{i,max}$ | $t_i$ (years) |
|---|---|---|---|---|---|---|---|
| | 1 | 4 | 1 | $2, 30$ | $10, 10000$ | $0.01, 50$ | $1.10^2, 8.10^5$ |
| | 2.5 | 5 | 0.2 | $2, 30$ | $10, 10000$ | $0.01, 50$ | $3.10^3, 5.10^6$ |
| *Miso* | 5 | 6 | 0.2 | $2, 30$ | $10, 10000$ | $0.01, 50$ | $3.10^3, 5.10^6$ |
| | 10 | 4 | 0.5 | $2, 30$ | $10, 10000$ | $0.01, 50$ | $3.10^3, 5.10^6$ |
| | 25 | 5 | 0.5 | $2, 60$ | $10, 10000$ | $0.01, 50$ | $5.10^3, 1.10^7$ |
| | 1 | 4 | 0.2 | $2, 30$ | $10, 10000$ | $0.01, 50$ | $1.10^2, 1.5.10^5$ |
| | 2.5 | 5 | 0.2 | $2, 30$ | $10, 10000$ | $0.01, 50$ | $3.10^3, 5.10^6$ |
| *Mconn* | 5 | 4 | 0.5 | $2, 30$ | $10, 10000$ | $0.01, 50$ | $3.10^3, 5.10^6$ |
| | 10 | 4 | 0.5 | $2, 60$ | $10, 10000$ | $0.01, 50$ | $8.10^2, 2.10^6$ |
| | 25 | 5 | 0.5 | $2, 60$ | $10, 10000$ | $0.01, 50$ | $3.10^3, 8.10^6$ |

g: Generation time (in years)
$c$: Number of SNIF components
$\omega$: Weight parameter for the SNIF distance computation
$n_{min}, n_{max}$: Lower and upper numbers of demes
$N_{min}, N_{max}$: Lower and upper deme sizes (in diploids)
$t_i$: Time bounds for the migration rate changes (in years). Here, time bounds were the same for all demographic changes.

Table S2: Parameters used to run SNIF on theoretical IICR. Note that here the times are given in years (column $t_i$), but they have to be divided by the generation time when given to SNIF.

| Scenario | g | $c$ | $\omega$ | $n_{min}$, $n_{max}$ | $N_{min}$, $N_{max}$ | $M_{i,min}$, $M_{i,max}$ | $t_i$ (years) |
|---|---|---|---|---|---|---|---|
| *Miso* | 1 | 6 | 0.2 | $2,30$ | $10,10000$ | $0.005,50$ | $(8.10^3, 2.10^4), (8.10^4, 1.5.10^5)$<br>$(8.10^4, 1.5.10^5), (1.5.10^5, 2.5.10^5)$<br>$(1.5.10^5, 2.5.10^5)$ |
| | 25 | 8 | 0.5 | $2,60$ | $10,10000$ | $0.005,50$ | $(1.5.10^5, 2.5.10^5), (1.5.10^5, 2.5.10^5)$<br>$(2.5.10^5, 3.5.10^5), (2.5.10^5, 3.5.10^5)$<br>$(3.5.10^5, 4.5.10^5), (3.5.10^5, 4.5.10^5)$<br>$(4.5.10^5, 5.5.10^5)$ |
| *Mconn* | 1 | 6 | 0.2 | $2,30$ | $10,10000$ | $0.005,50$ | $(8.10^3, 2.10^4), (7.10^4, 1.5.10^5)$<br>$(7.10^4, 1.5.10^5), (1.5.10^5, 2.5.10^5)$<br>$(1.5.10^5, 2.5.10^5), (2.5.10^5, 3.5.10^5)$<br>$(2.5.10^5, 3.5.10^5)$ |
| | 25 | 8 | 0.1 | $2,60$ | $10,10000$ | $0.005,50$ | $(1.10^4, 1.8.10^5), (1.8.10^4, 2.5.10^5)$<br>$(1.8.10^4, 2.5.10^5), (2.5.10^5, 3.5.10^5)$<br>$(2.5.10^5, 3.5.10^5), (3.5.10^5, 4.5.10^5)$<br>$(3.5.10^5, 4.5.10^5)$ |

g: Generation time (in years)
$c$: Number of SNIF components
$\omega$: Weight parameter for the SNIF distance computation
$n_{min}$, $n_{max}$: Lower and upper numbers of demes
$N_{min}$, $N_{max}$: Lower and upper deme sizes (in diploids)
$t_i$: Time bounds for the migration rate changes (in years). Here, each interval corresponds to one demographic change.

### Supplementary Figures

In the following we have plotted a number of figures that are mentioned throughout the manuscript and which we have tried to organize here is a way which we hope will be meaningful and helpful to the reader. In the next couple of pages we have tried to clarify the general logic of the groups of figures, reminding the readers what the different types of curves represent.

##### Theoretical IICR

The IICR is the theoretical object that PSMC aims to infer (Mazet et al., 2016; Li and Durbin, 2011), and can therefore be considered the expectation of PSMC inferences if PSMC was a perfect estimator and if molecular data perfectly represented coalescence time.

- **S1-S6**: Figures showing the theoretical IICR for the different tested demographic scenarios
- **S7**: Figure showing the effect of the cycle length on the amplitudes of the theoretical IICR oscillations

##### Simulated IICR

The theoretical IICR can be estimated from a finite number of $T_2$ values under a model of interest (here called "simulated IICR"), and can be seen as the best that PSMC could give if it inferred the exact distribution of $T_2$ from the distribution of heterozygous sites in the diploid genome. Simulated IICR curves were computed from $10^6$ values of independent $T_2$.

- **S8-S13**: Figures showing the simulated IICR curves for the different tested demographic scenarios

##### PSMC curves

Using whole-genome sequence data from a single diploid individual, PSMC estimates the distribution of coalescent times for two haploid samples (i.e. $T_2$) by walking along the diploid genome, identifying segments delimited by recombination points and inferring the T2 of each of these segments. The inferred distribution of $T_2$ is then translated into an estimate of the IICR (Mazet et al., 2016; Li and Durbin, 2011).

- **S14-S19**: Figures showing the PSMC curves obtained under the different tested demographic scenarios
- **S20-S25**: Figures showing the mean relative error of the PSMC curves in their estimation of the theoretical IICR curves

##### SNIF analyses

The SNIF method was developed to infer the parameters of a piecewise stationary n-island model using a PSMC curve as a summary statistics (Arredondo et al., 2021). It allows to infer the number of demes $n$ and their diploid size $N$ (both assumed constant here) and the population scaled migration rates ($M_i$) over time (see the Material and Methods section) (Arredondo et al., 2021).

- **S26-S27**: Figures showing SNIF inferences using PSMC curves as input data
- **S28**: Figure showing SNIF inferences using theoretical IICR curves as input data

##### PSMC curves of all simulated generation lengths

We compared the different PSMC curves obtained under the same demographic scenarios (in units of years) but different generation times.

- **S29-S30**: Figures showing the PSMC curves for all generation lengths

##### Stairway Plot inferences

Stairway Plot 2 is a demographic inference method that relies on the (1D) Site Frequency Spectrum (SFS), i.e. the distribution of the number of sites in a genome at given allele frequencies in a population (Liu and Fu, 2020). We tested Stairway Plot on a subset of our scenarios for generation times of 1, 10 and 25 years, to see if the method is able to detect the demographic oscillations of our cyclic demographic scenarios.

- **S31-S32**: Figures showing the Stairwayplot inferences tested on some of our demographic scenarios

##### Heterozygosity over time

We asked whether having a direct access to genetic diversity over time would allow us to recover the demographic cycles. Following the work of (Vishwakarma et al., 2026), we computed the expected and the simulated genetic diversity trajectories over time on some of our simulated scenarios.

- **S33-S34**: Figures showing the expected and simulated genetic diversity over time for some of our demographic scenarios

##### All PSMC curves

Finally, we plotted all the PSMC curves simulated in this study to observe the general trends of PSMC trajectories.

- **S35**: Figure showing all the PSMC curves simulated for this study

### Theoretical IICR

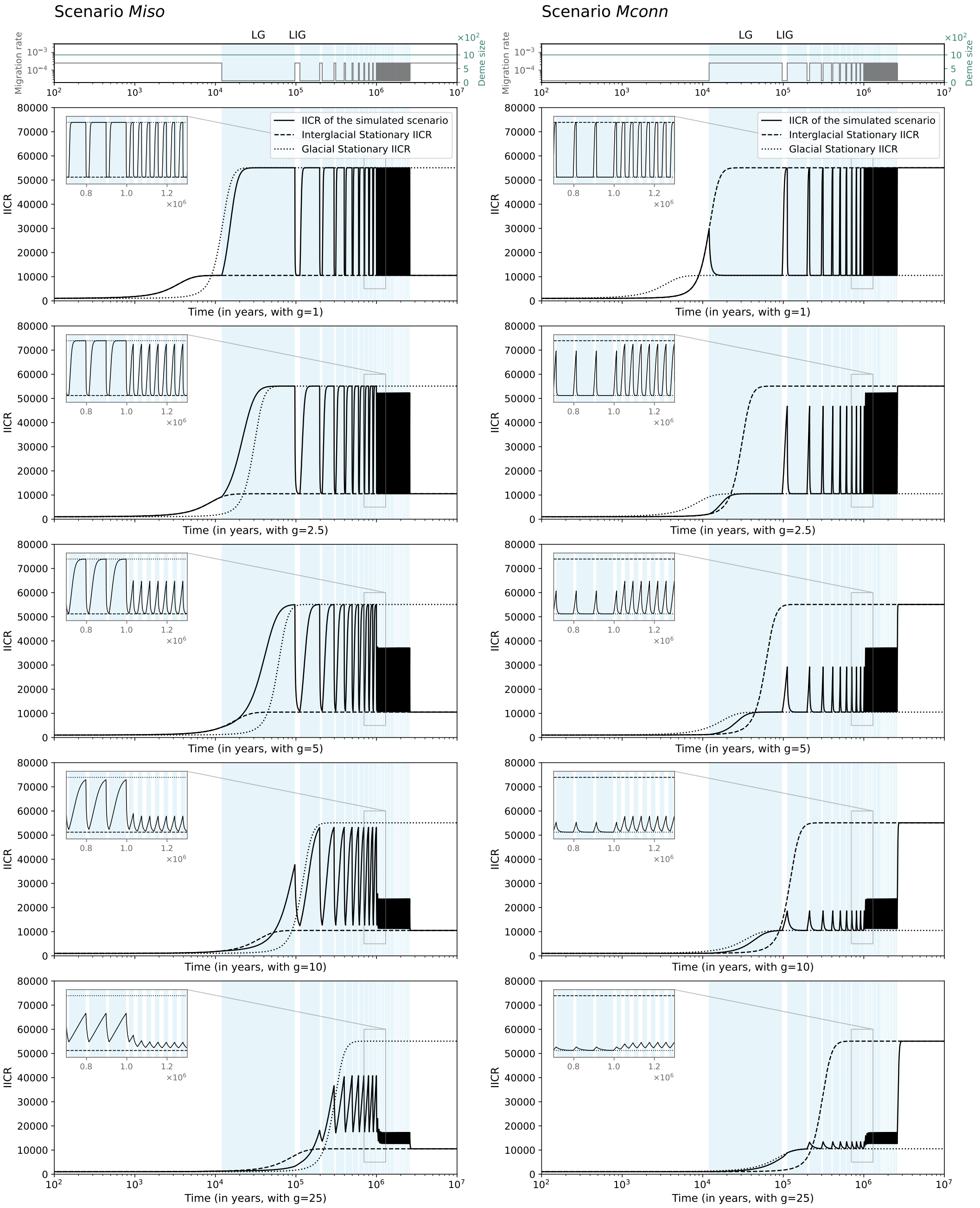


**Figure S1.** Theoretical IICR of the scenarios *M* for *m* varying between $2.5.10^{-5}$ and $2.5.10^{-4}$ for several generation times. The left column of panels corresponds to the scenario *Miso* and the right column represents the scenario *Mconn*. Top panels represent migration rates and deme size over time with grey and turquoise lines, respectively. Panels below represent the theoretical IICR for increasing generation times: 1 (second row), 2.5 (third row), 5 (fourth row), 10 (fifth row) and 25 (sixth row) years. Black dashed and dotted lines represent the stationary IICR corresponding to the migration rate of the interglacial and glacial periods, respectively. Continuous black lines correspond to the IICR of the oscillating scenario. The pale blue areas in the background correspond to glacial periods. LG and LIG stand for Last glacial and Last interglacial, respectively.

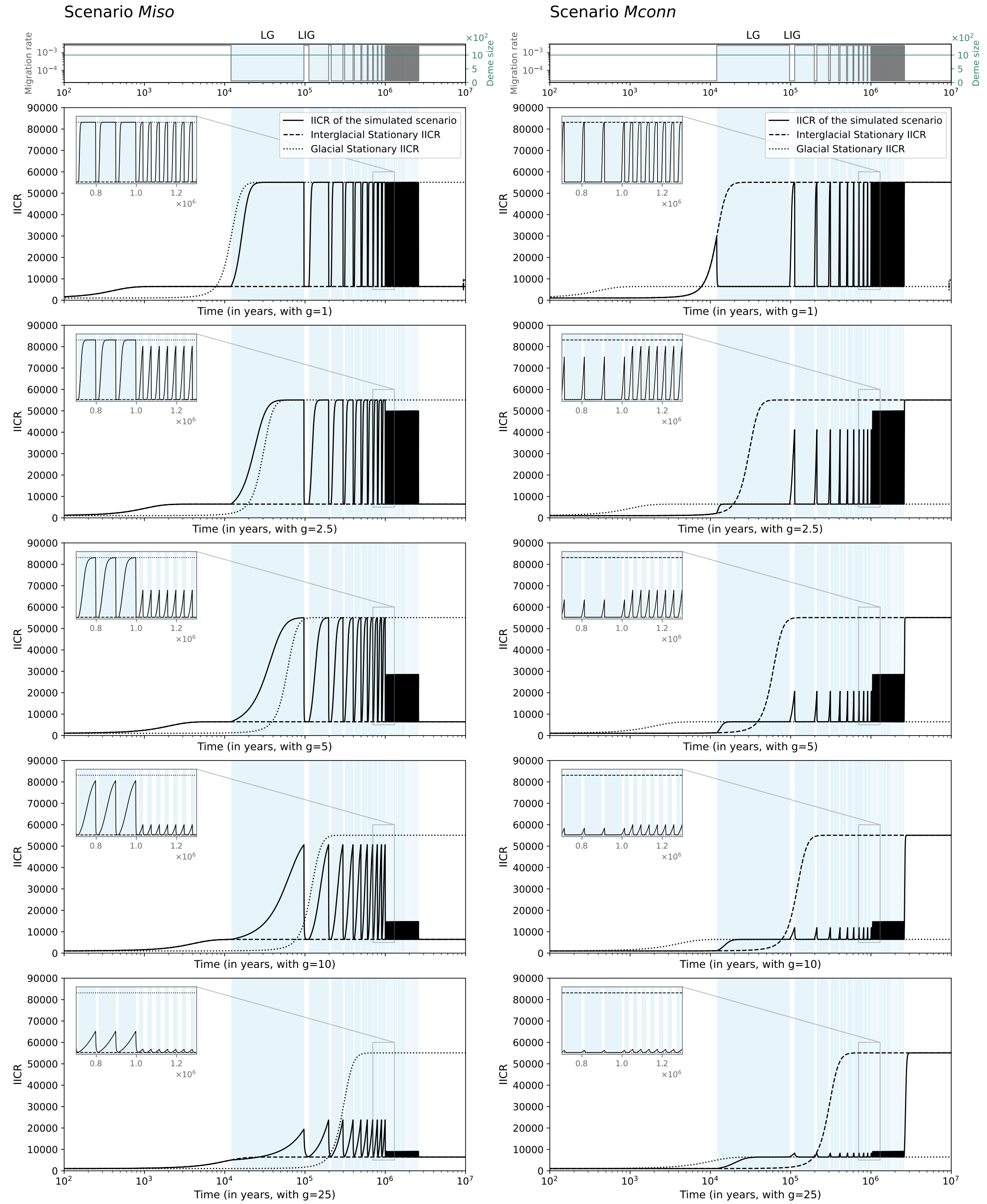


**Figure S2.** **Theoretical IICR of the scenarios *M* for $m$ varying between $2.5.10^{-5}$ and $2.5.10^{-3}$ for several generation times.** The left column of panels corresponds to the scenario *Miso* and the right column represents the scenario *Mconn*. Top panels represent migration rates and deme size over time with grey and turquoise lines, respectively. Panels below represent the theoretical IICR for increasing generation times: 1 (second row), 2.5 (third row), 5 (fourth row), 10 (fifth row) and 25 (sixth row) years. Black dashed and dotted lines represent the stationary IICR corresponding to the migration rate of the interglacial and glacial periods, respectively. Continuous black lines correspond to the IICR of the oscillating scenario. The pale blue areas in the background correspond to glacial periods. LG and LIG stand for Last glacial and Last interglacial, respectively.

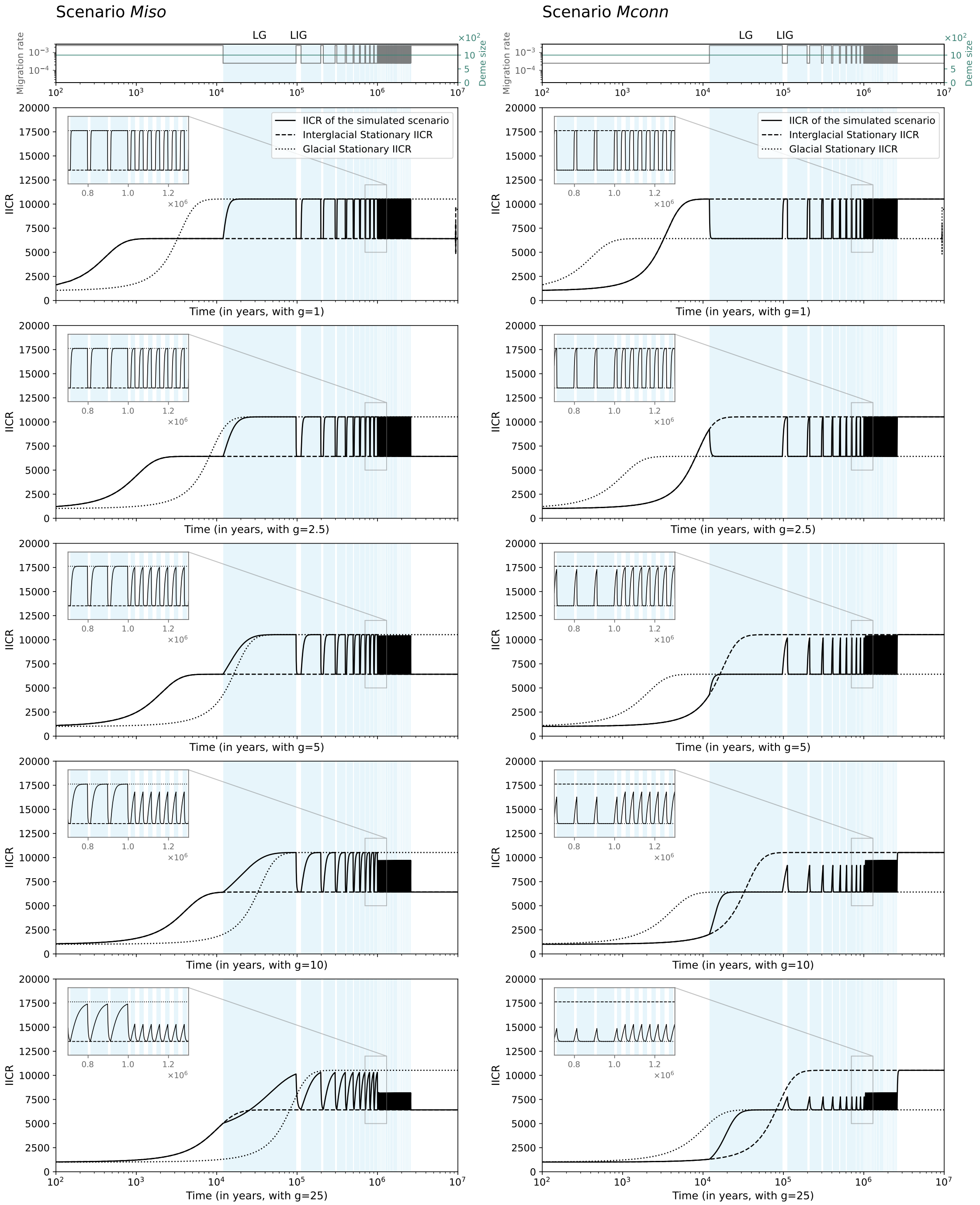


**Figure S3.** **Theoretical IICR of the scenarios *M* for $m$ varying between $2.5.10^{-4}$ and $2.5.10^{-3}$ for several generation times.** The left column of panels corresponds to the scenario *Miso* and the right column represents the scenario *Mconn*. Top panels represent migration rates and deme size over time with grey and turquoise lines, respectively. Panels below represent the theoretical IICR for increasing generation times: 1 (second row), 2.5 (third row), 5 (fourth row), 10 (fifth row) and 25 (sixth row) years. Black dashed and dotted lines represent the stationary IICR corresponding to the migration rate of the interglacial and glacial periods, respectively. Continuous black lines correspond to the IICR of the oscillating scenario. The pale blue areas in the background correspond to glacial periods. LG and LIG stand for Last glacial and Last interglacial, respectively.

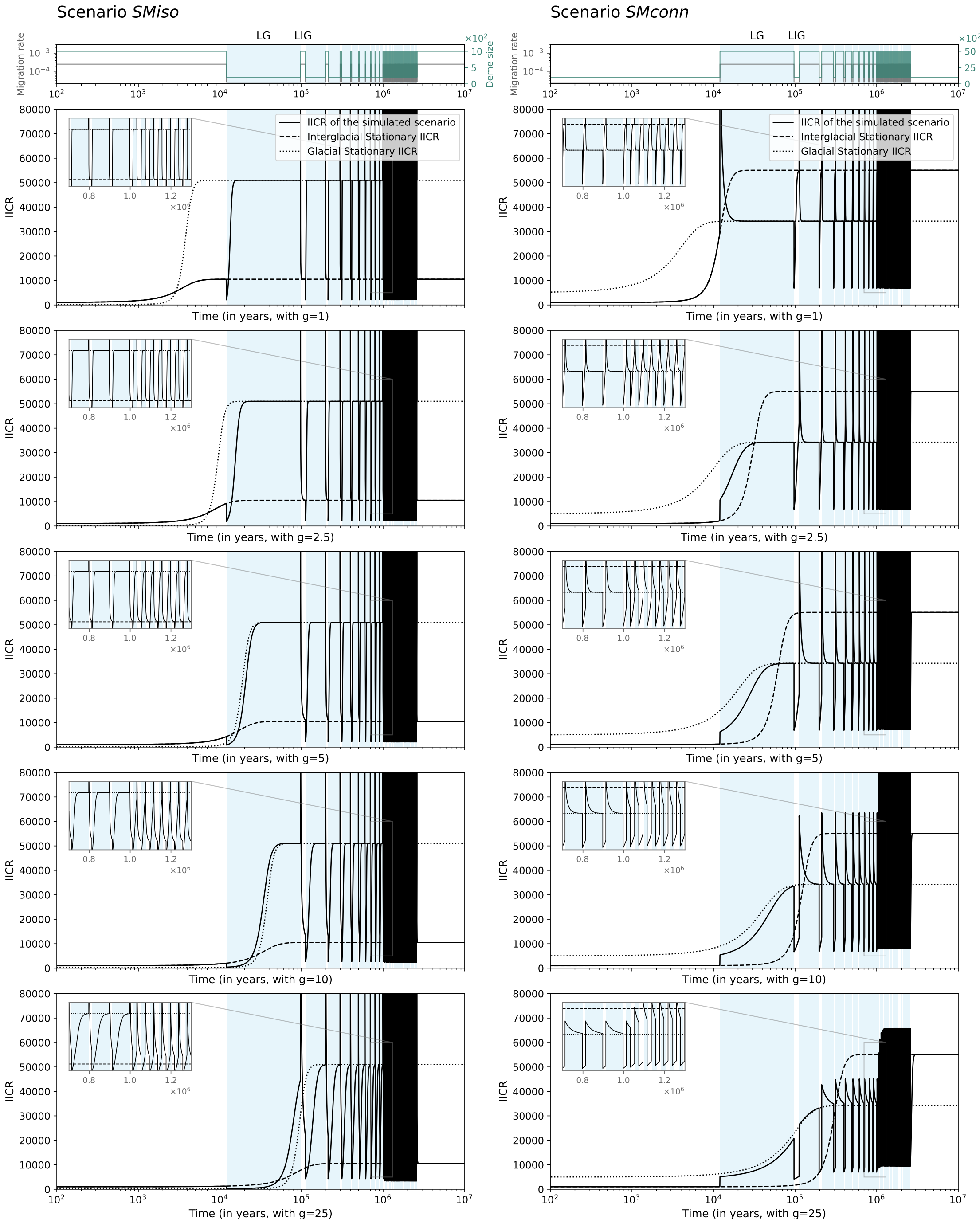


**Figure S4. Theoretical IICR of scenario *SM* for $m$ varying between $2.5.10^{-5}$ and $2.5.10^{-4}$ and $N$ 5-fold increasing or decreasing during glacials, for several generation times.** The left column of panels corresponds to the scenario *SMiso* with $m_{IG} = 2.5.10^{-4}$, $N_{IG} = 1000$ and $m_G = 2.5.10^{-5}$, $N_G = 200$, and the right column represents the scenario *SMconn* with $m_{IG} = 2.5.10^{-5}$, $N_{IG} = 1000$ and $m_G = 2.5.10^{-4}$, $N_G = 5000$. Top panels represent migration rates and deme size over time with grey and turquoise lines, respectively. Panels below represent the theoretical IICR for increasing generation times: 1 (second row), 2.5 (third row), 5 (fourth row), 10 (fifth row) and 25 (sixth row) years. Black dashed and dotted lines represent the stationary IICR corresponding to the migration rate and deme size of the interglacial and glacial periods, respectively. Continuous black lines correspond to the IICR of the oscillating scenario. The pale blue areas in the background correspond to glacial periods. LG and LIG stand for Last glacial and Last interglacial, respectively.

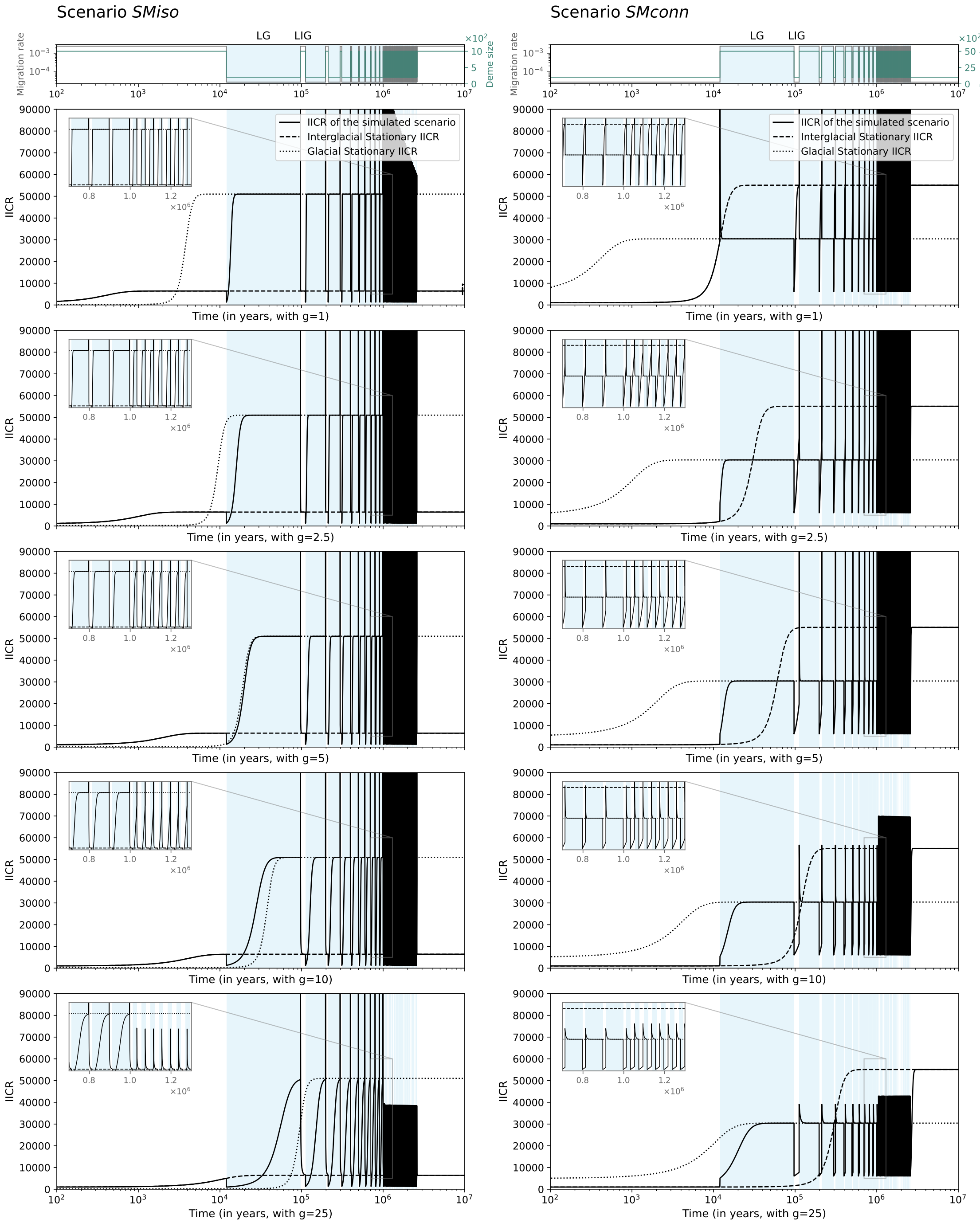


**Figure S5.** **Theoretical IICR of scenario *SM* for $m$ varying between $2.5.10^{-5}$ and $2.5.10^{-3}$ and $N$ 5-fold increasing or decreasing during glacials, for several generation times.** The left column of panels corresponds to the scenario *SMiso* with $m_{IG} = 2.5.10^{-3}$, $N_{IG} = 1000$ and $m_G = 2.5.10^{-5}$, $N_G = 200$, and the right column represents the scenario *SMconn* with $m_{IG} = 2.5.10^{-5}$, $N_{IG} = 1000$ and $m_G = 2.5.10^{-3}$, $N_G = 5000$. Top panels represent migration rates and deme size over time with grey and turquoise lines, respectively. Panels below represent the theoretical IICR for increasing generation times: 1 (second row), 2.5 (third row), 5 (fourth row), 10 (fifth row) and 25 (sixth row) years. Black dashed and dotted lines represent the stationary IICR corresponding to the migration rate and deme size of the interglacial and glacial periods, respectively. Continuous black lines correspond to the IICR of the oscillating scenario. The pale blue areas in the background correspond to glacial periods. LG and LIG stand for Last glacial and Last interglacial, respectively.

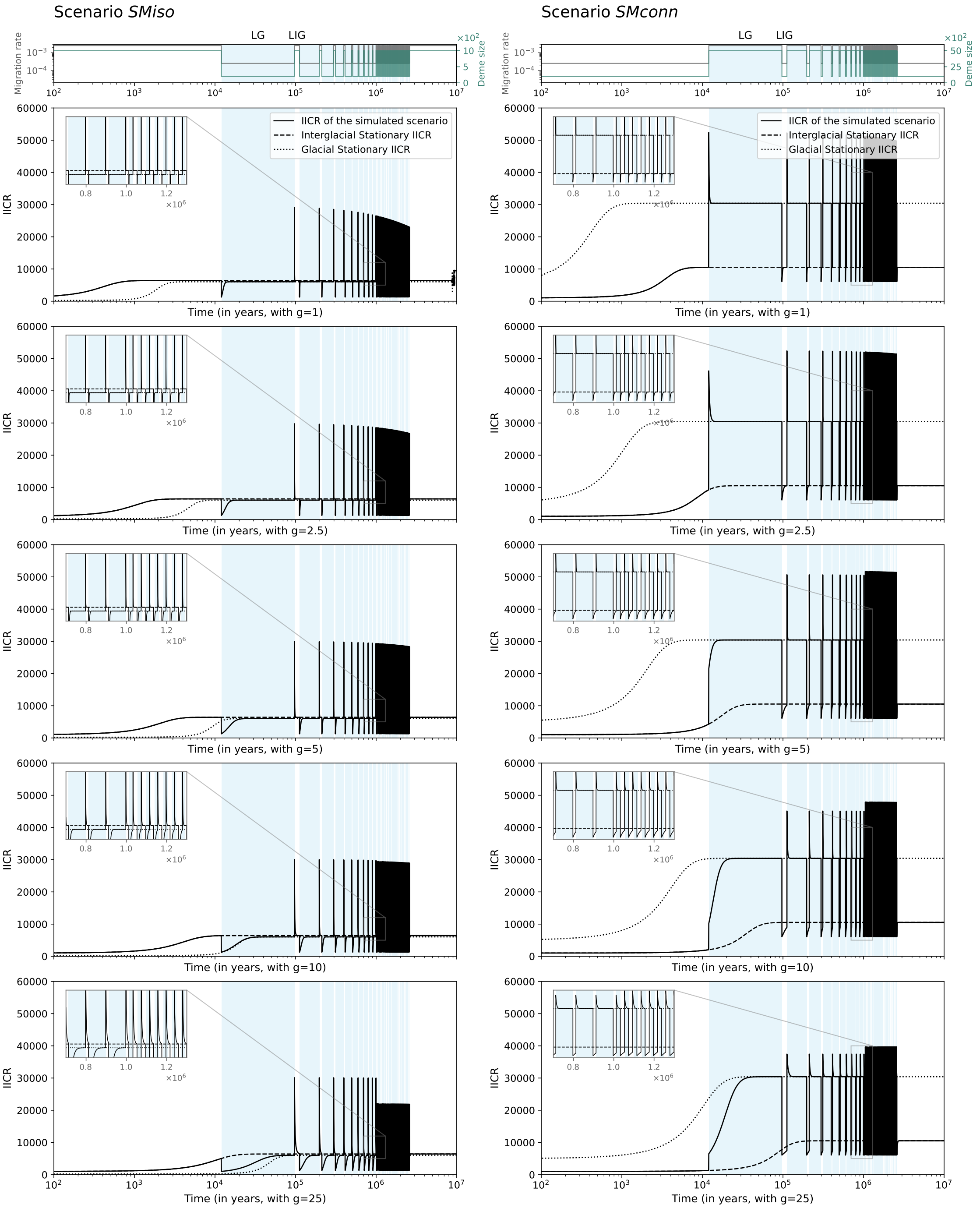


**Figure S6. Theoretical IICR of scenario *SM* for $m$ varying between $2.5.10^{-4}$ and $2.5.10^{-3}$ and $N$ 5-fold increasing or decreasing during glacials, for several generation times.** The left column of panels corresponds to the scenario *SMiso* with $m_{IG} = 2.5.10^{-3}$, $N_{IG} = 1000$ and $m_G = 2.5.10^{-4}$, $N_G = 200$, and the right column represents the scenario *SMconn* with $m_{IG} = 2.5.10^{-4}$, $N_{IG} = 1000$ and $m_G = 2.5.10^{-3}$, $N_G = 5000$. Top panels represent migration rates and deme size over time with grey and turquoise lines, respectively. Panels below represent the theoretical IICR for increasing generation times: 1 (second row), 2.5 (third row), 5 (fourth row), 10 (fifth row) and 25 (sixth row) years. Black dashed and dotted lines represent the stationary IICR corresponding to the migration rate and deme size of the interglacial and glacial periods, respectively. Continuous black lines correspond to the IICR of the oscillating scenario. The pale blue areas in the background correspond to glacial periods. LG and LIG stand for Last glacial and Last interglacial, respectively.

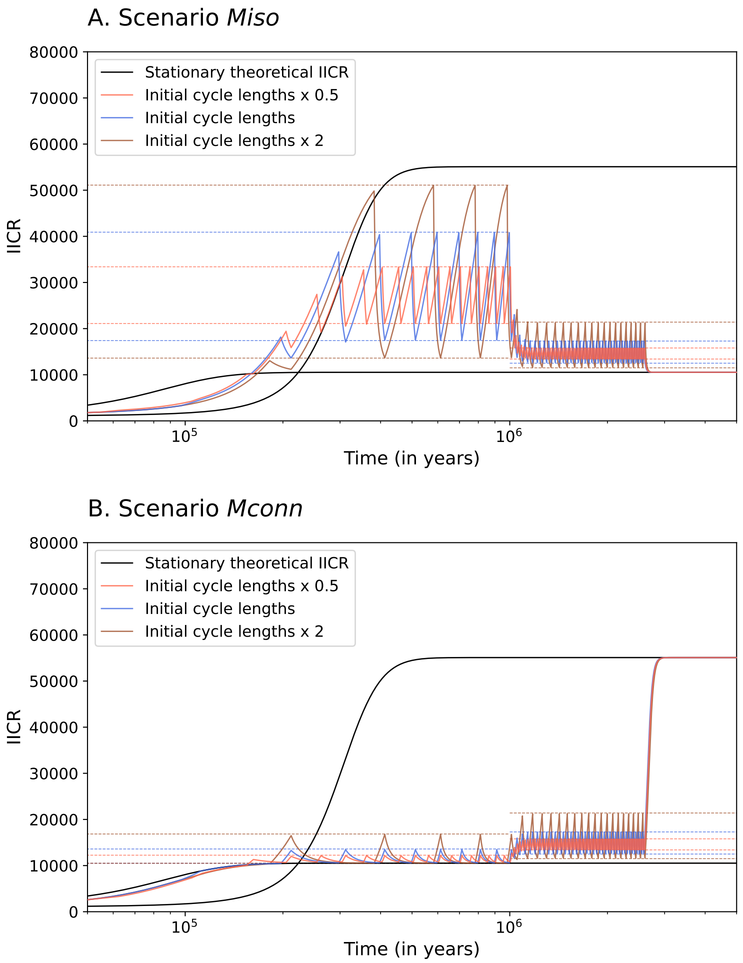


**Figure S7. Theoretical IICR for a generation time of 25 and different cycle lengths.** The initial cycle lengths correspond to the lengths described in Figure 1, ie. cycles of 100 ky from the present up to 1 Mya with 15ky-long interglacials and 85ky-long glacials, and cycles of 41 ky from 1 Mya to 2.6 Mya with 21.5ky-long interglacials and 21.5ky-long glacials. The black plain curves correspond to the stationary theoretical IICR and the coloured curves correspond to the theoretical IICR of the scenarios with oscillating migration rate (scenarios *Miso* and *Mconn*) with $m_{IG} = 2.5.10^{-4}$ and $m_G = 2.5.10^{-4}$ (A) or with $m_{IG} = 2.5.10^{-5}$ and $m_G = 2.5.10^{-4}$ (B). Dashed lines correspond to the "pseudo-plateau" between which the IICR oscillates and which depend on the cycle lengths.

### Simulated IICR

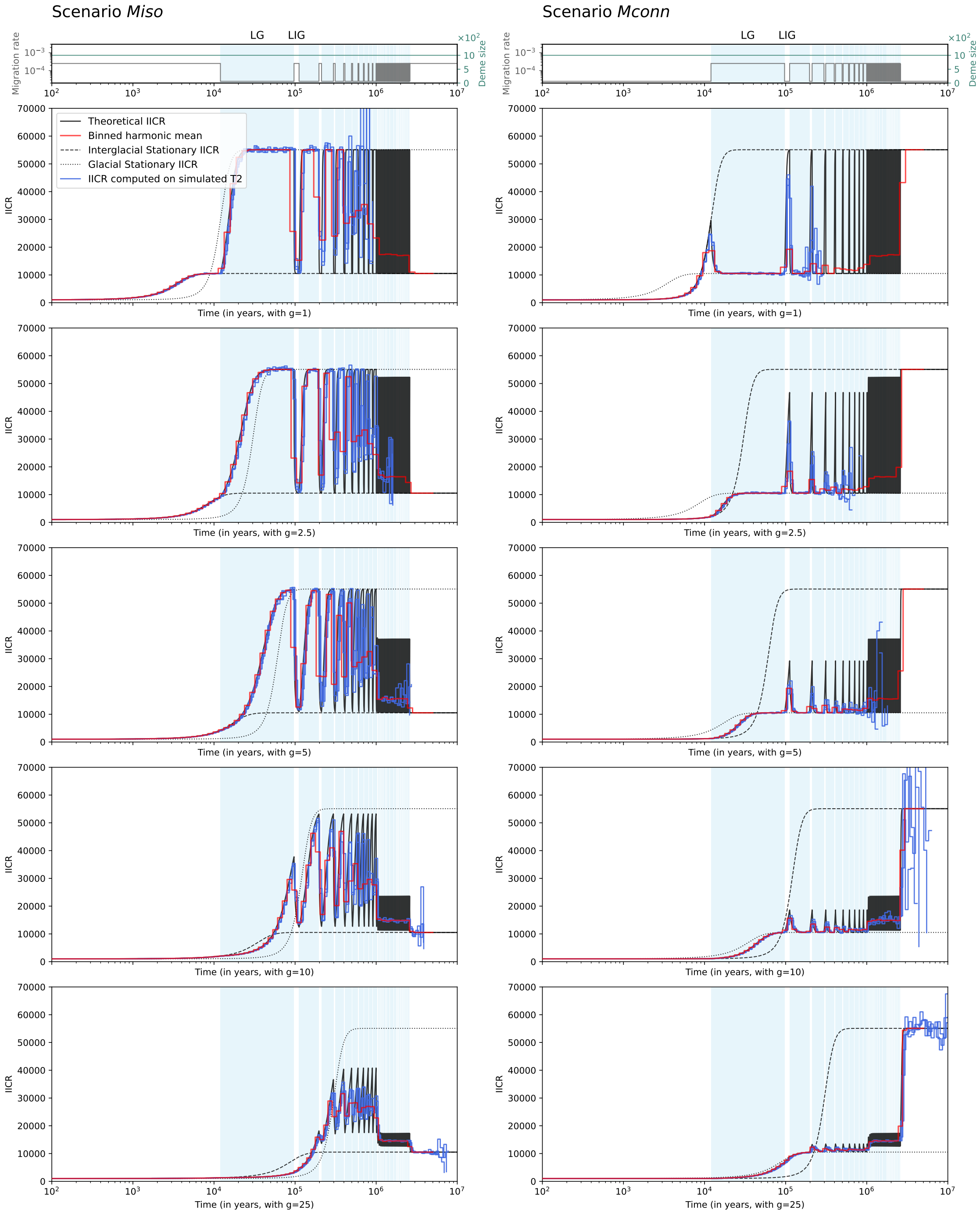


**Figure S8. Simulated IICR (computed from simulated $T_2$) of the scenario $M$ for $m$ varying between** $2.5.10^{-5}$ **and** $2.5.10^{-4}$ **for several generation times.** The left column of panels corresponds to the scenario *Miso* and the right column represents the scenario *Mconn*. Top panels represent migration rates and deme size over time with grey and turquoise lines, respectively. Panels below represent the simulated IICR in blue (three curves computed on three sets of simulated $T_2$), and the theoretical IICR in black in the background, for increasing generation times: 1 (second row), 2.5 (third row), 5 (fourth row), 10 (fifth row) and 25 (sixth row) years. The red curves correspond to the IICR harmonic mean computed over 64 time bins. The pale blue areas in the background correspond to the glacial periods. LG and LIG stand for Last glacial and Last interglacial, respectively.

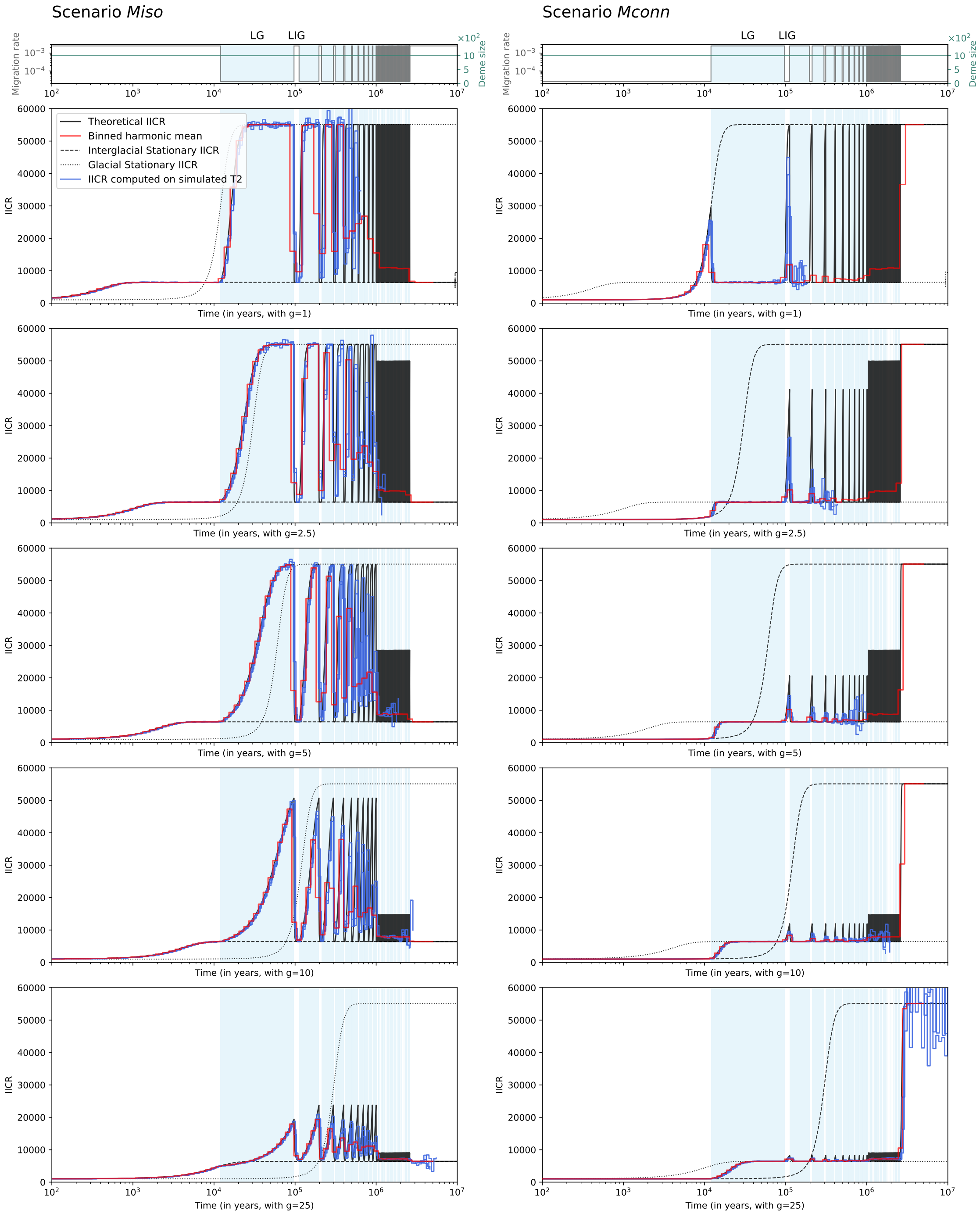


**Figure S9. Simulated IICR (computed from simulated $T_2$) of the scenario $M$ for $m$ varying between** $2.5.10^{-5}$ **and** $2.5.10^{-3}$ **for several generation times.** The left column of panels corresponds to the scenario *Miso* and the right column represents the scenario *Mconn*. Top panels represent migration rates and deme size over time with grey and turquoise lines, respectively. Panels below represent the simulated IICR in blue (three curves computed on three sets of $T_2$), and the theoretical IICR in black in the background, for increasing generation times: 1 (second row), 2.5 (third row), 5 (fourth row), 10 (fifth row) and 25 (sixth row) years. The red curves correspond to the IICR harmonic mean computed over 64 time bins. The pale blue areas in the background correspond to the glacial periods. LG and LIG stand for Last glacial and Last interglacial, respectively.

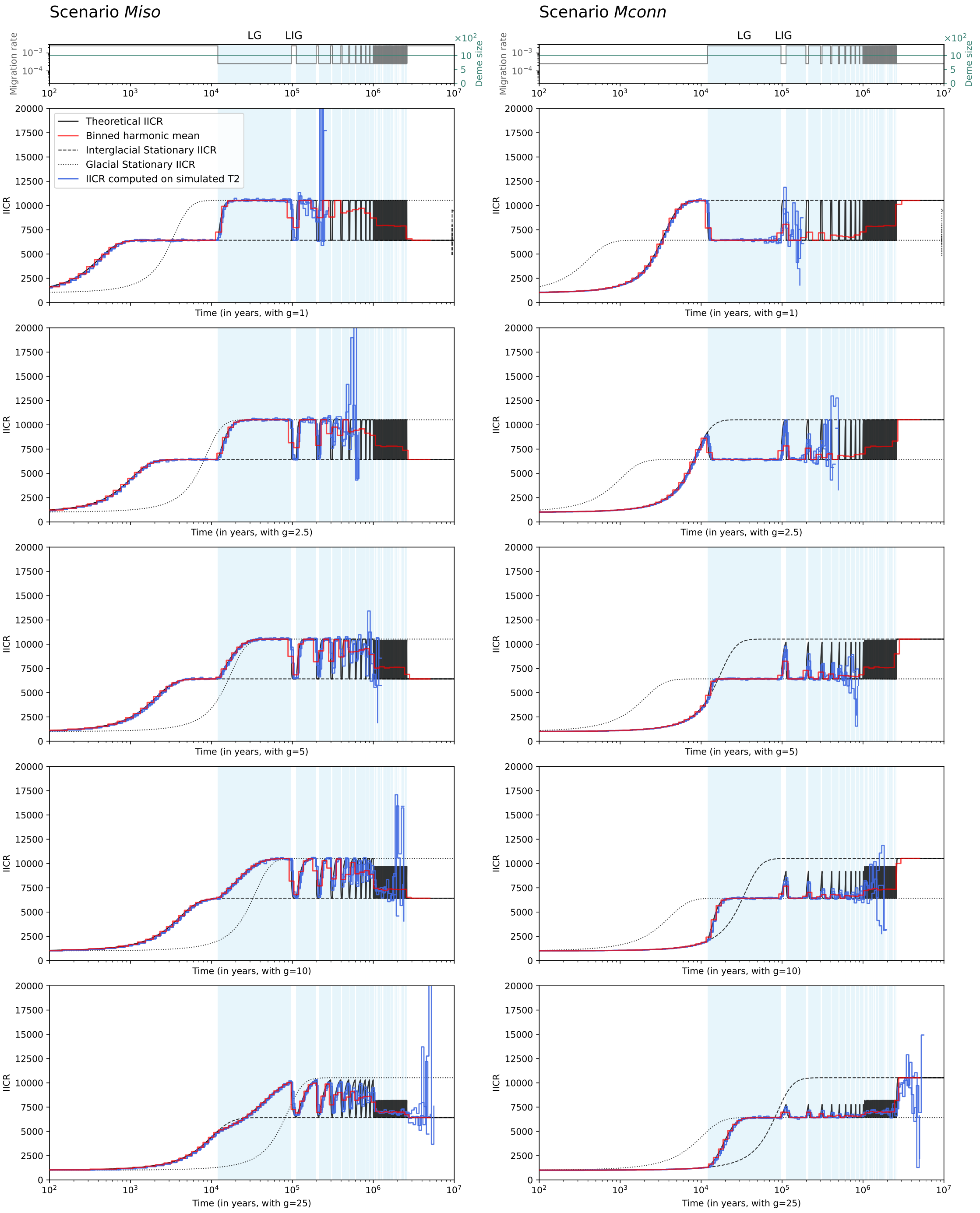


**Figure S10. Simulated IICR (computed from simulated $T_2$) of the scenario *M* for $m$ varying between $2.5.10^{-4}$ and $2.5.10^{-3}$ for several generation times.** The left column of panels corresponds to the scenario *Miso* and the right column represents the scenario *Mconn*. Top panels represent migration rates and deme size over time with grey and turquoise lines, respectively. Panels below represent the simulated IICR in blue (three curves computed on three sets of simulated $T_2$), and the theoretical IICR in black in the background, for increasing generation times: 1 (second row), 2.5 (third row), 5 (fourth row), 10 (fifth row) and 25 (sixth row) years. The red curves correspond to the IICR harmonic mean computed over 64 time bins. The pale blue areas in the background correspond to the glacial periods. LG and LIG stand for Last glacial and Last interglacial, respectively.

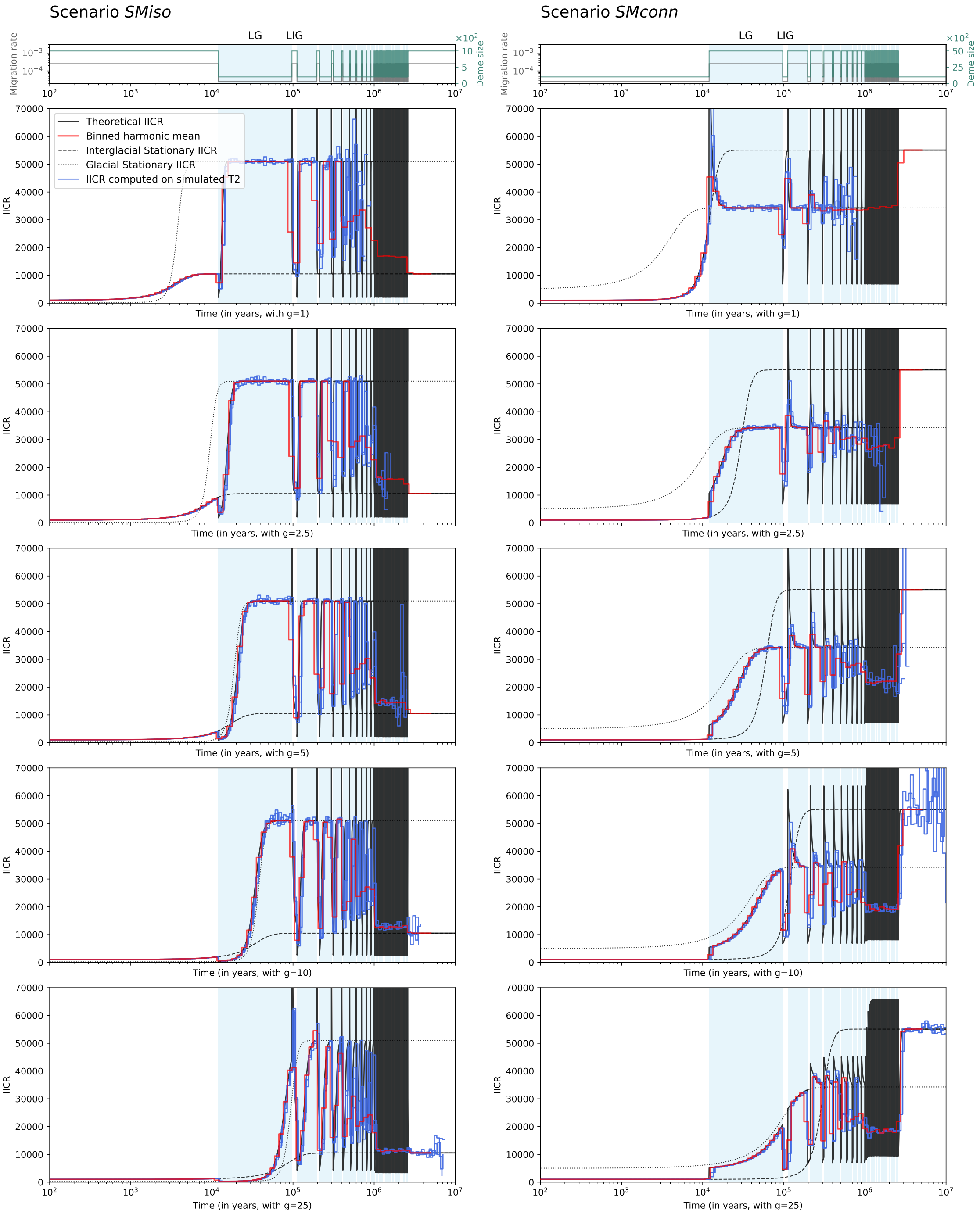


**Figure S11. Simulated IICR (computed from simulated $T_2$) of scenario *SM* for $m$ varying between $2.5.10^{-5}$ and $2.5.10^{-4}$ and $N$ 5-fold increasing or decreasing during glacials, for several generation times.** The left column of panels corresponds to the scenario *SMiso* with $m_{IG} = 2.5.10^{-4}$, $N_{IG} = 1000$ and $m_G = 2.5.10^{-5}$, $N_G = 200$, and the right column represents the scenario *SMconn* with $m_{IG} = 2.5.10^{-5}$, $N_{IG} = 1000$ and $m_G = 2.5.10^{-4}$, $N_G = 5000$. Top panels represent migration rates and deme size over time with grey and turquoise lines, respectively. Panels below represent the simulated IICR in blue (three curves computed on three sets of simulated $T_2$), and the theoretical IICR in black in the background, for increasing generation times: 1 (second row), 2.5 (third row), 5 (fourth row), 10 (fifth row) and 25 (sixth row) years. The red curves correspond to the IICR harmonic mean computed over 64 time bins. The pale blue areas in the background correspond to the glacial periods. LG and LIG stand for Last glacial and Last interglacial, respectively.

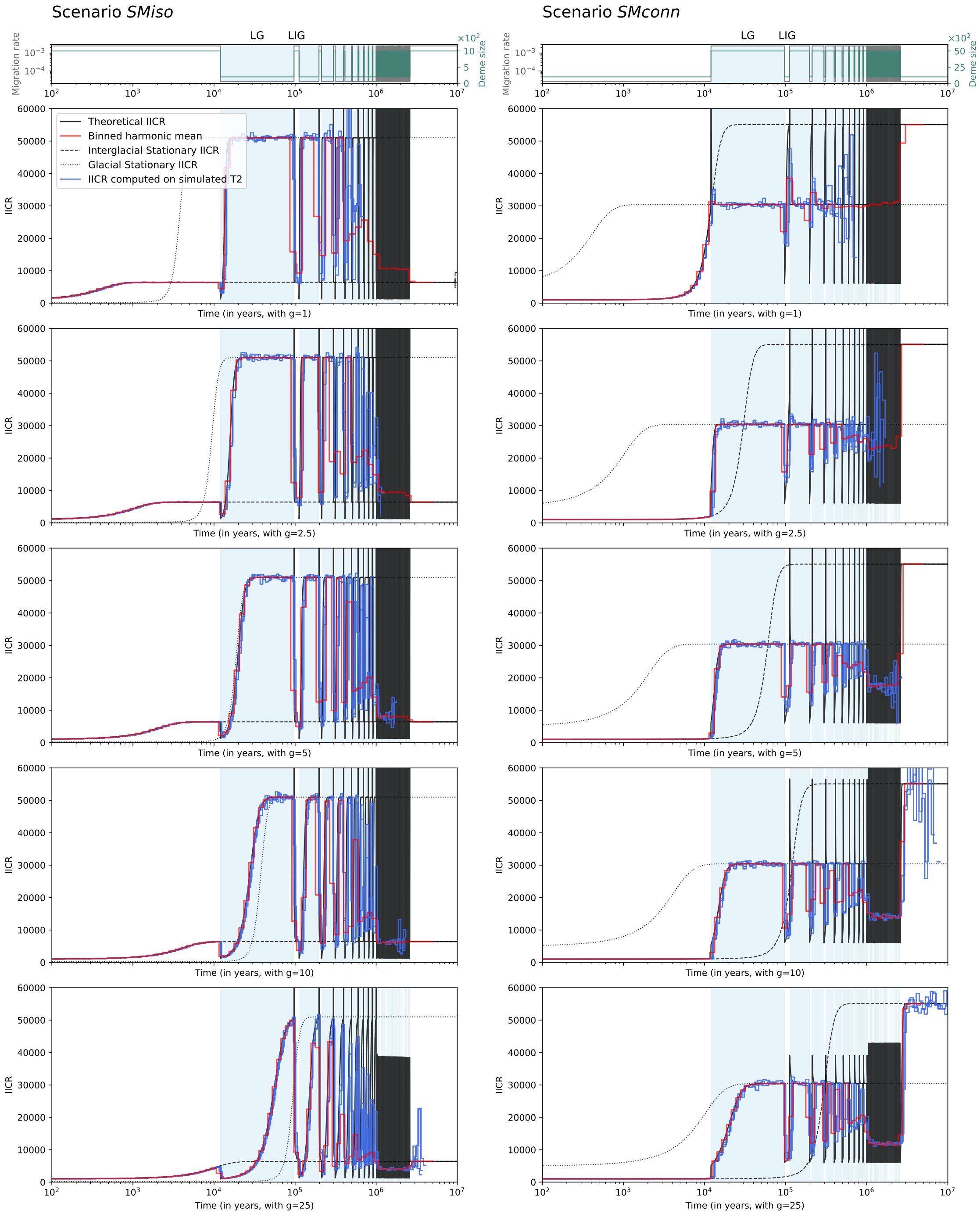


**Figure S12.** **Simulated IICR (computed from simulated $T_2$) of scenario *SM* for $m$ varying between $2.5.10^{-5}$ and $2.5.10^{-3}$ and $N$ 5-fold increasing or decreasing during glacials, for several generation times.** The left column of panels corresponds to the scenario *SMiso* with $m_{IG} = 2.5.10^{-3}$, $N_{IG} = 1000$ and $m_G = 2.5.10^{-5}$, $N_G = 200$, and the right column represents the scenario *SMconn* with $m_{IG} = 2.5.10^{-5}$, $N_{IG} = 1000$ and $m_G = 2.5.10^{-3}$, $N_G = 5000$. Top panels represent migration rates and deme size over time with grey and turquoise lines, respectively. Panels below represent the simulated IICR in blue (three curves computed on three sets of simulated $T_2$), and the theoretical IICR in black in the background, for increasing generation times: 1 (second row), 2.5 (third row), 5 (fourth row), 10 (fifth row) and 25 (sixth row) years. The red curves correspond to the IICR harmonic mean computed over 64 time bins. The pale blue areas in the background correspond to the glacial periods. LG and LIG stand for Last glacial and Last interglacial, respectively.

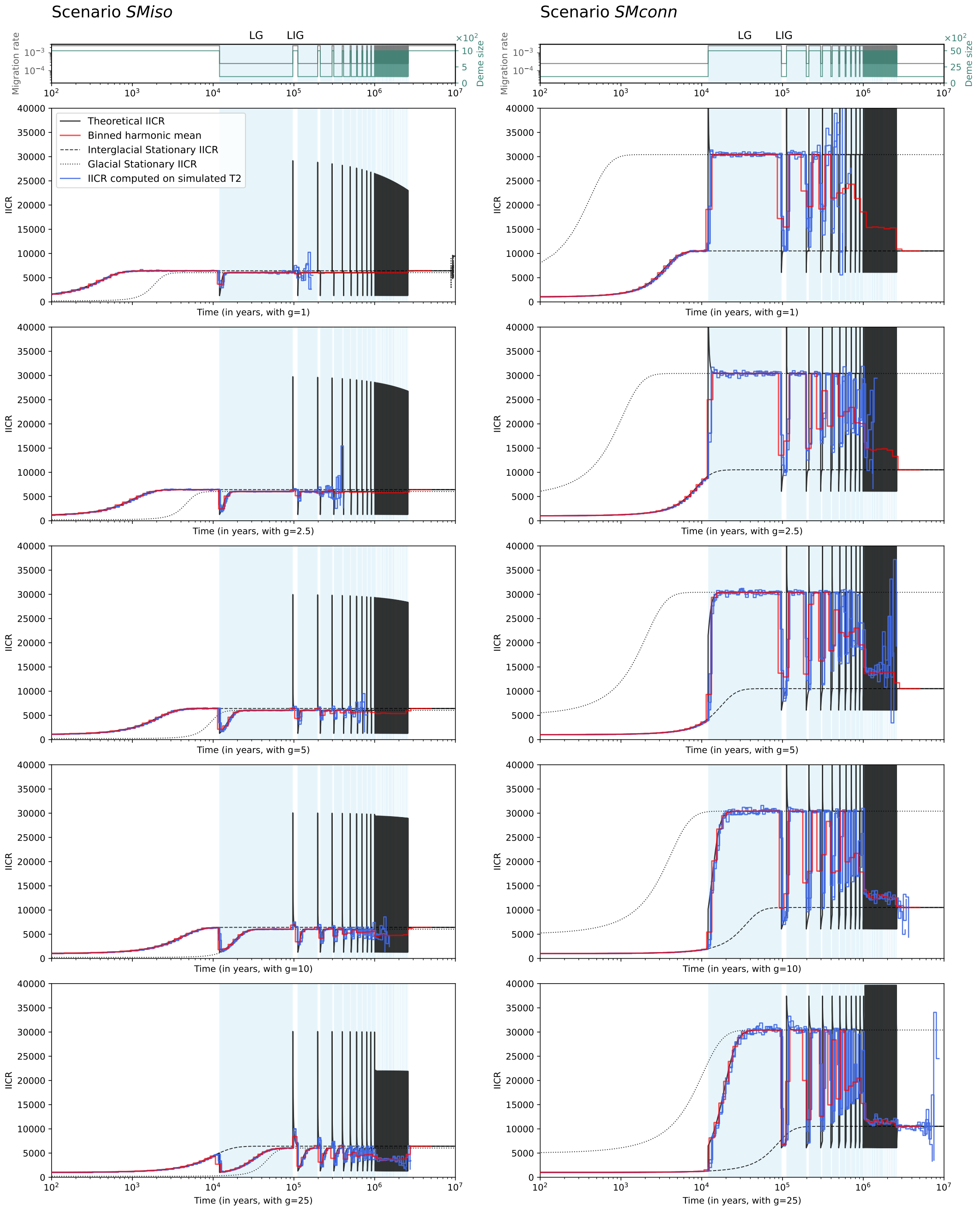


**Figure S13. Simulated IICR (computed from simulated $T_2$) of scenario *SM* for $m$ varying between $2.5.10^{-4}$ and $2.5.10^{-3}$ and $N$ 5-fold increasing or decreasing during glacials, for several generation times.** The left column of panels corresponds to the scenario *SMiso* with $m_{IG} = 2.5.10^{-3}$, $N_{IG} = 1000$ and $m_G = 2.5.10^{-4}$, $N_G = 200$, and the right column represents the scenario *SMconn* with $m_{IG} = 2.5.10^{-4}$, $N_{IG} = 1000$ and $m_G = 2.5.10^{-3}$, $N_G = 5000$. Top panels represent migration rates and deme size over time with grey and turquoise lines, respectively. Panels below represent the simulated IICR in blue (three curves computed on three sets of simulated $T_2$), and the theoretical IICR in black in the background, for increasing generation times: 1 (second row), 2.5 (third row), 5 (fourth row), 10 (fifth row) and 25 (sixth row) years. The red curves correspond to the IICR harmonic mean computed over 64 time bins. The pale blue areas in the background correspond to the glacial periods. LG and LIG stand for Last glacial and Last interglacial, respectively.

### PSMC curves

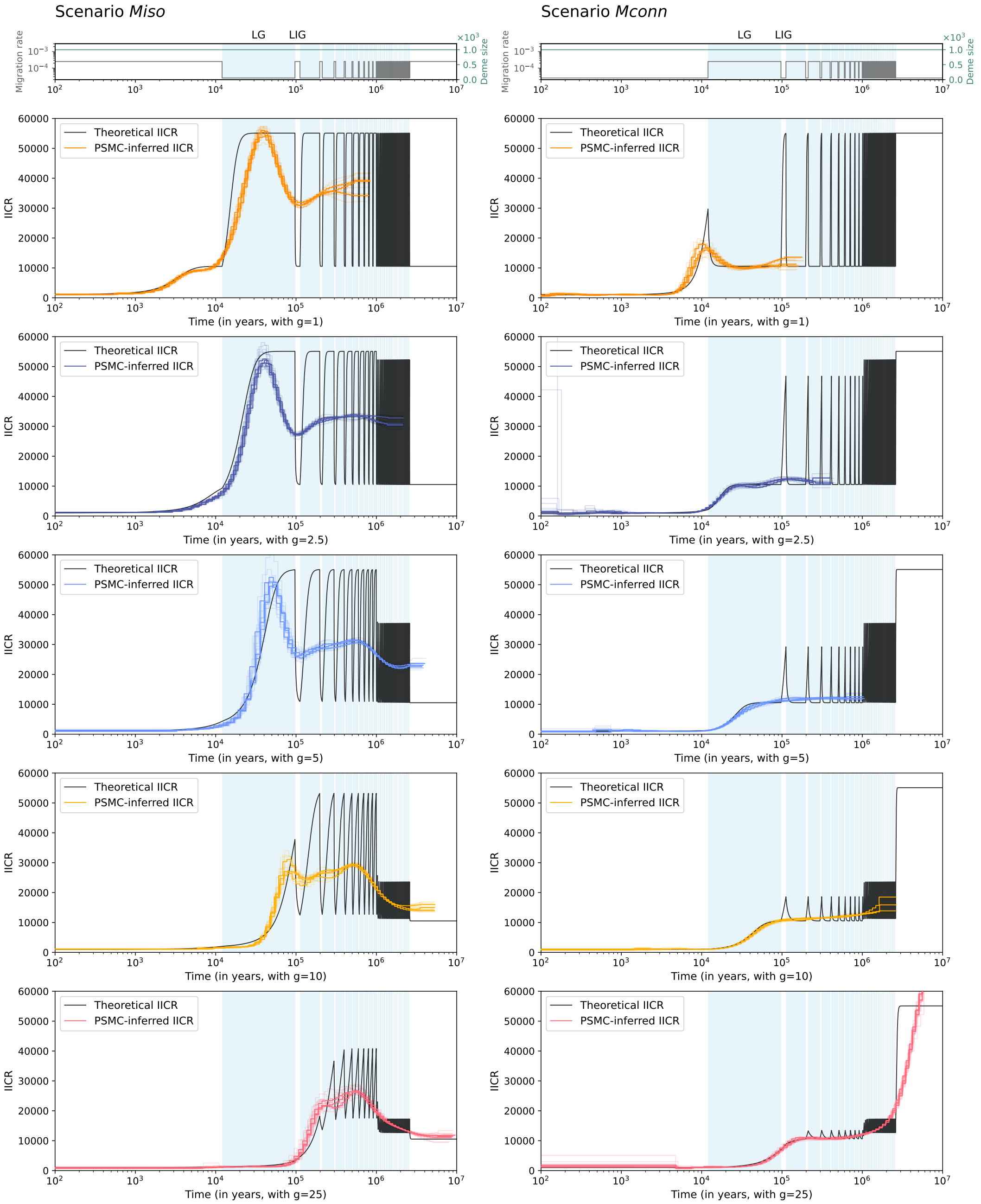


**Figure S14. PSMC curves of scenarios *M* for *m* varying between** $2.5.10^{-5}$ **and** $2.5.10^{-4}$ **for several generation times.** The left column of panels corresponds to the scenario *Miso*, and the right column represents the scenario *Mconn*. Top panels represent migration rates and deme sizes over time with grey and turquoise lines, respectively. Panels below represent the inferred PSMC curves and the theoretical IICR in black in the background, for increasing generation times: 1 (second row), 2.5 (third row), 5 (fourth row), 10 (fifth row) and 25 (sixth row) years. Each coloured curve corresponds to the PSMC curve of one simulated diploid genome sampled in the present and in the same deme (3 simulated genomes in total), and the lighter coloured curves correspond to PSMC bootstraps (5 bootstraps per PSMC curve). The pale blue areas in the background correspond to glacial periods. LG and LIG stand for Last glacial and Last interglacial, respectively.

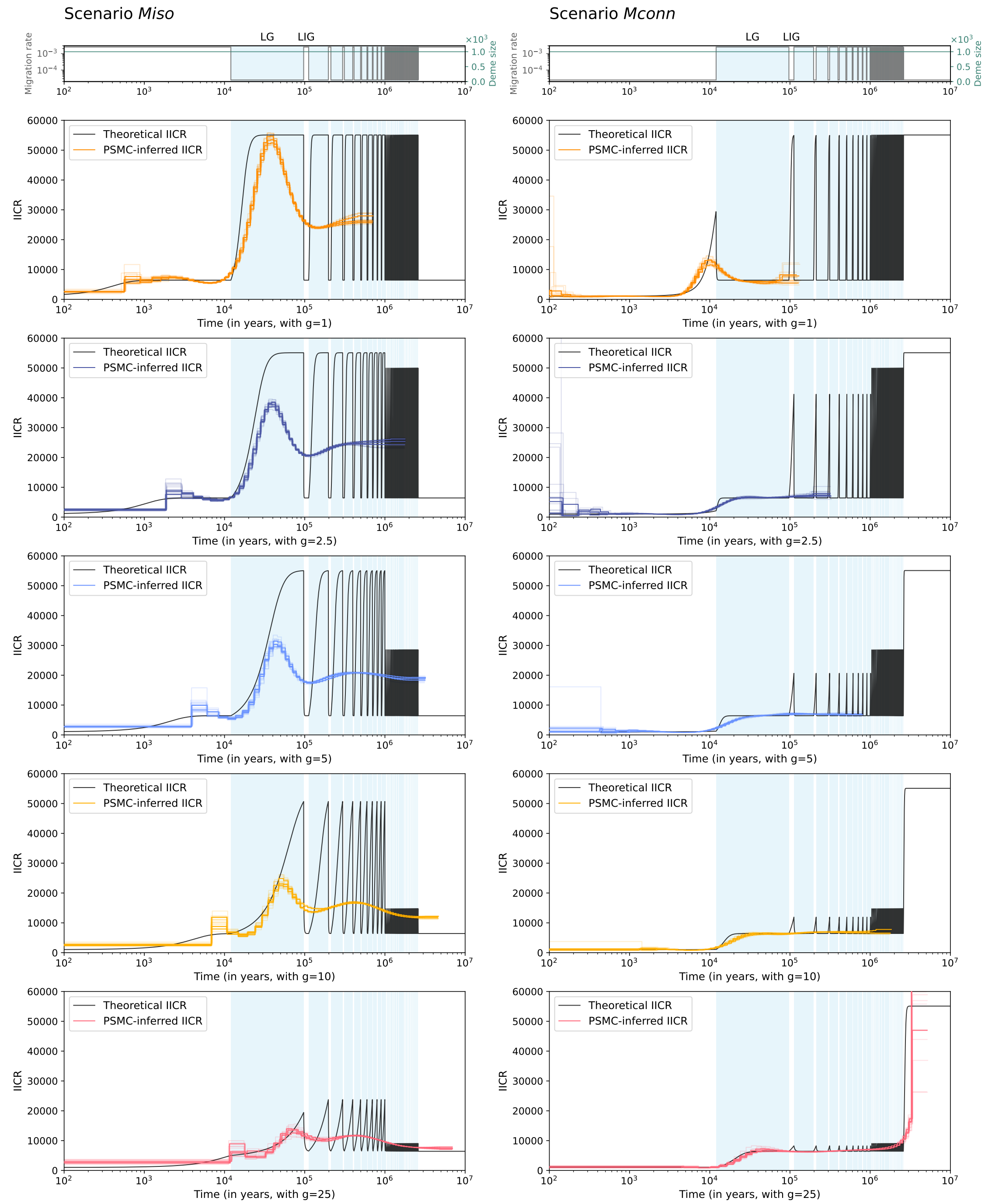


**Figure S15.** **PSMC curves of scenarios *M* for $m$ varying between $2.5.10^{-5}$ and $2.5.10^{-3}$ for several generation times.** The left column of panels corresponds to the scenario *Miso*, and the right column represents the scenario *Mconn*. Top panels represent migration rates and deme sizes over time with grey and turquoise lines, respectively. Panels below represent the inferred PSMC curves and the theoretical IICR in black in the background, for increasing generation times: 1 (second row), 2.5 (third row), 5 (fourth row), 10 (fifth row) and 25 (sixth row) years. Each coloured curve corresponds to the PSMC curve of one simulated diploid genome sampled in the present and in the same deme (3 simulated genomes in total), and the lighter coloured curves correspond to PSMC bootstraps (5 bootstraps per PSMC curve). The pale blue areas in the background correspond to glacial periods. LG and LIG stand for Last glacial and Last interglacial, respectively.

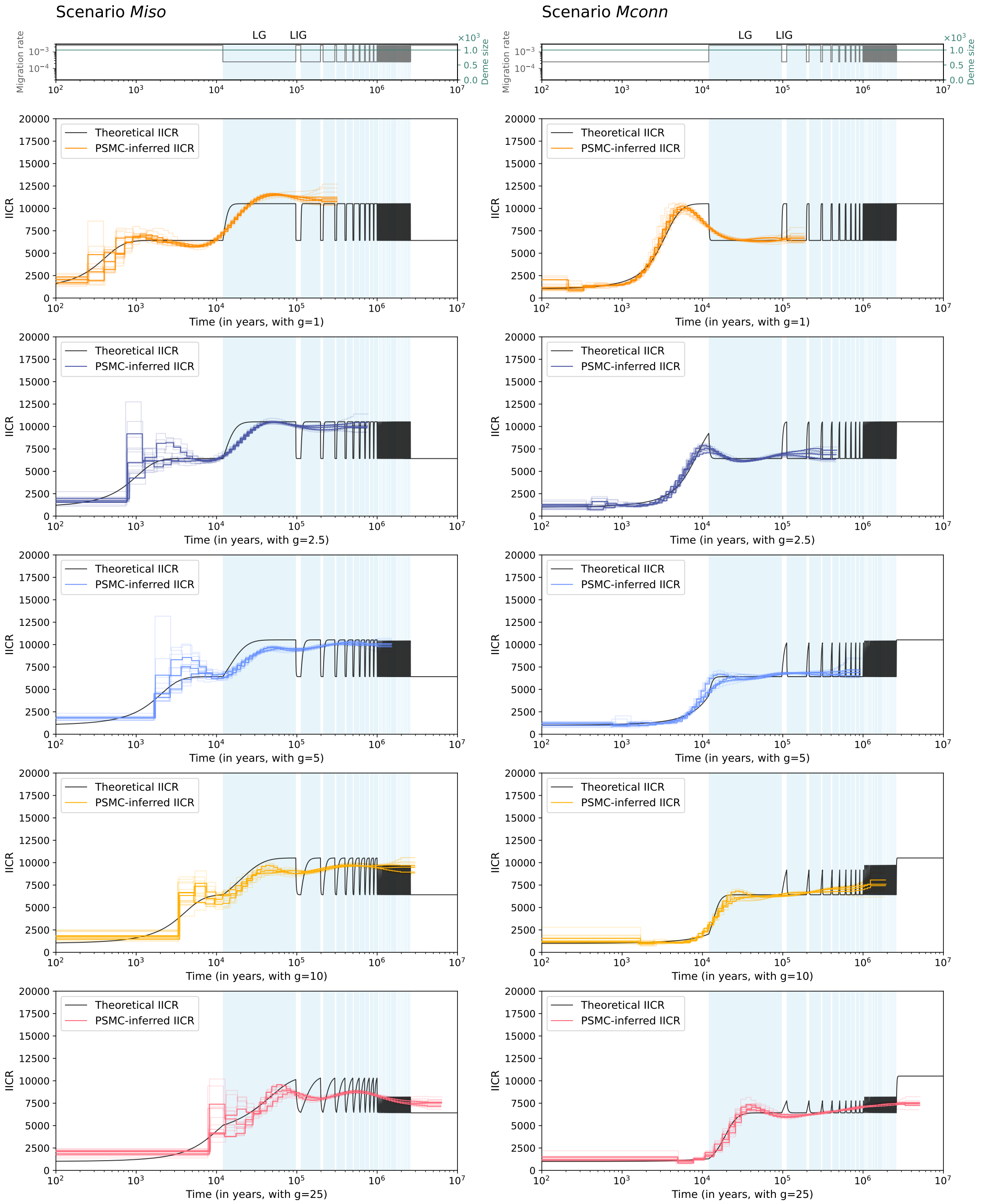


**Figure S16.** **PSMC curves of scenarios *M* for *m* varying between** $2.5.10^{-4}$ **and** $2.5.10^{-3}$ **for several generation times.** The left column of panels corresponds to the scenario *Miso*, and the right column represents the scenario *Mconn*. Top panels represent migration rates and deme sizes over time with grey and turquoise lines, respectively. Panels below represent the inferred PSMC curves and the theoretical IICR in black in the background, for increasing generation times: 1 (second row), 2.5 (third row), 5 (fourth row), 10 (fifth row) and 25 (sixth row) years. Each coloured curve corresponds to the PSMC curve of one simulated diploid genome sampled in the present and in the same deme (3 simulated genomes in total), and the lighter coloured curves correspond to PSMC bootstraps (5 bootstraps per PSMC curve). The pale blue areas in the background correspond to glacial periods. LG and LIG stand for Last glacial and Last interglacial, respectively.

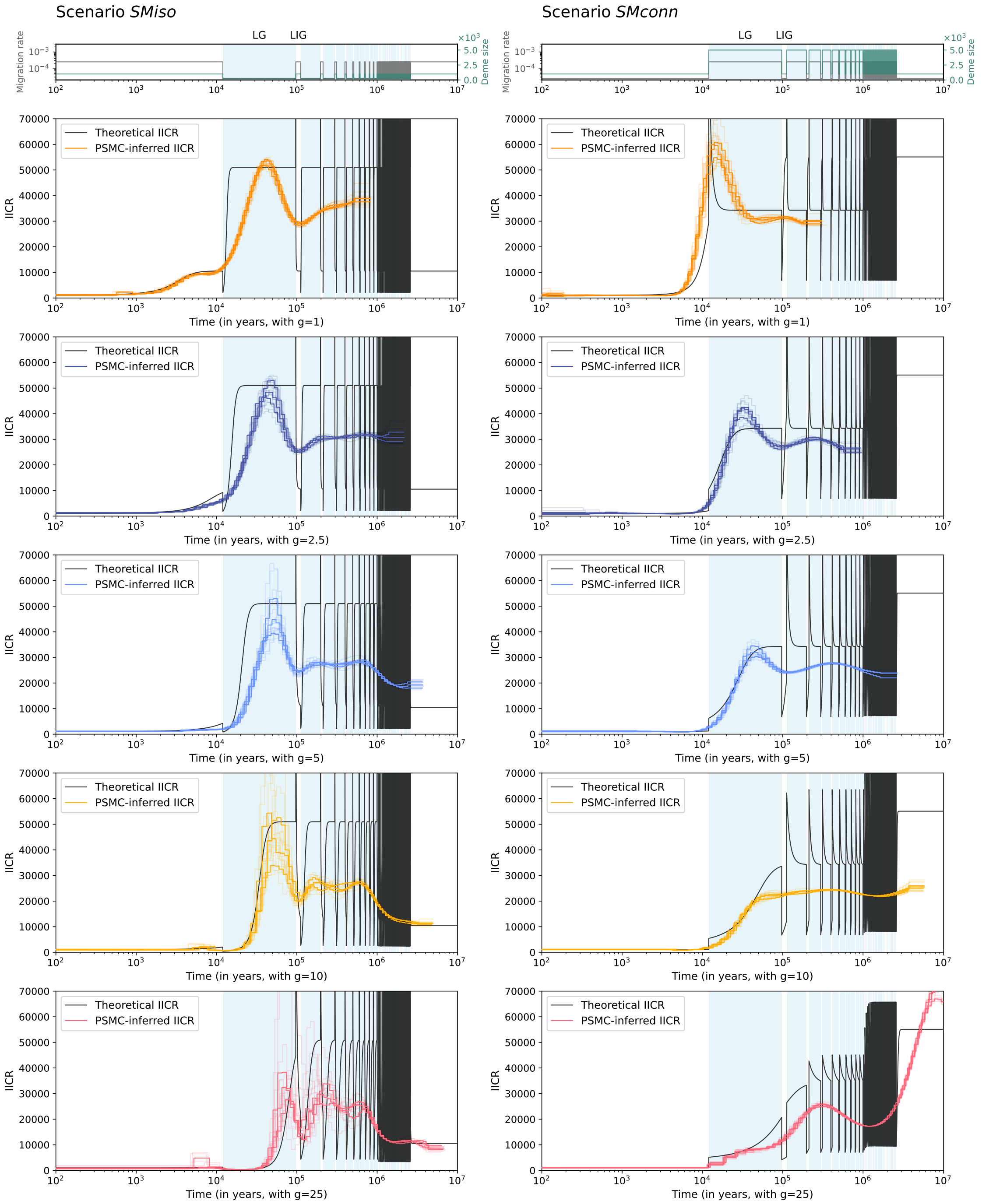


**Figure S17.** **PSMC curves of scenarios *SM* for $m$ varying between $2.5.10^{-5}$ and $2.5.10^{-4}$ and $N$ 5-fold increasing or decreasing during glacials, for several generation times.** Left column of panels corresponds to scenario *SMiso* with $m_{IG} = 2.5.10^{-4}$, $N_{IG} = 1000$ and $m_G = 2.5.10^{-5}$, $N_G = 200$, and right column represents scenario *SMconn* with $m_{IG} = 2.5.10^{-5}$, $N_{IG} = 1000$ and $m_G = 2.5.10^{-4}$, $N_G = 5000$. Top panels represent migration rates and deme sizes over time with grey and turquoise lines, respectively. Panels below represent the inferred PSMC curves and the theoretical IICR in black in the background, for increasing generation times: 1 (second row), 2.5 (third row), 5 (fourth row), 10 (fifth row) and 25 (sixth row) years. Each coloured curve corresponds to the PSMC curve of one simulated diploid genome sampled in the present and in same deme (3 simulated genomes in total), and the lighter coloured curves correspond to PSMC bootstraps (5 bootstraps per PSMC curve). Pale blue areas in the background correspond to glacial periods. LG and LIG stand for Last glacial and Last interglacial, respectively.

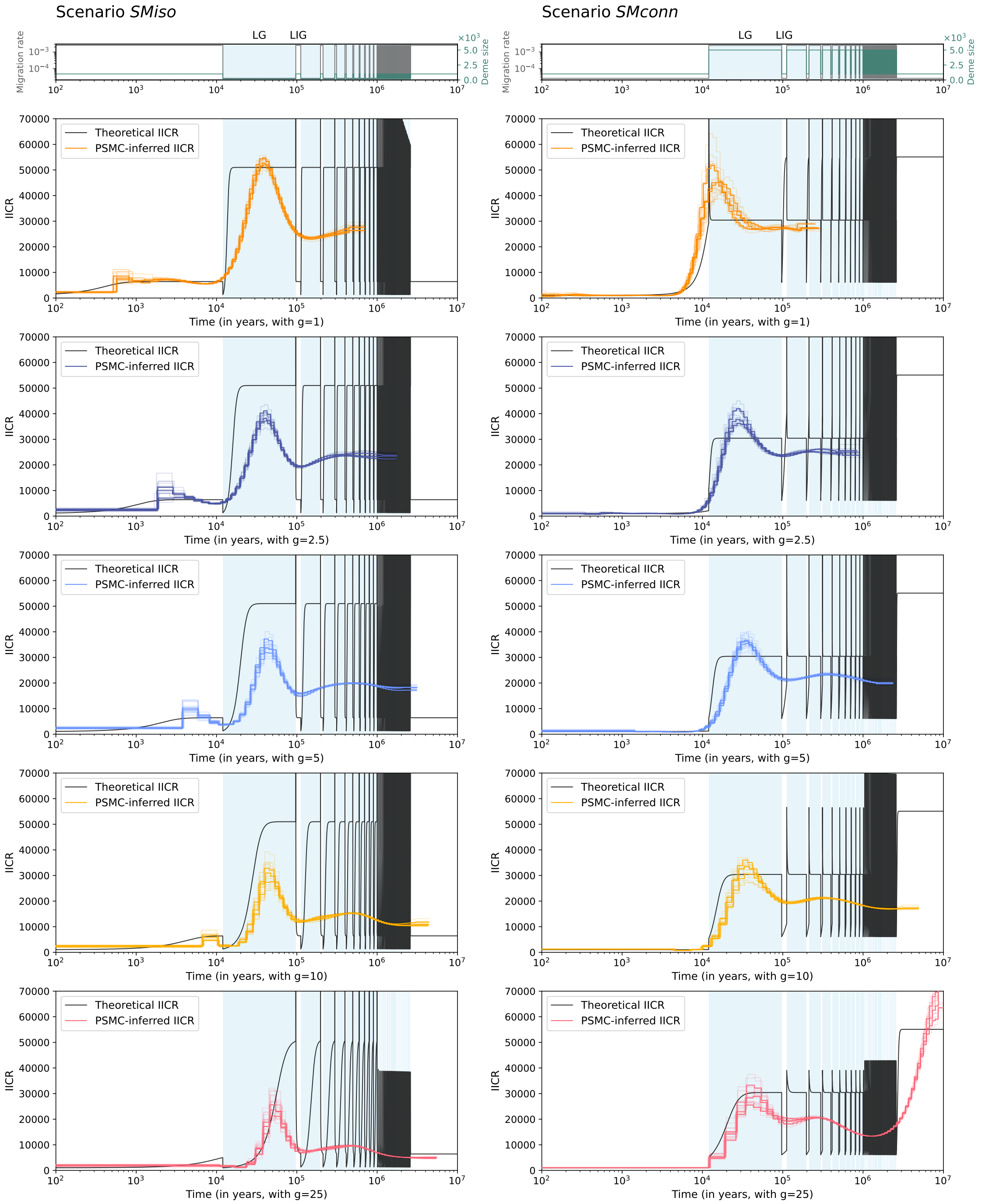


**Figure S18. PSMC curves of scenarios *SM* for $m$ varying between $2.5.10^{-5}$ and $2.5.10^{-3}$ and $N$ 5-fold increasing or decreasing during glacials, for several generation times.** Left column of panels corresponds to scenario *SMiso* with $m_{IG} = 2.5.10^{-3}$, $N_{IG} = 1000$ and $m_G = 2.5.10^{-5}$, $N_G = 200$, and right column represents scenario *SMconn* with $m_{IG} = 2.5.10^{-5}$, $N_{IG} = 1000$ and $m_G = 2.5.10^{-3}$, $N_G = 5000$. Top panels represent migration rates and deme sizes over time with grey and turquoise lines, respectively. Panels below represent the inferred PSMC curves and the theoretical IICR in black in the background, for increasing generation times: 1 (second row), 2.5 (third row), 5 (fourth row), 10 (fifth row) and 25 (sixth row) years. Each coloured curve corresponds to the PSMC curve of one simulated diploid genome sampled in the present and in the same deme (3 simulated genomes in total), and the lighter coloured curves correspond to PSMC bootstraps (5 bootstraps per PSMC curve). Pale blue areas in the background correspond to glacial periods. LG and LIG stand for Last glacial and Last interglacial, respectively.

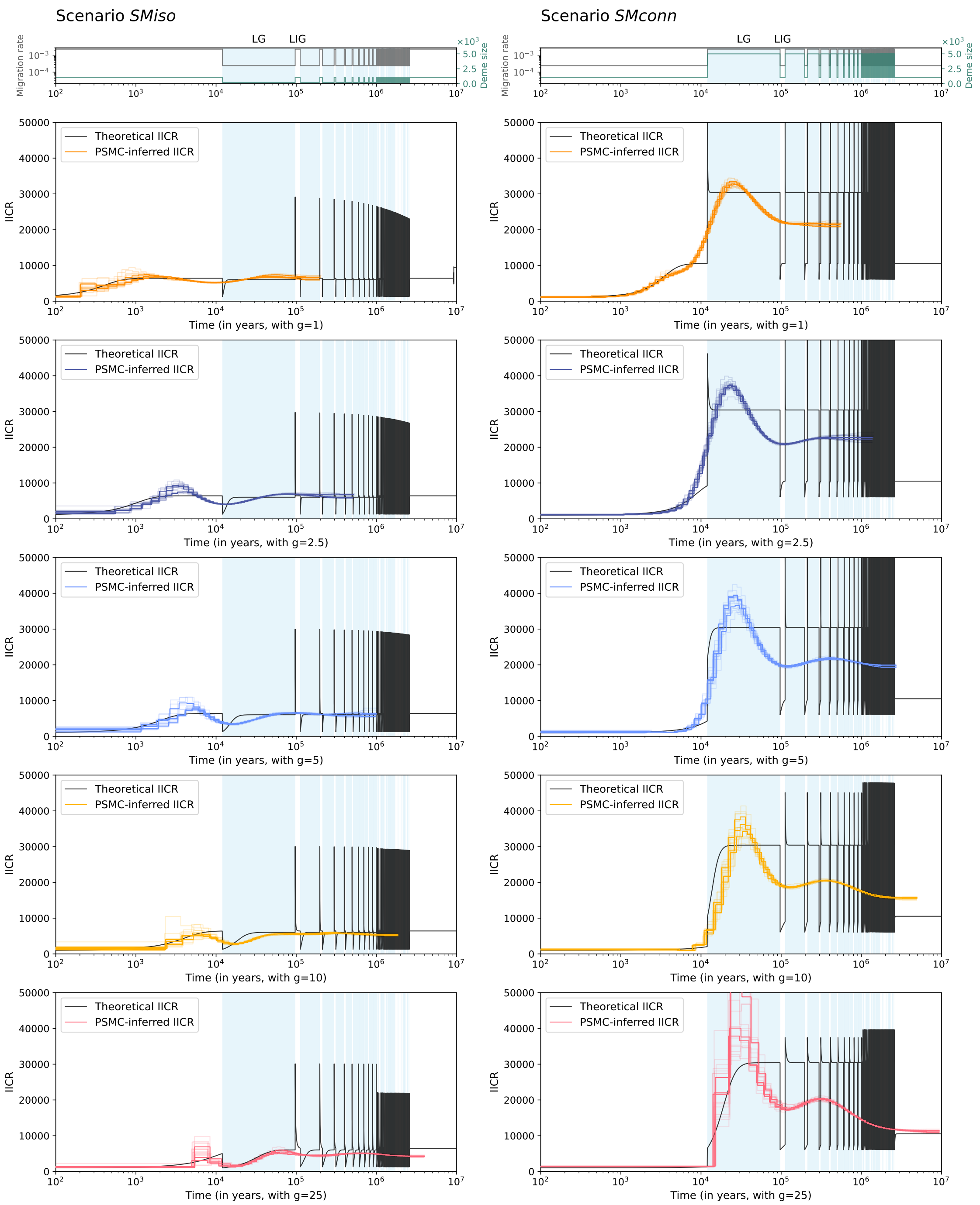


**Figure S19. PSMC curves of scenarios *SM* for $m$ varying between $2.5.10^{-4}$ and $2.5.10^{-3}$ and $N$ 5-fold increasing or decreasing during glacials, for several generation times.** Left column of panels corresponds to scenario *SMiso* with $m_{IG} = 2.5.10^{-3}$, $N_{IG} = 1000$ and $m_G = 2.5.10^{-4}$, $N_G = 200$, and right column represents scenario *SMconn* with $m_{IG} = 2.5.10^{-4}$, $N_{IG} = 1000$ and $m_G = 2.5.10^{-3}$, $N_G = 5000$. Top panels represent migration rates and deme sizes over time with grey and turquoise lines, respectively. Panels below represent the inferred PSMC curves and the theoretical IICR in black in the background, for increasing generation times: 1 (second row), 2.5 (third row), 5 (fourth row), 10 (fifth row) and 25 (sixth row) years. Each coloured curve corresponds to the PSMC curve of one simulated diploid genome sampled in the present and in the same deme (3 simulated genomes in total), and the lighter coloured curves correspond to PSMC bootstraps (5 bootstraps per PSMC curve). Pale blue areas in the background correspond to glacial periods. LG and LIG stand for Last glacial and Last interglacial, respectively.

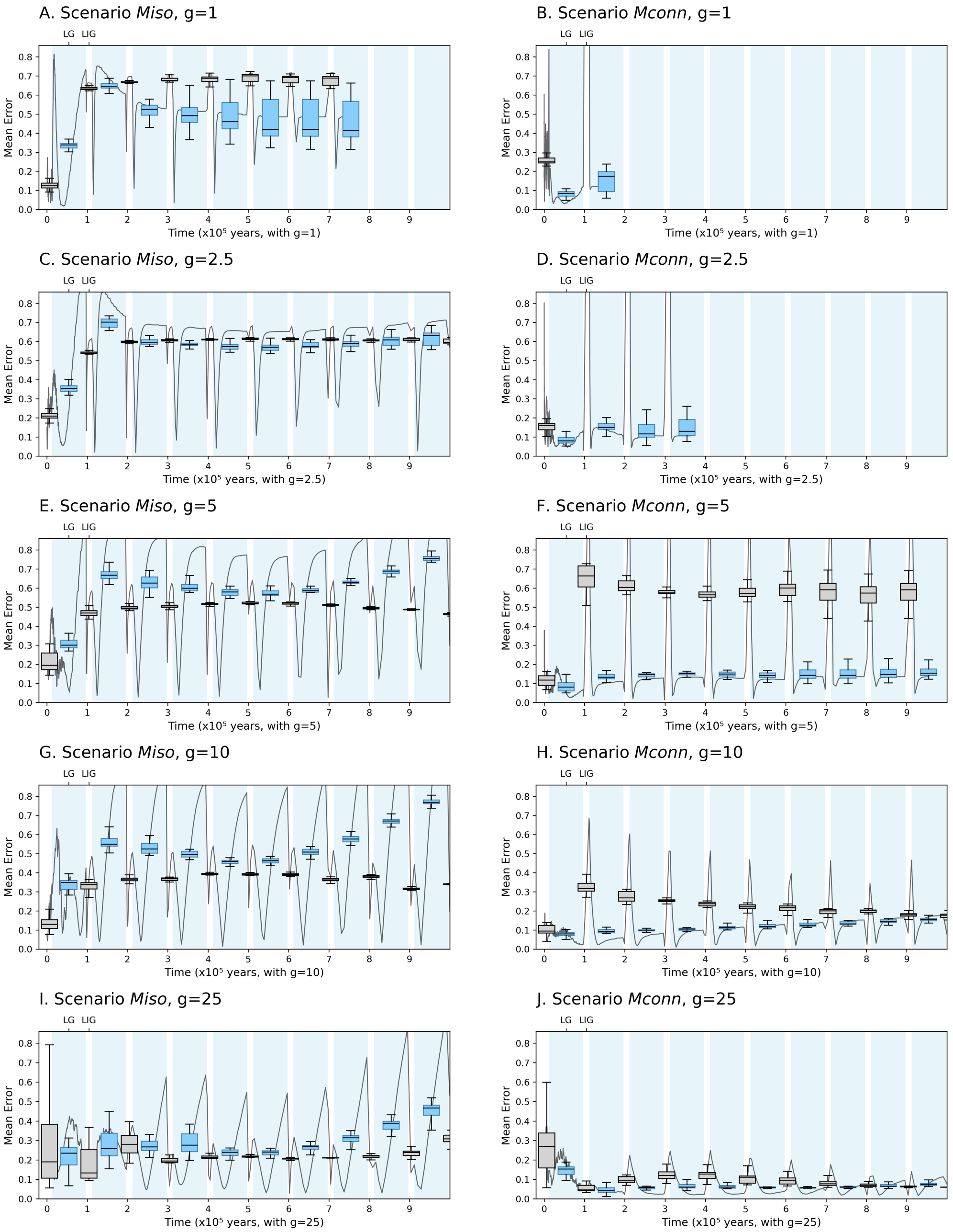


**Figure S20. Mean relative error (RE) of PSMC estimation of the theoretical IICR across time for scenarios *M* with** $m$ **varying between** $2.5.10^{-5}$ **and** $2.5.10^{-4}$ **for several generation times.** A: *Miso* for g = 1 year, B: *Mconn* for g = 1 year, C: *Miso* for g = 2.5 years, D: *Mconn* for g = 2.5 years, E: *Miso* for g = 5 years, F: *Mconn* for g = 5 years, G: *Miso* for g = 10 years, H: *Mconn* for g = 10 years, I: *Miso* for g = 25 years, and J: *Mconn* for g = 25 years. The RE was defined as $\frac{|IICR_t - PSMC_t|}{IICR_t}$ and its mean was computed across the three PSMC curves and 15 bootstraps. In each sub-figure, the box-plots correspond to a mean RE computed every 99,9 years (10,000 time points) and averaged for each interglacial and interglacial period (1-12 kya, 12-97 kya, 97-112 kya and so on, up to 1 Mya). The grey curve corresponds to a mean RE computed every 999 years (1,000 time point in total between 1 kya and 1 Mya). Light blue rectangles in the background correspond to glacial periods.

Please note that, due to the cyclic nature of our scenarios and the oscillations of the IICR, the mean RE across time is not necessarily informative about the ability of PSMC to infer the demographic oscillations (see Material and Methods and Supp. Information S1).

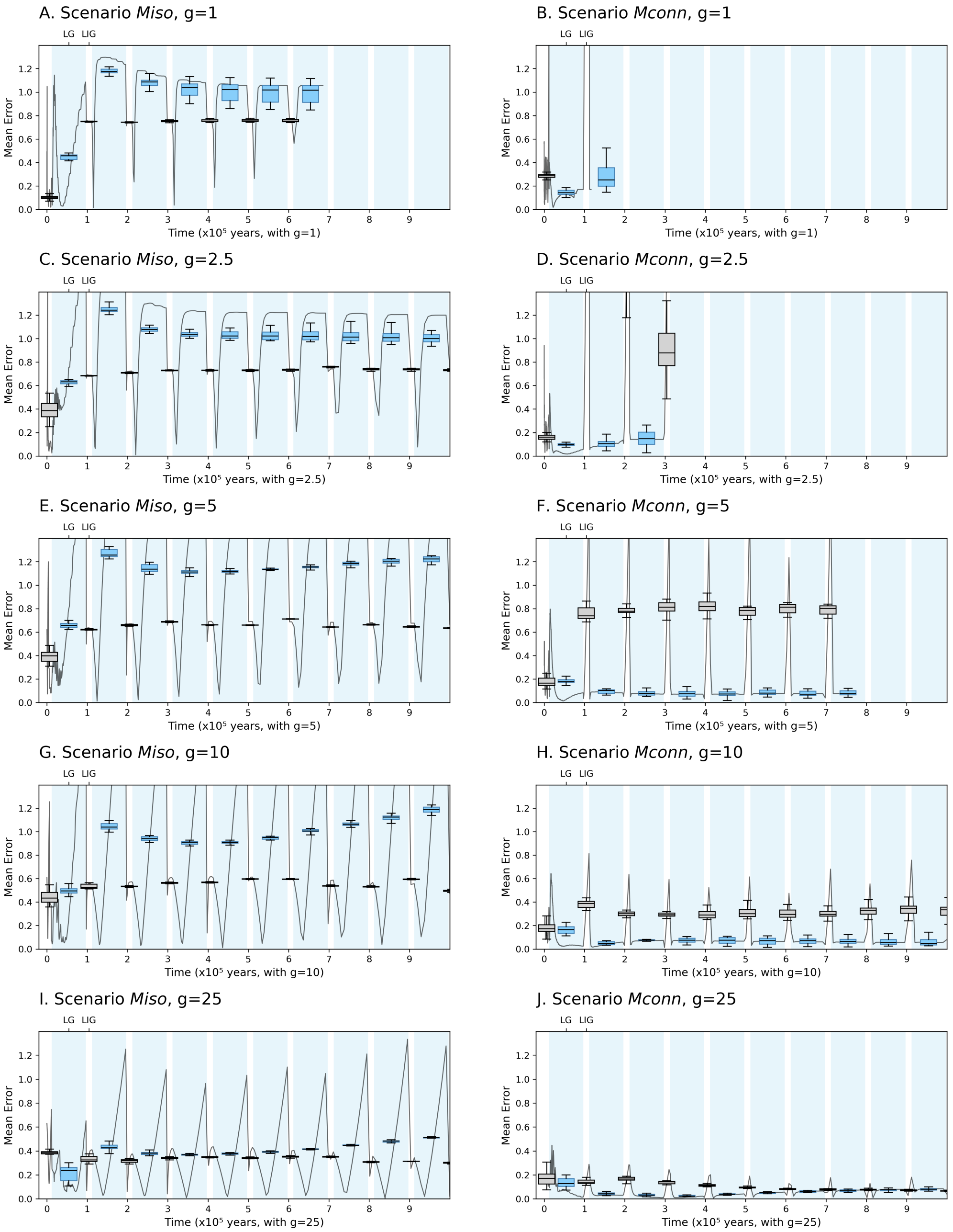


**Figure S21.** **Mean error (ME) of PSMC estimation of the theoretical IICR across time for scenarios *M* with $m$ varying between** $2.5.10^{-5}$ **and** $2.5.10^{-3}$ **for several generation times.** A: *Miso* for g = 1 year, B: *Mconn* for g = 1 year, C: *Miso* for g = 2.5 years, D: *Mconn* for g = 2.5 years, E: *Miso* for g = 5 years, F: *Mconn* for g = 5 years, G: *Miso* for g = 10 years, H: *Mconn* for g = 10 years, I: *Miso* for g = 25 years, and J: *Mconn* for g = 25 years. The RE was defined as $\frac{|IICR_t - PSMC_t|}{IICR_t}$ and its mean was computed across the three PSMC curves and 15 bootstraps. In each sub-figure, the box-plots correspond to a mean RE computed every 99,9 years (10,000 time points) and averaged for each interglacial and interglacial period (1-12 kya, 12-97 kya, 97-112 kya and so on, up to 1 Mya). The grey curve corresponds to a mean RE computed every 999 years (1,000 time point in total between 1 kya and 1 Mya). Light blue rectangles in the background correspond to glacial periods.
Please note that, due to the cyclic nature of our scenarios and the oscillations of the IICR, the mean RE across time is not necessarily informative about the ability of PSMC to infer the demographic oscillations (see Material and Methods and Supp. Information S1).

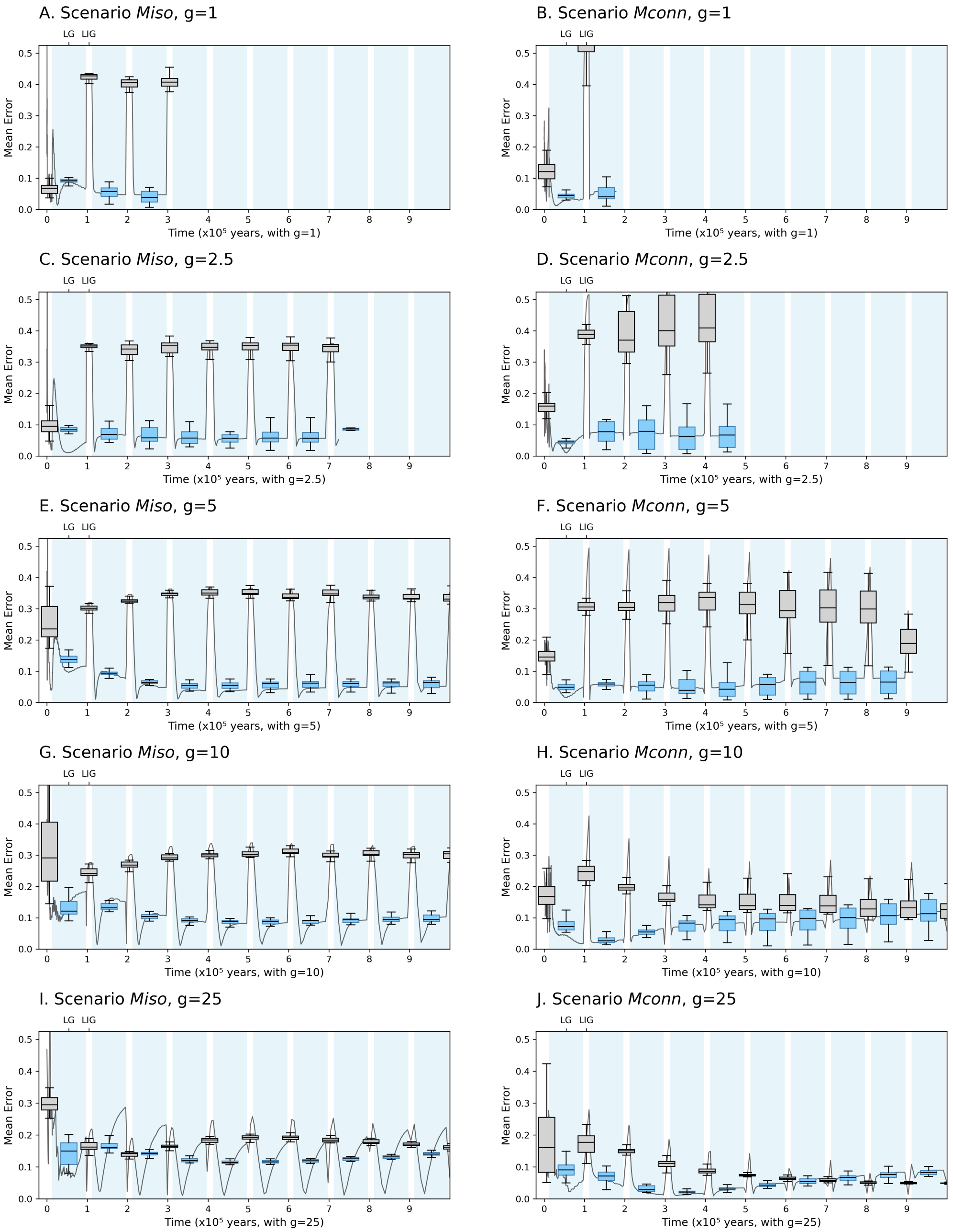


**Figure S22. Mean error (ME) of PSMC estimation of the theoretical IICR across time for scenarios *M* with *m* varying between** $2.5.10^{-4}$ **and** $2.5.10^{-3}$ **for several generation times.** A: *Miso* for g = 1 year, B: *Mconn* for g = 1 year, C: *Miso* for g = 2.5 years, D: *Mconn* for g = 2.5 years, E: *Miso* for g = 5 years, F: *Mconn* for g = 5 years, G: *Miso* for g = 10 years, H: *Mconn* for g = 10 years, I: *Miso* for g = 25 years, and J: *Mconn* for g = 25 years. The RE was defined as $\frac{|IICR_t - PSMC_t|}{IICR_t}$ and its mean was computed across the three PSMC curves and 15 bootstraps. In each sub-figure, the box-plots correspond to a mean RE computed every 99,9 years (10,000 time points) and averaged for each interglacial and interglacial period (1-12 kya, 12-97 kya, 97-112 kya and so on, up to 1 Mya). The grey curve corresponds to a mean RE computed every 999 years (1,000 time point in total between 1 kya and 1 Mya). Light blue rectangles in the background correspond to glacial periods.

Please note that, due to the cyclic nature of our scenarios and the oscillations of the IICR, the mean RE across time is not necessarily informative about the ability of PSMC to infer the demographic oscillations (see Material and Methods and Supp. Information S1).

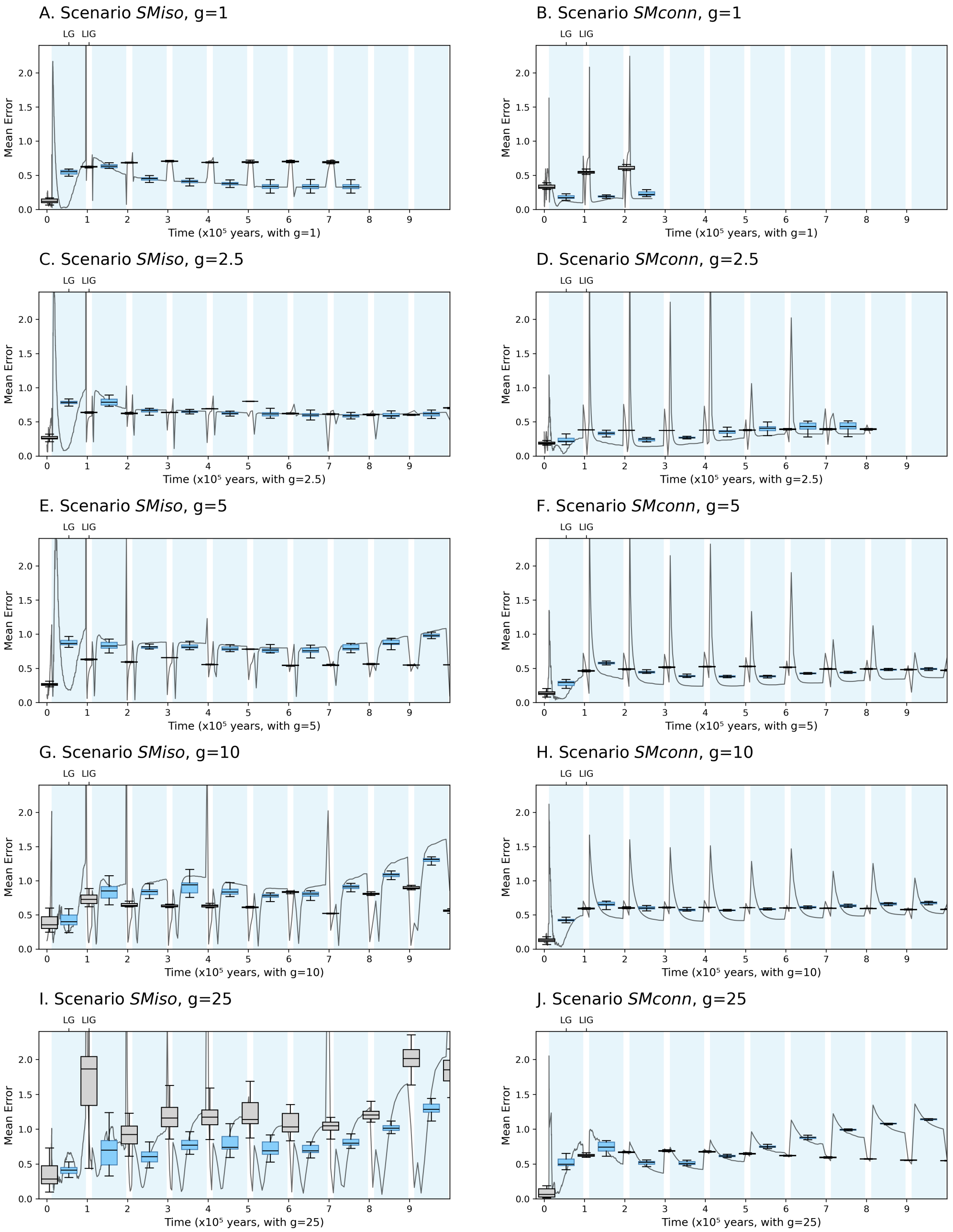


**Figure S23.** **Mean error (ME) of PSMC estimation of the theoretical IICR across time for scenarios *SM* with $m$ varying between** $2.5.10^{-5}$ **and** $2.5.10^{-4}$ **and** $N$ **5-fold increasing or decreasing during glacials for several generation times.** A: *SMiso* for g = 1 year, B: *SMconn* for g = 1 year, C: *SMiso* for g = 2.5 years, D: *SMconn* for g = 2.5 years, E: *SMiso* for g = 5 years, F: *SMconn* for g = 5 years, G: *SMiso* for g = 10 years, H: *SMconn* for g = 10 years, I: *SMiso* for g = 25 years, and J: *SMconn* for g = 25 years. The RE was defined as $\frac{|IICR_t - PSMC_t|}{IICR_t}$ and its mean was computed across the three PSMC curves and 15 bootstraps. In each sub-figure, the box-plots correspond to a mean RE computed every 99,9 years (10,000 time points) and averaged for each interglacial and interglacial period (1-12 kya, 12-97 kya, 97-112 kya and so on, up to 1 Mya). The grey curve corresponds to a mean RE computed every 999 years (1,000 time point in total between 1 kya and 1 Mya). Light blue rectangles in the background correspond to glacial periods.

Please note that, due to the cyclic nature of our scenarios and the oscillations of the IICR, the mean RE across time is not necessarily informative about the ability of PSMC to infer the demographic oscillations (see Material and Methods and Supp. Information S1).

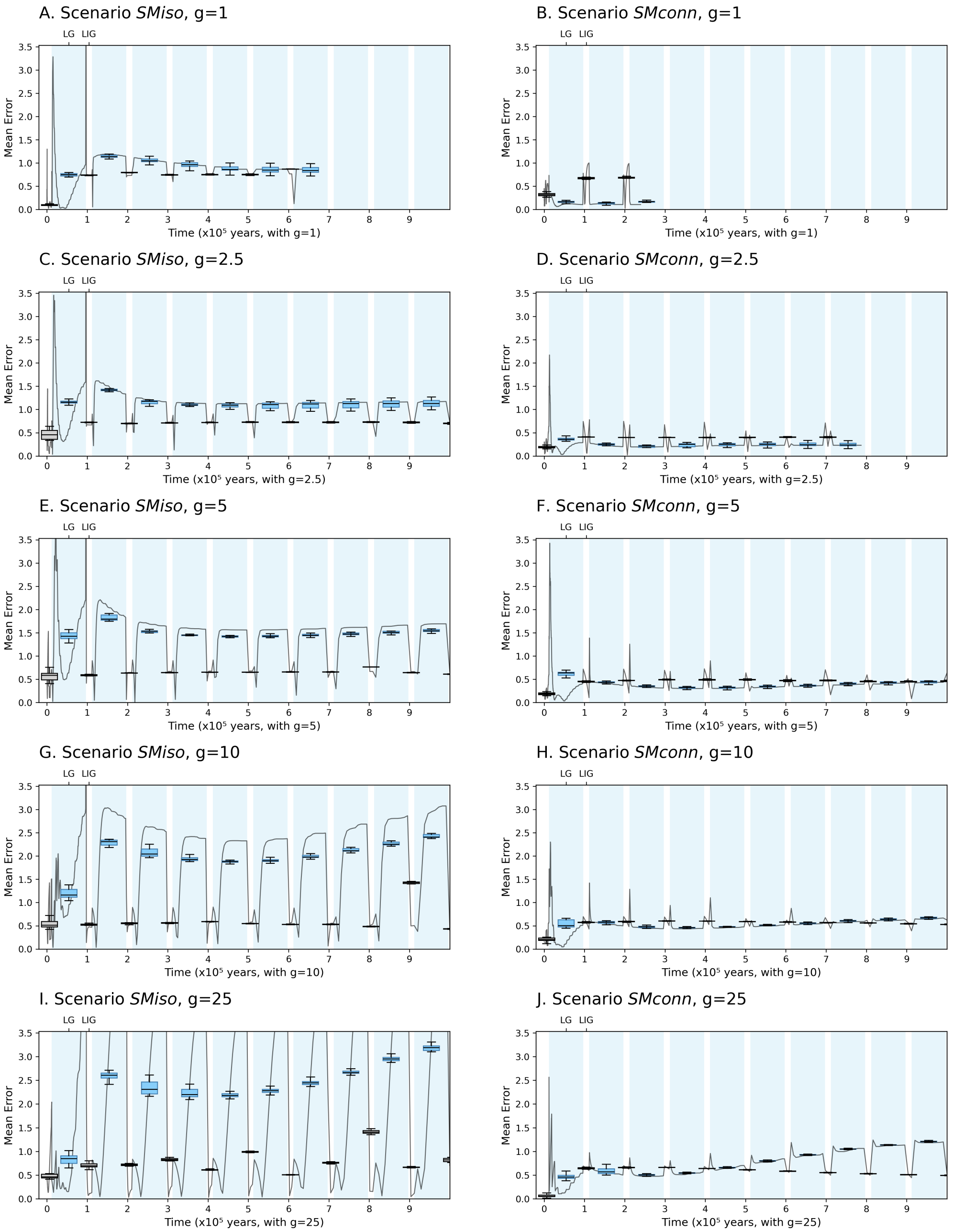


**Figure S24. Mean error (ME) of PSMC estimation of the theoretical IICR across time for scenarios *SM* with $m$ varying between $2.5.10^{-5}$ and $2.5.10^{-3}$ and $N$ 5-fold increasing or decreasing during glacials for several generation times.** A: *SMiso* for g = 1 year, B: *SMconn* for g = 1 year, C: *SMiso* for g = 2.5 years, D: *SMconn* for g = 2.5 years, E: *SMiso* for g = 5 years, F: *SMconn* for g = 5 years, G: *SMiso* for g = 10 years, H: *SMconn* for g = 10 years, I: *SMiso* for g = 25 years, and J: *SMconn* for g = 25 years. The RE was defined as $\frac{|IICR_t - PSMC_t|}{IICR_t}$ and its mean was computed across the three PSMC curves and 15 bootstraps. In each sub-figure, the box-plots correspond to a mean RE computed every 99,9 years (10,000 time points) and averaged for each interglacial and interglacial period (1-12 kya, 12-97 kya, 97-112 kya and so on, up to 1 Mya). The grey curve corresponds to a mean RE computed every 999 years (1,000 time point in total between 1 kya and 1 Mya). Light blue rectangles in the background correspond to glacial periods.

Please note that, due to the cyclic nature of our scenarios and the oscillations of the IICR, the mean RE across time is not necessarily informative about the ability of PSMC to infer the demographic oscillations (see Material and Methods and Supp. Information S1).

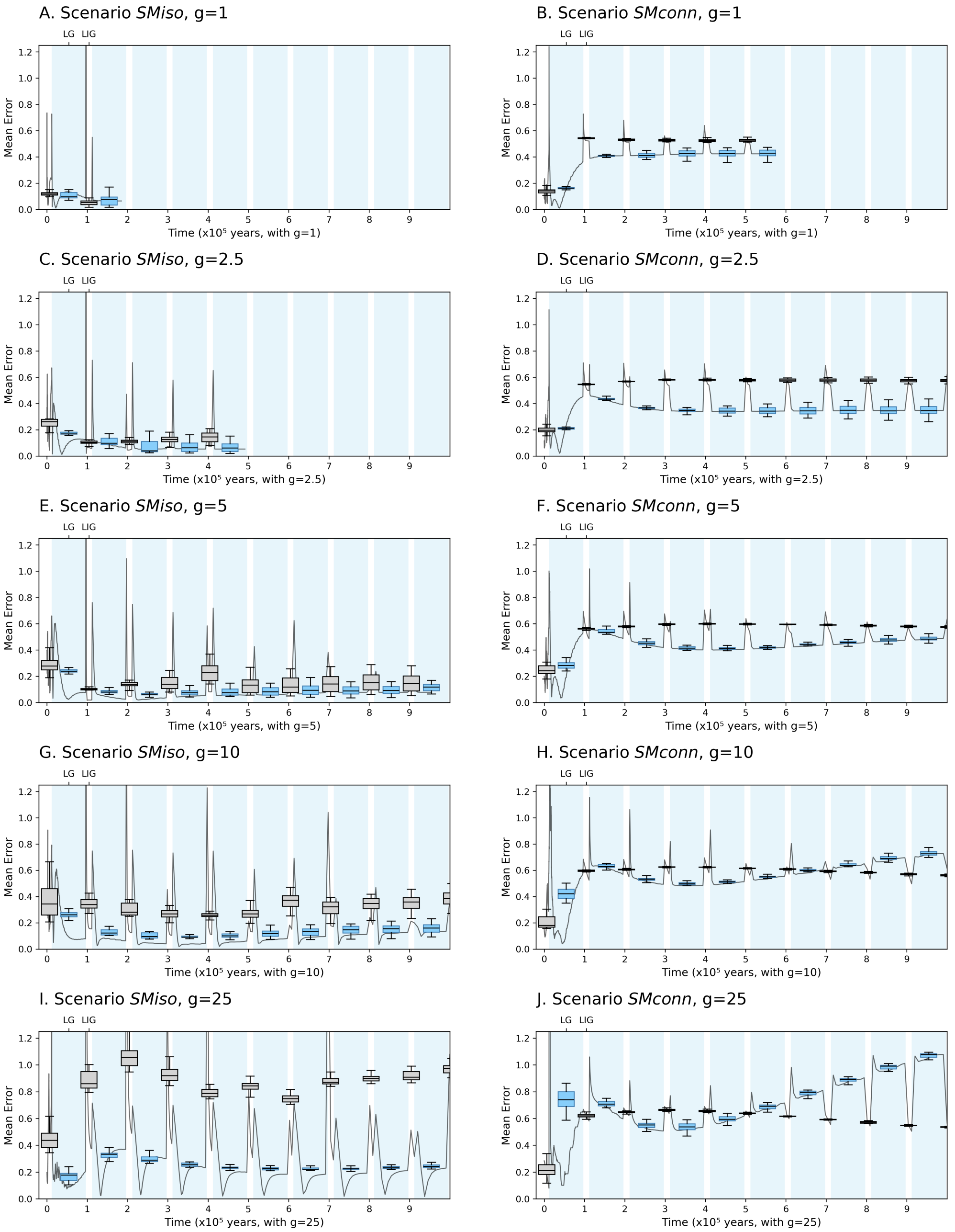


**Figure S25.** **Mean error (ME) of PSMC estimation of the theoretical IICR across time for scenarios *SM* with $m$ varying between $2.5.10^{-4}$ and $2.5.10^{-3}$ and $N$ 5-fold increasing or decreasing during glacials for several generation times.** A: *SMiso* for g = 1 year, B: *SMconn* for g = 1 year, C: *SMiso* for g = 2.5 years, D: *SMconn* for g = 2.5 years, E: *SMiso* for g = 5 years, F: *SMconn* for g = 5 years, G: *SMiso* for g = 10 years, H: *SMconn* for g = 10 years, I: *SMiso* for g = 25 years, and J: *SMconn* for g = 25 years. The RE was defined as $\frac{|IICR_t - PSMC_t|}{IICR_t}$ and its mean was computed across the three PSMC curves and 15 bootstraps. In each sub-figure, the box-plots correspond to a mean RE computed every 99,9 years (10,000 time points) and averaged for each interglacial and interglacial period (1-12 kya, 12-97 kya, 97-112 kya and so on, up to 1 Mya). The grey curve corresponds to a mean RE computed every 999 years (1,000 time point in total between 1 kya and 1 Mya). Light blue rectangles in the background correspond to glacial periods.

Please note that, due to the cyclic nature of our scenarios and the oscillations of the IICR, the mean RE across time is not necessarily informative about the ability of PSMC to infer the demographic oscillations (see Material and Methods and Supp. Information S1).

#### SNIF analyses

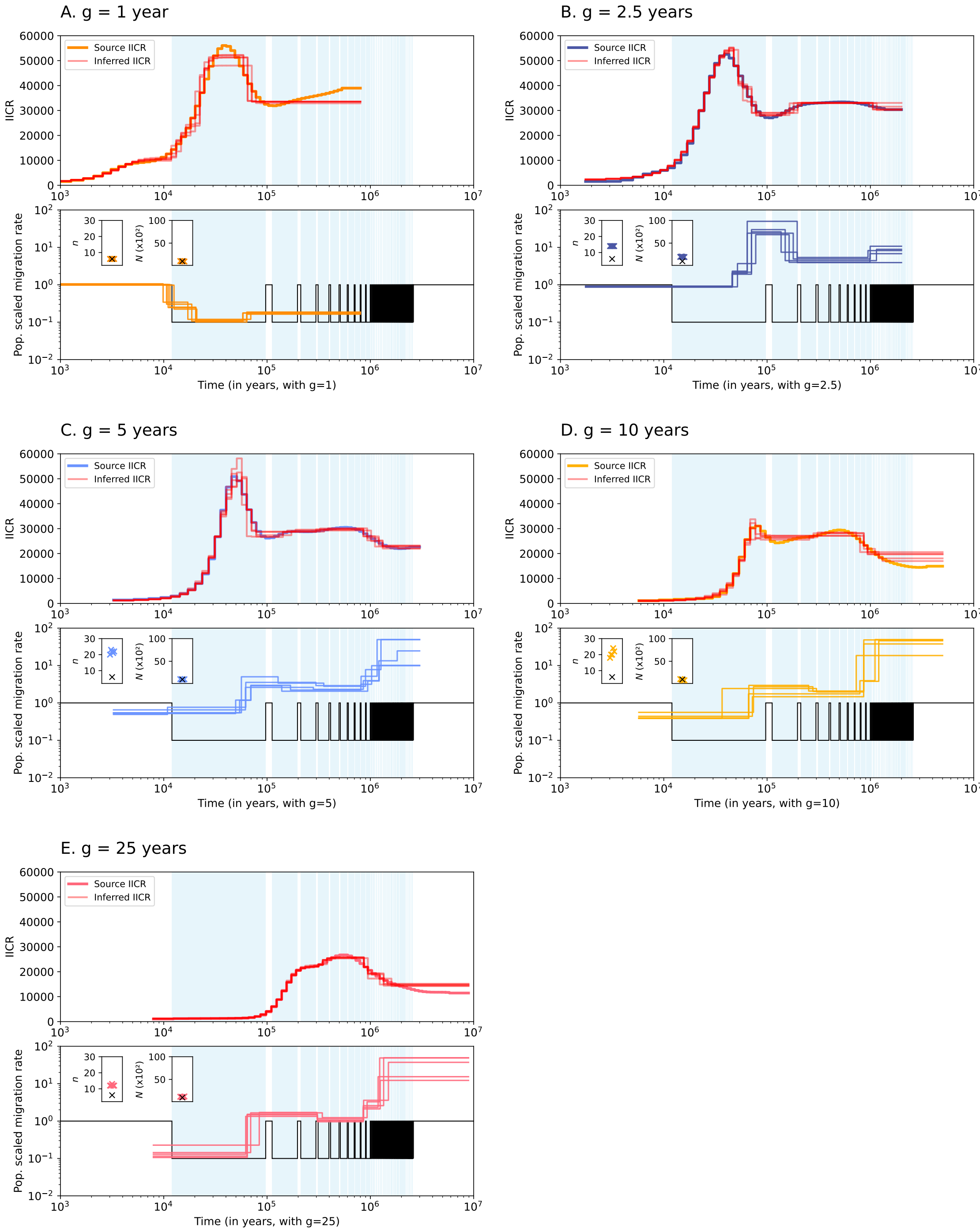


**Figure S26. SNIF-inferred IICR, deme size $N$, number of demes $n$ and (population scaled) migration rates $M_i$ over time for scenarios *Miso* and different generation times, using PSMC curves as input data.** A. 1 year, B. 2.5 years, C. 5 years, D. 10 years and E. 25 years. Top panels show the input PSMC curve and the inferred IICR curves (in red) for five repetitions of SNIF. Bottom panels show the inferred population scaled migration rates ($M_i = 4Nm_i$) and the target (simulated) migration rates, in coloured and black lines, respectively. The embedded plots in the bottom panels show the inferred numbers of demes $n$ (left) and the inferred diploid deme sizes $N$ (right). The coloured and black crosses correspond to the inferred (for five repetitions of SNIF) and the target (simulated) value, respectively.

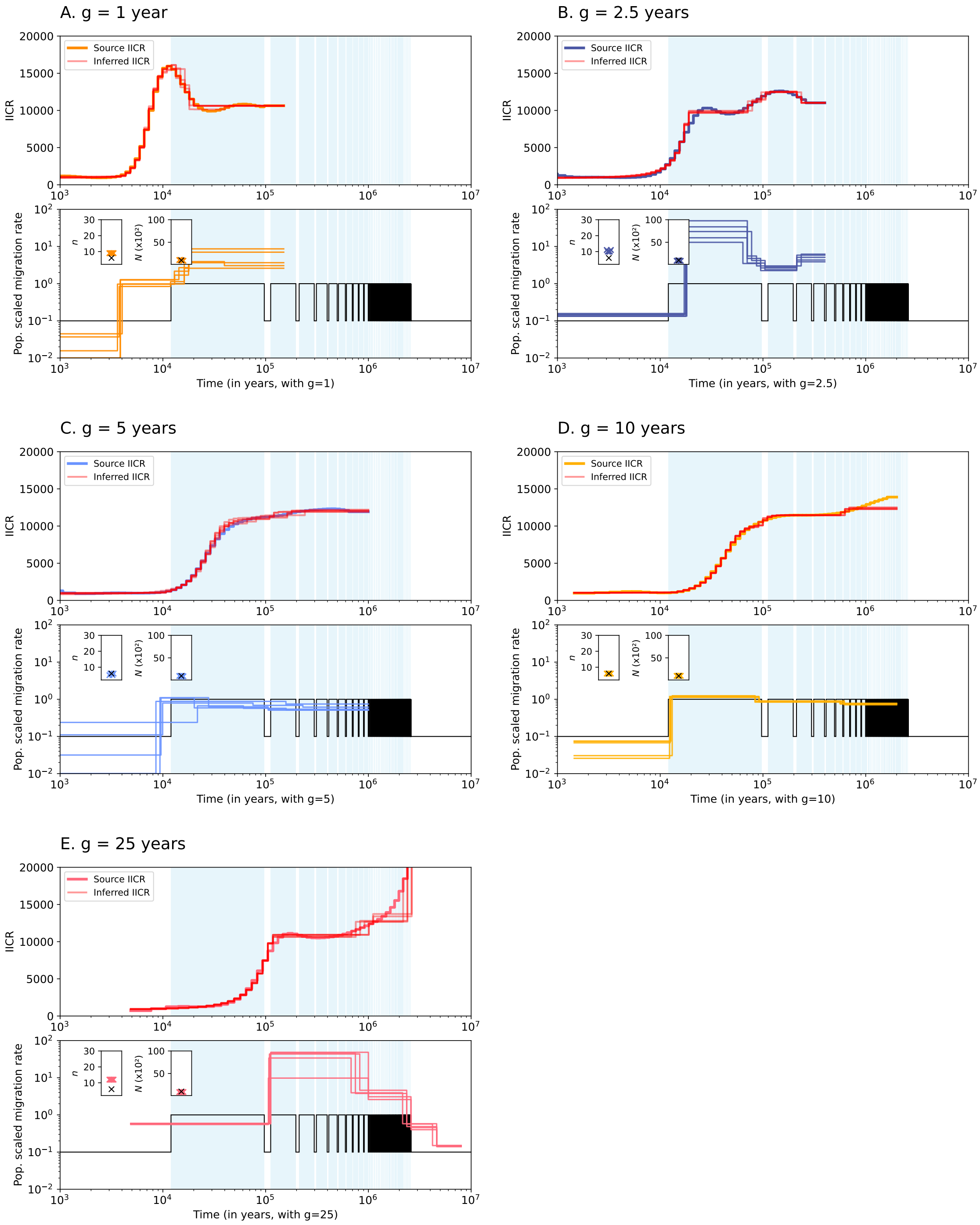


**Figure S27. SNIF-inferred IICR, deme size $N$, number of demes $n$ and (population scaled) migration rates $M_i$ over time for scenarios *Mcon* and different generation times, using PSMC curves as input data.** A. 1 year, B. 2.5 years, C. 5 years, D. 10 years and E. 25 years. Top panels show the input PSMC curve and the inferred IICR curves (in red) for five repetitions of SNIF. Bottom panels show the inferred population scaled migration rates ($M_i = 4Nm_i$) and the target (simulated) migration rates, in coloured and black lines, respectively. The embedded plots in the bottom panels show the inferred numbers of demes $n$ (left) and the inferred diploid deme sizes $N$ (right). The coloured and black crosses correspond to the inferred (for five repetitions of SNIF) and the target (simulated) value, respectively.

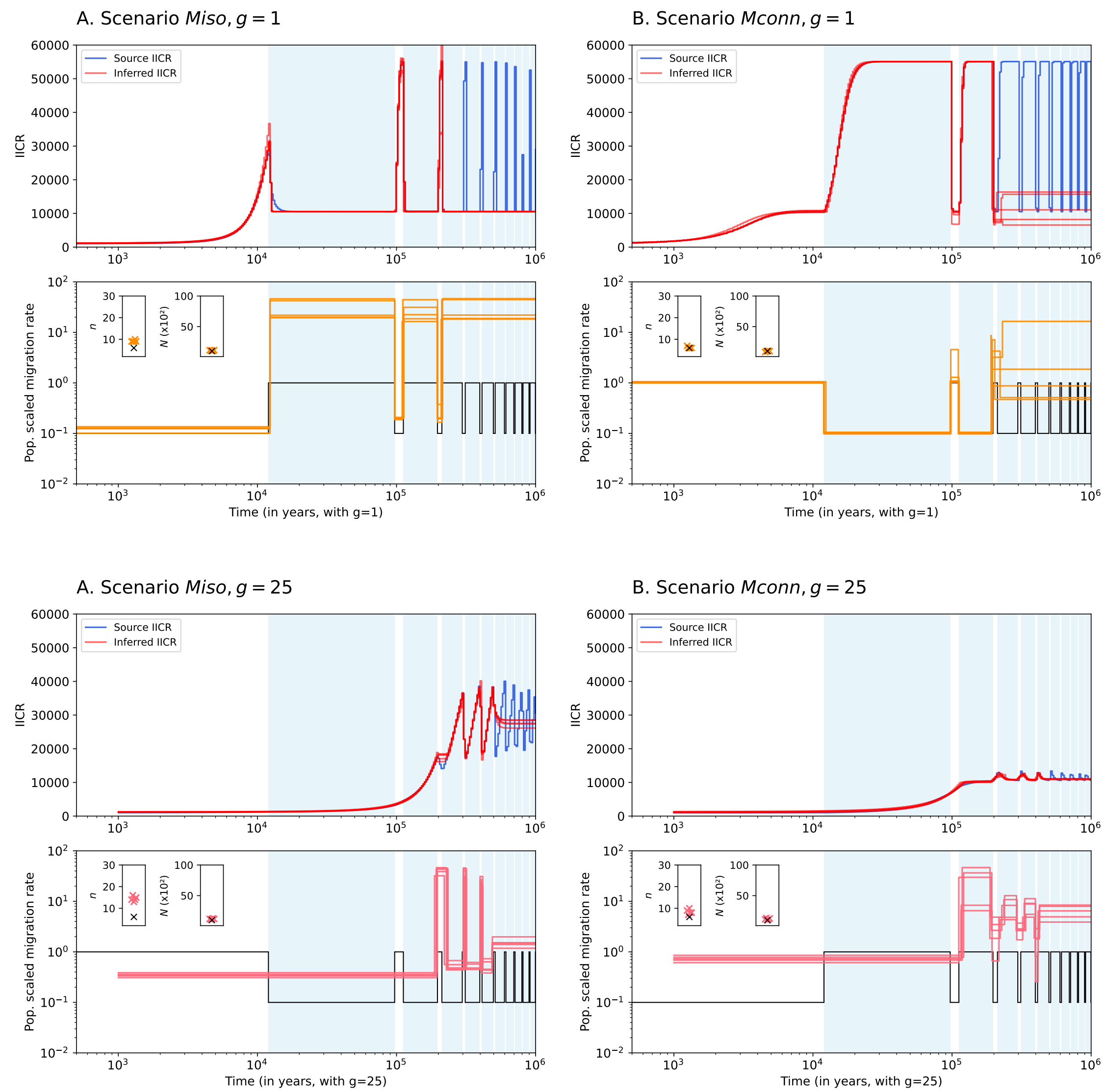


**Figure S28.** **SNIF-inferred IICR, deme size $N$, number of demes $n$ and (population scaled) migration rates $M_i$ over time for scenarios *Mcon* and different generation times, using the theoretical IICR as input data.** A. 1 year, B. 2.5 years, C. 5 years, D. 10 years and E. 25 years. Top panels show the input PSMC curve and the inferred IICR curves (in red) for five repetitions of SNIF. Bottom panels show the inferred population scaled migration rates ($M_i = 4Nm_i$) and the target (simulated) migration rates, in coloured and black lines, respectively. The embedded plots in the bottom panels show the inferred numbers of demes $n$ (left) and the inferred diploid deme sizes $N$ (right). The coloured and black crosses correspond to the inferred (for five repetitions of SNIF) and the target (simulated) value, respectively.

### Inferences across generation times

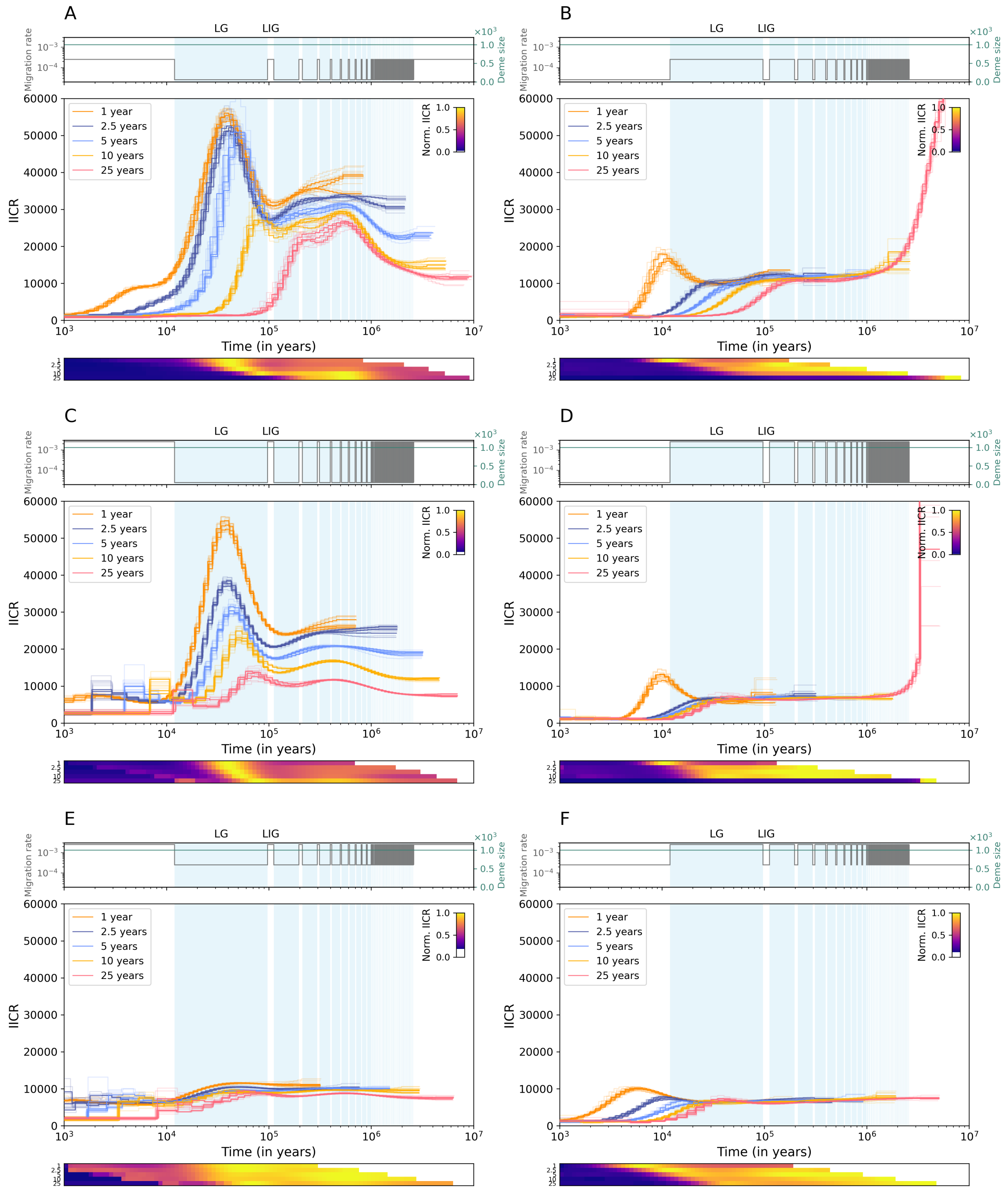


**Figure S29.** **PSMC curves for scenarios $M$ for different combinations of migration rates.** A. $m_{IG} = 2.5.10^{-4}$ and $m_G = 2.5.10^{-5}$, B. $m_{IG} = 2.5.10^{-5}$ and $m_G = 2.5.10^{-4}$, C. $m_{IG} = 2.5.10^{-3}$ and $m_G = 2.5.10^{-5}$, D. $m_{IG} = 2.5.10^{-5}$ and $m_G = 2.5.10^{-3}$, E. $m_{IG} = 2.5.10^{-2}$ and $m_G = 2.5.10^{-3}$ and E. $m_{IG} = 2.5.10^{-3}$ and $m_G = 2.5.10^{-2}$. In each top panel are represented the migration rates and the deme sizes over time, with grey and turquoise lines, respectively. In each middle panel, the coloured lines correspond to the PSMC curves coloured by generation time: 1 (orange), 2.5 (dark blue), 5 (light blue), 10 (yellow) and 25 (pink) years. The lighter coloured curves correspond to the bootstraps (5 per PSMC curve). Each bottom panel represents the normalized IICR over 64 time bins.

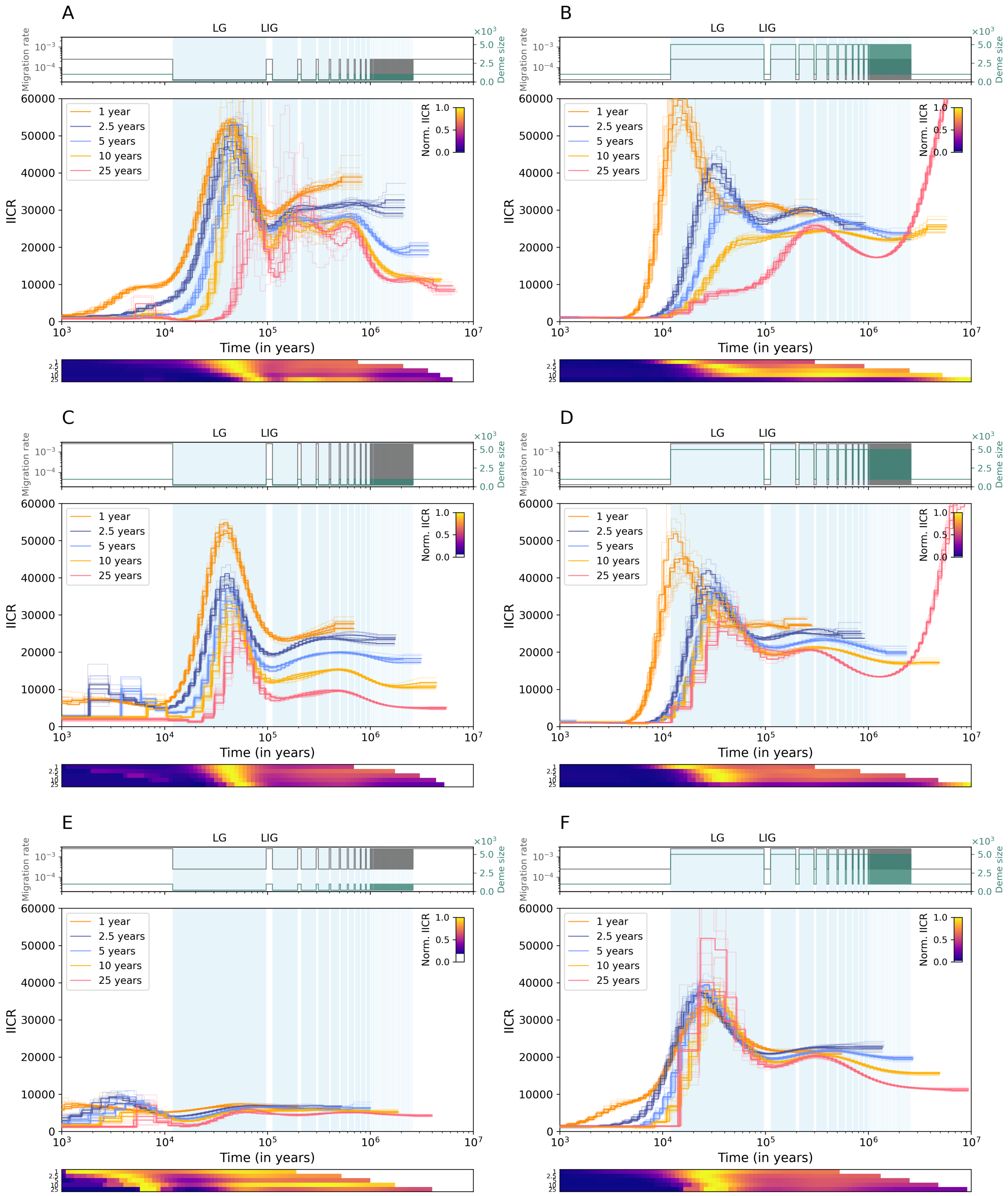


**Figure S30.** **PSMC curves for scenarios *SM* for different combinations of migration rates.** A. $m_{IG} = 2.5.10^{-4}$, $N_{IG} = 1000$ and $m_G = 2.5.10^{-5}$, $N_G = 200$, B. $m_{IG} = 2.5.10^{-5}$, $N_{IG} = 1000$ and $m_G = 2.5.10^{-4}$, $N_G = 5000$, C. $m_{IG} = 2.5.10^{-3}$, $N_{IG} = 1000$ and $m_G = 2.5.10^{-5}$, $N_G = 200$, D. $m_{IG} = 2.5.10^{-5}$, $N_{IG} = 1000$ and $m_G = 2.5.10^{-3}$, $N_G = 5000$, E. $m_{IG} = 2.5.10^{-2}$, $N_{IG} = 1000$ and $m_G = 2.5.10^{-3}$, $N_G = 200$ and E. $m_{IG} = 2.5.10^{-3}$,$N_{IG} = 1000$ and $m_G = 2.5.10^{-2}$, $N_G = 5000$. In each top panel are represented the migration rates and the deme sizes over time, with grey and turquoise lines, respectively. In each middle panel, the coloured curves correspond to the PSMC curves coloured by generation time: 1 (orange), 2.5 (dark blue), 5 (light blue), 10 (yellow) and 25 (pink) years. The lighter coloured curves correspond to the bootstraps (5 per PSMC curve). Each bottom panel represents the normalized IICR over 64 time bins.

#### Stairway Plot analyses

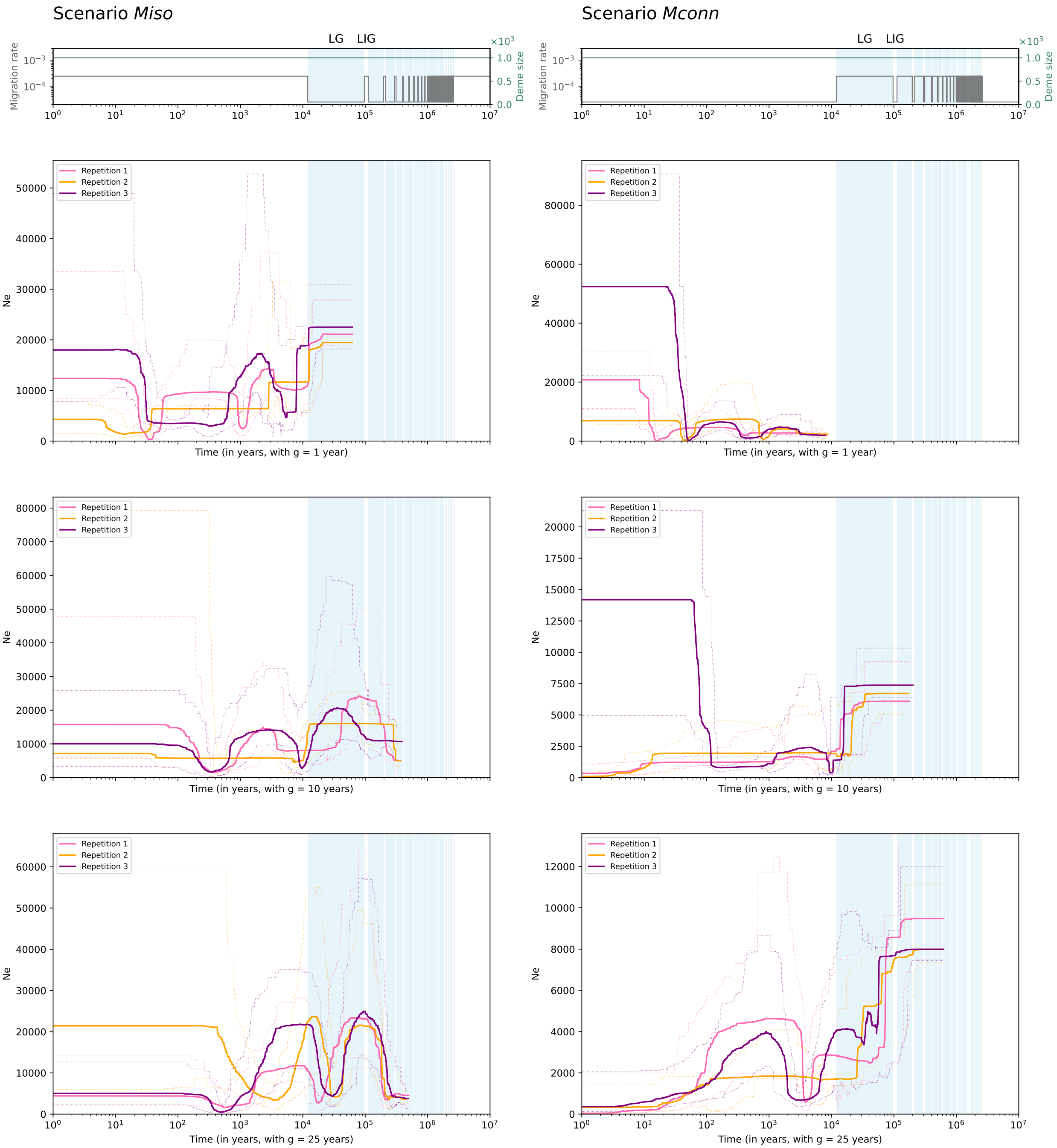


**Figure S31.** Inferences using Stairway Plot 2 on simulated folded SFS for demographic scenarios with changes in migration rates (*Miso* and *Mconn*). Top panels show migration rates and deme sizes over time, represented with grey and turquoise lines, respectively. In the panels below, coloured lines correspond to the median Stairway Plot trajectory and the lighter intervals correspond to the 95% confidence intervals, for a generation of 1, 10 and 25 years for row 2, 3, and 4 respectively. Left column corresponds to *Miso* and right column corresponds to *Mconn*. In these simulated scenarios, migration rates oscillates between $2.5.10^{-5}$ and $2.5.10^{-4}$, and deme size in constant with $N = 1000$. The pale blue areas in the background correspond to glacial periods.

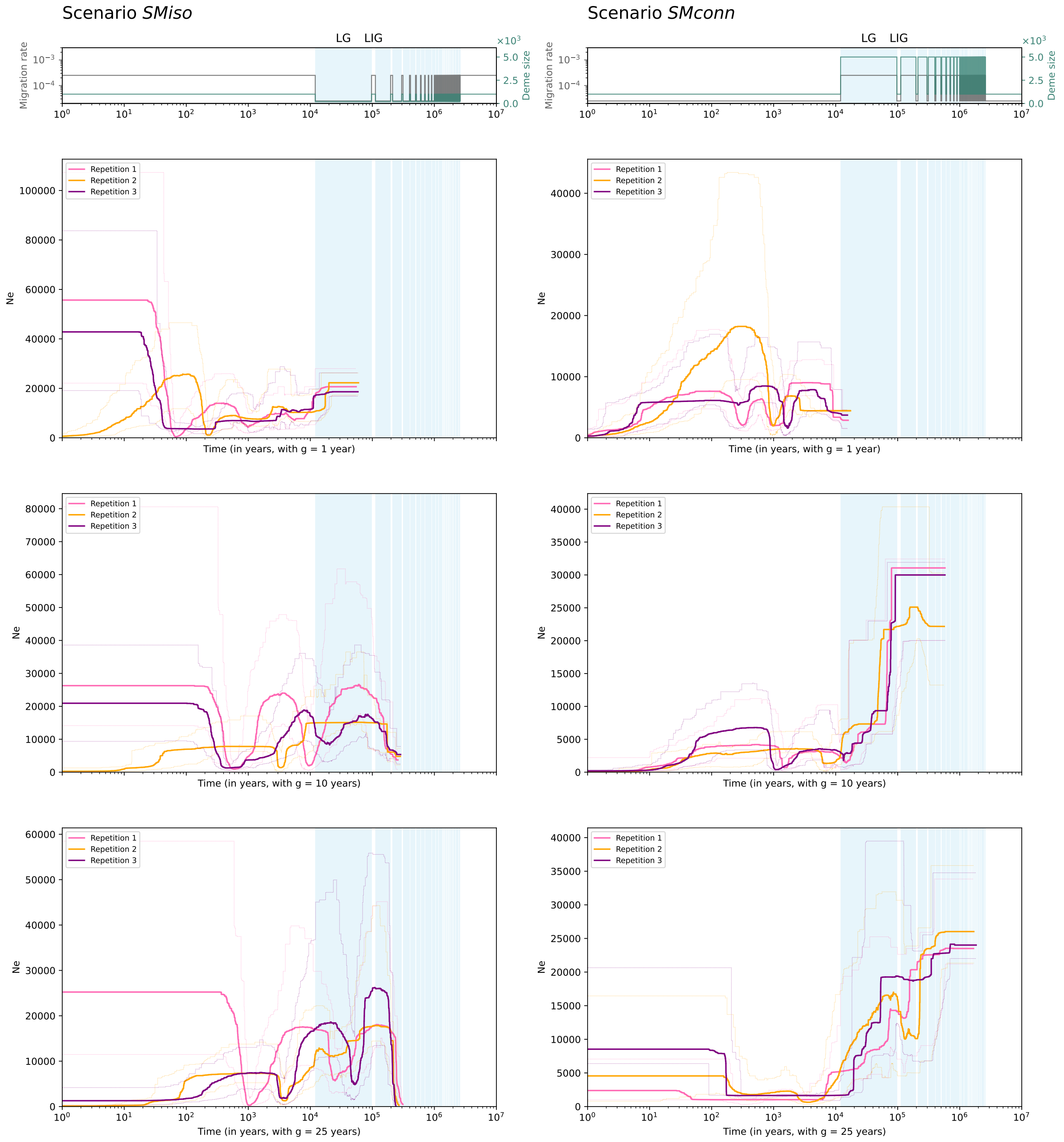


**Figure S32.** Inferences using Stairway Plot 2 on simulated folded SFS for demographic scenarios with changes in migration rates (*SMiso* and *SMconn*). Top panels show migration rates and deme sizes over time, represented with grey and turquoise lines, respectively. In the panels below, coloured lines correspond to the median Stairway Plot trajectory and the lighter intervals correspond to the 95% confidence intervals, for a generation of 1, 10 and 25 years for row 2, 3, and 4 respectively. Left column corresponds to scenario *SMiso*, with $m_{IG} = 2.5.10^{-5}$, $N_{IG} = 1000$ and $m_G = 2.5.10^{-4}$, $N_G = 200$, and right column corresponds to scenario *SMconn* with $m_{IG} = 2.5.10^{-5}$, $N_{IG} = 1000$ and $m_G = 2.5.10^{-4}$, $N_{IG} = 5000$. The pale blue areas in the background correspond to glacial periods.

### Heterozygosity over time

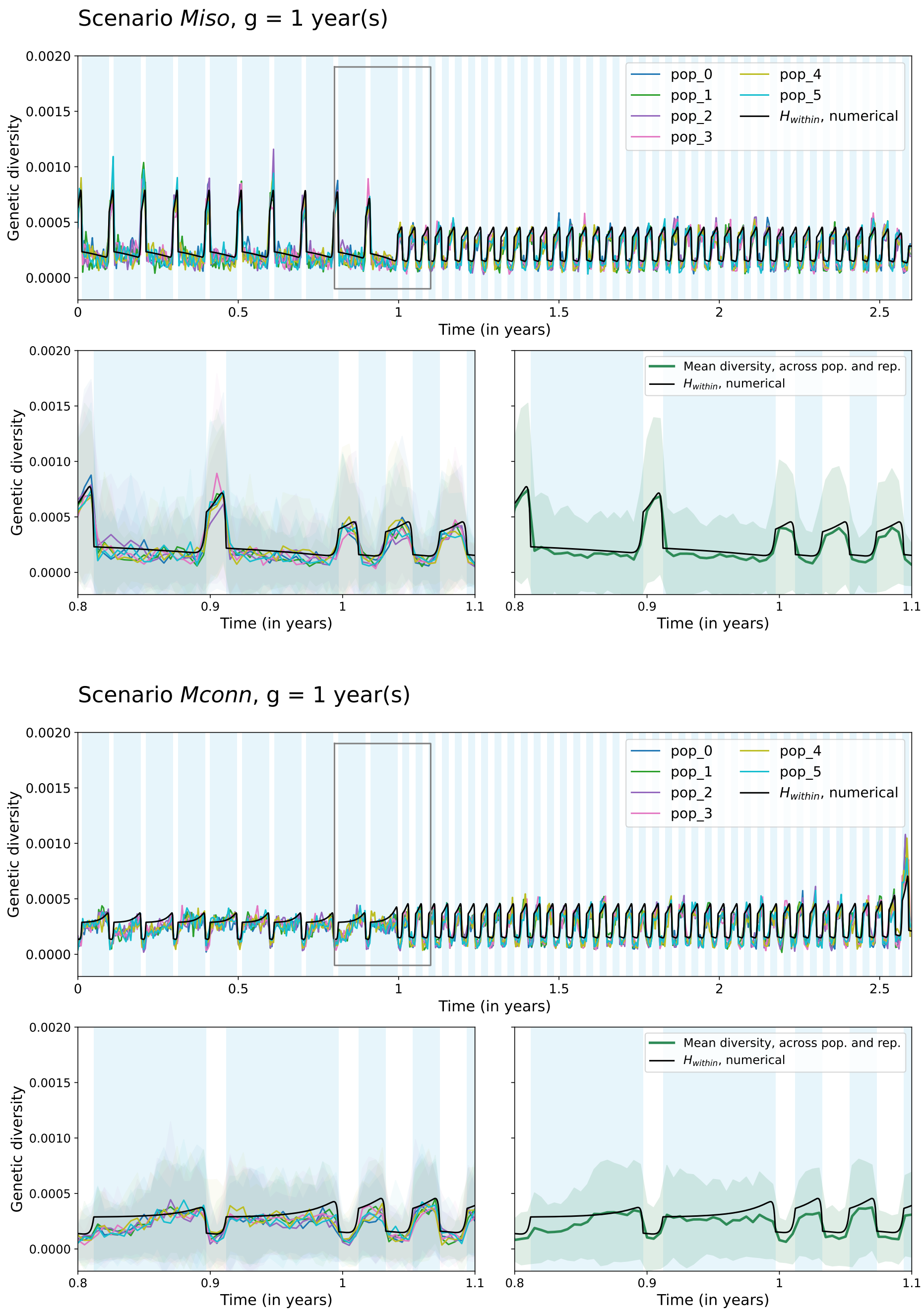


**Figure S33. Numerical and simulated within-deme genetic diversity over time for scenarios *Miso* and *Mconn* and a generation time of 1 year.** A. Scenario *Miso*, B. Scenario *Mconn*. In each sub-figure and panel, the black line represents the (expected) heterozygosity within deme computed numerically following Alcala and Vuilleumier (2014) and Vishwakarma et al. (2026). In all panels, the coloured lines represent the mean genetic diversity within deme computed on simulated genetic data using msprime. In the bottom left panels, the plain coloured lines represent the mean genetic diversity for each deme, and the lighter coloured areas around them show the variance. In the bottom right panels, the green line shows the mean genetic diversity across demes, and the lighter green areas around it shows the variance. In the top panels, the grey rectangle shows the time frame of the two bottom panels. Here, $m$ vary between $2.5.10^{-5}$ and $2.5.10^{-4}$, $N$ is constant and equal to $N = 1000$, and the number of demes is constant and set to $n = 6$. More information regarding this work is available Supplementary Section S3.

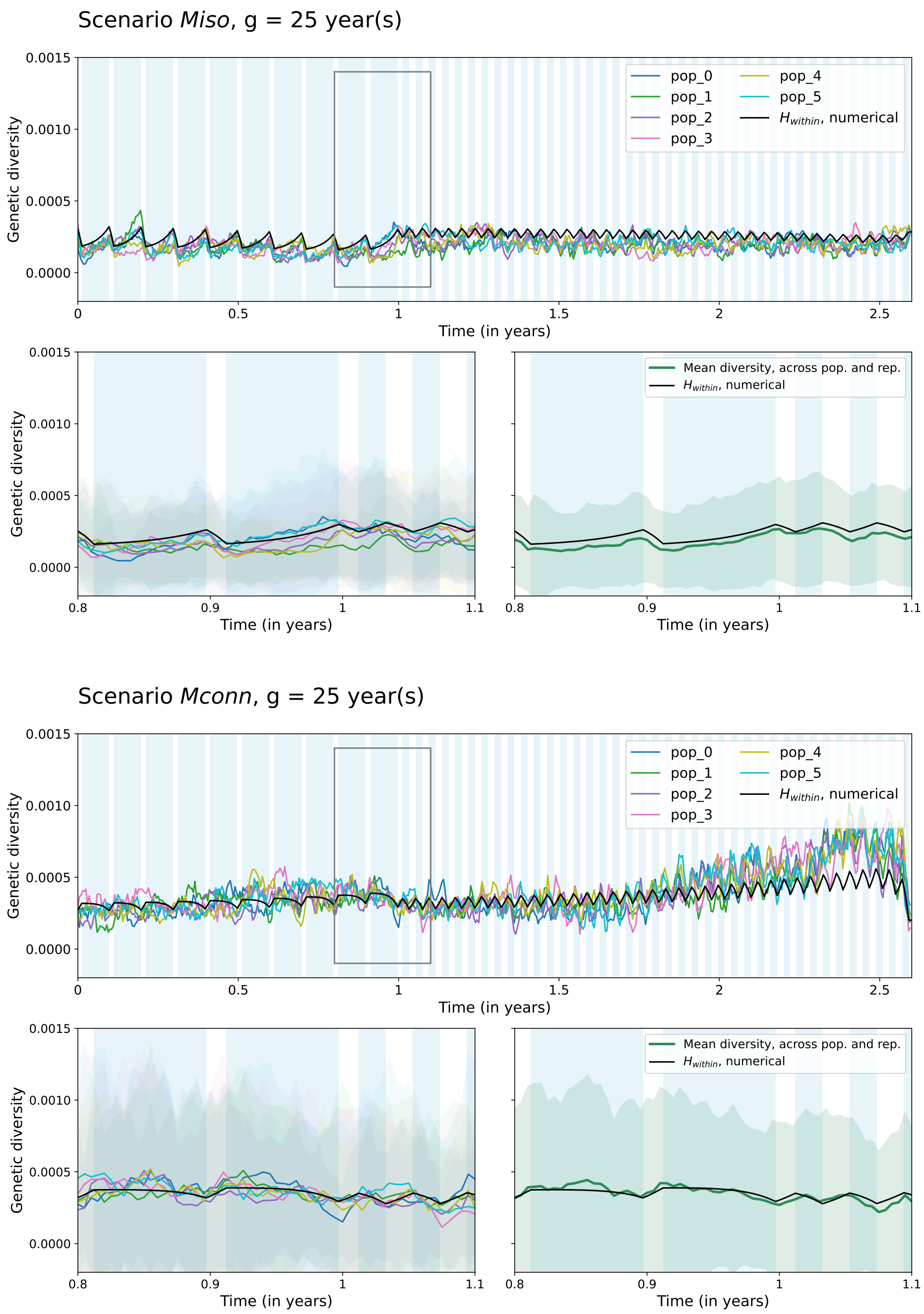


**Figure S34. Numerical and simulated within-deme genetic diversity over time for scenarios *Miso* and *Mconn* and a generation time of 25 years.** A. Scenario *Miso*, B. Scenario *Mconn*. In each sub-figure and panel, the black line represents the (expected) heterozygosity within deme computed numerically following Alcala and Vuilleumier (2014) and Vishwakarma et al. (2026). In all panels, the coloured lines represent the mean genetic diversity within deme computed on simulated genetic data using msprime. In the bottom left panels, the plain coloured lines represent the mean genetic diversity for each deme, and the lighter coloured areas around them show the variance. In the bottom right panels, the green line shows the mean genetic diversity across demes, and the lighter green areas around it shows the variance. In the top panels, the grey rectangle shows the time frame of the two bottom panels. Here, $m$ vary between $2.5.10^{-5}$ and $2.5.10^{-4}$, $N$ is constant and equal to $N = 1000$, and the number of demes is constant and set to $n = 6$. More information regarding this work is available Supplementary Section S3.

#### All PSMC curves

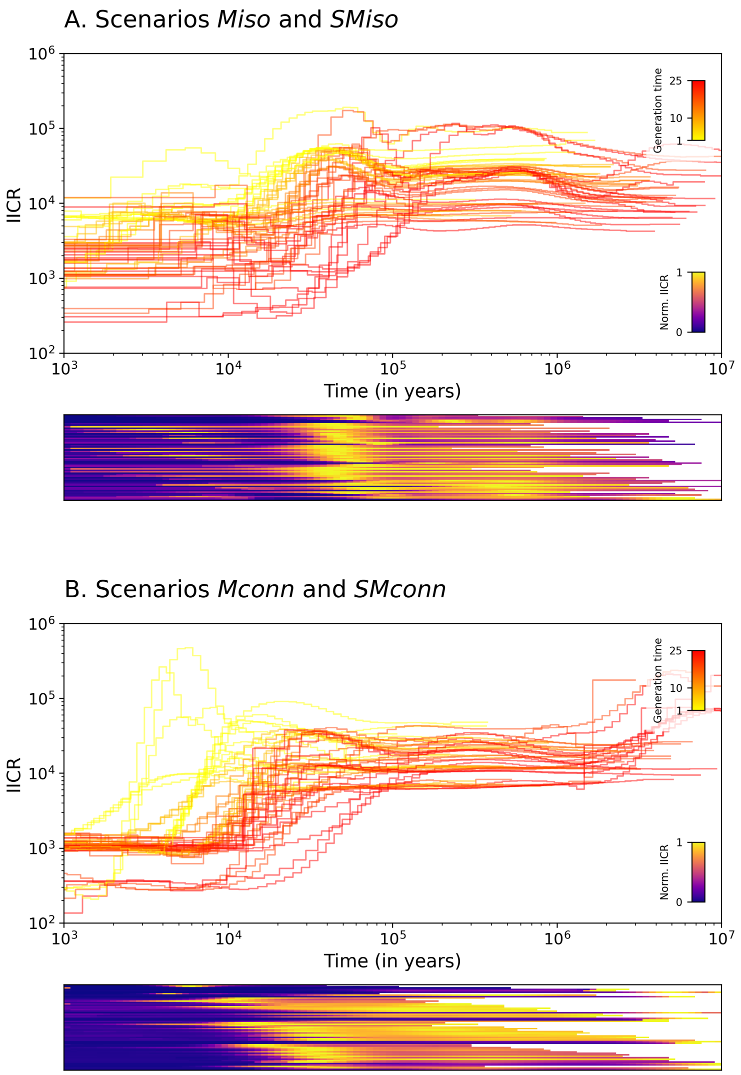


**Figure S35.** **PSMC curves of all the scenarios tested for this study.** A. Scenarios where glacial periods have: (i) lower migration rates or (ii) lower migration rates and deme sizes (*Miso* and *SMiso*). B. Scenarios where glacial periods have: (i) higher migration rates or (ii) higher migration rates and deme sizes (*Mconn* and *SMconn*). Top panels show the PSMC curves, coloured by generation time. Bottom panels show the normalized IICR computed across 100 time bins. Each row of heatmap corresponds to one PSMC curve plotted in the top panel.
